# Topographic structure and function of locus coeruleus norepinephrine neurons

**DOI:** 10.64898/2026.04.10.717727

**Authors:** Zhixiao Su, Polina Kosillo, Kanghoon Jung, Shuonan Chen, Mathew T. Summers, Alex Piet, Han Hou, Kenta M. Hagihara, Drew Friedmann, Olivia Ho-Shing, Matthew I. Becker, Thomas Chartrand, Peter Grotz, Ella Hilton-VanOsdall, Margaret Lee, Rajvi Javeri, Samantha L. Tuggle, Naveen Ouellette, Holly Myers, Camilo Laiton, Kaelin Wulf, John Rohde, Alessio P. Buccino, Cameron Arshadi, Di Wang, Sharmishtaa Seshamani, Sonya Vasquez, Carolyn M. Eng, Douglas R. Ollerenshaw, Nick Dee, Tamara Casper, Windy Ho, Matthew Jungert, Atlas Jordan, Elliot Phillips, Anish Bhaswanth Chakka, Kamiliam Nasirova, Krista Blake, Audrey McCutcheon, Megan Koch, Maria Camila Vergara, Kimberly A. Smith, Tim Jarsky, Nicholas Lusk, Mara C.P. Rue, Xiaoyin Chen, Joshua H. Siegle, Adam K. Glaser, Brian R. Lee, Karel Svoboda, Yoh Isogai, Jayaram V. Chandrashekar, Jeremiah Y. Cohen

**Affiliations:** Allen Institute for Neural Dynamics, Seattle, Washington, USA; Allen Institute for Brain Science, Seattle, Washington, USA

## Abstract

Norepinephrine (NE) is released throughout most of the central nervous system by neurons in the locus coeruleus (LC). We found a relationship between the morphologies, gene expression, and activity of LC-NE neurons in mice. Axonal projections of individual neurons were extensive but largely confined to subsets of brain regions. Axonal projections and graded gene expression correlated with locations of cell bodies in LC. In a behavioral task requiring ongoing learning from actions, neurons in dorsal LC projecting to the cerebral cortex were excited when mice made a different choice from the previous one and by reward prediction errors, a signal driving learning. Background activity of neurons in ventral LC was higher when mice ignored stimuli indicating potential reward availability. These observations reveal a topographically organized structure and function of a neurotransmitter system and show that it contains learning signals for flexible behavior.

## 1. Introduction

Monoamine neurotransmitters are released by small populations of neurons that project extensively throughout the central nervous system (CNS). The locus coeruleus (LC) in the pons supplies the monoamine norepinephrine (NE, also known as noradrenaline) to most of the CNS (Swanson 1976; Amaral & Sinnamon 1977; Jones & Moore 1977; Foote et al. 1983). NE binds G protein-coupled receptors on neurons and glia, activating intracellular signaling pathways (Hein 2006) and inducing synaptic plasticity (Seol et al. 2007; Tully & Bolshakov 2010). NE release has diverse effects on cellular activity across brain targets (Bergles et al. 1996; Kawaguchi & Shindou 1998; Dembrow et al. 2010). The LC-NE system regulates multiple aspects of behavior, including arousal (Foote et al. 1980; Berridge & Waterhouse 2003), sleep/wake transitions (Aston-Jones & Bloom 1981), stress responses (Valentino & Van Bockstaele 2008; McCall et al. 2015), attention (Aston-Jones et al. 1999; Aston-Jones & Cohen 2005), and learning (Bouret & Sara 2004; Dayan & Yu 2006; Jordan & Keller 2023).

The panoply of functions of LC-NE neurons is often ascribed to their complex axonal projections, which span most of the brain and spinal cord. Although the LC-NE system has often been treated as homogeneous, several observations point to heterogeneity among LC-NE neurons (Poe et al. 2020). LC neurons with different somatic morphologies show biases in their axonal projections (Loughlin et al. 1986). *Ex vivo* (Williams et al. 1984; Chandler et al. 2014; McKinney et al. 2023) and *in vivo* (Totah et al. 2018) electrophysiological recordings show diverse biophysical features in LC neurons. Subpopulations of LC-NE neurons exhibit distinct input-output patterns (Schwarz et al. 2015) and have different functions in different CNS regions (Hirschberg et al. 2017; Uematsu et al. 2017). Heterogeneity among LC-NE neurons thus could underlie their diverse functions.

We asked two questions about the LC-NE system: First, is NE released by different LC neurons in different brain regions? To answer this, we examined the morphologies and gene expression patterns of NE neurons across the LC. The projections of individual neurons largely targeted specific brain regions, correlating with patterns of gene expression in the LC. Second, is NE released in different target regions under different behavioral conditions? To answer this, we measured the activity of LC-NE neurons as mice made choices between two options that yielded probabilistic reward. LC-NE physiological properties and activity that correlated with choices varied along similar spatial dimensions as the morphologies and gene expression. Activity of neurons projecting to the cerebral cortex selectively correlated with reward prediction error (RPE), indicating that NE is released in different brain regions at different times and behavioral conditions.

## 2. Results

### 2.1 LC-NE projections are spatially organized

We begin by defining the boundary of mouse LC in the standardized common coordinate framework (CCFv3) (Q. Wang et al. 2020) by mapping NE neurons in the pons. We labeled nuclei of NE neurons expressing Cre recombinase under the control of dopamine beta hydroxylase (*Dbh*-Cre), the rate-limiting enzyme for NE production. We extracted whole brains, made them optically transparent, and imaged them with selective plane illumination microscopy (SPIM; 34,998 cells, 8 mice; Figures 1a, 1b, and S1). We defined LC in the CCF as a small dorsal region (0.13 mm^3^ in volume) containing approximately two-thirds of all pontine NE neurons (67th density percentile, 1507 ± 55, mean ± SD). Additional NE neurons were found in sub-coeruleus (367 ± 95) (Amaral & Sinnamon 1977).

**Figure 1:**
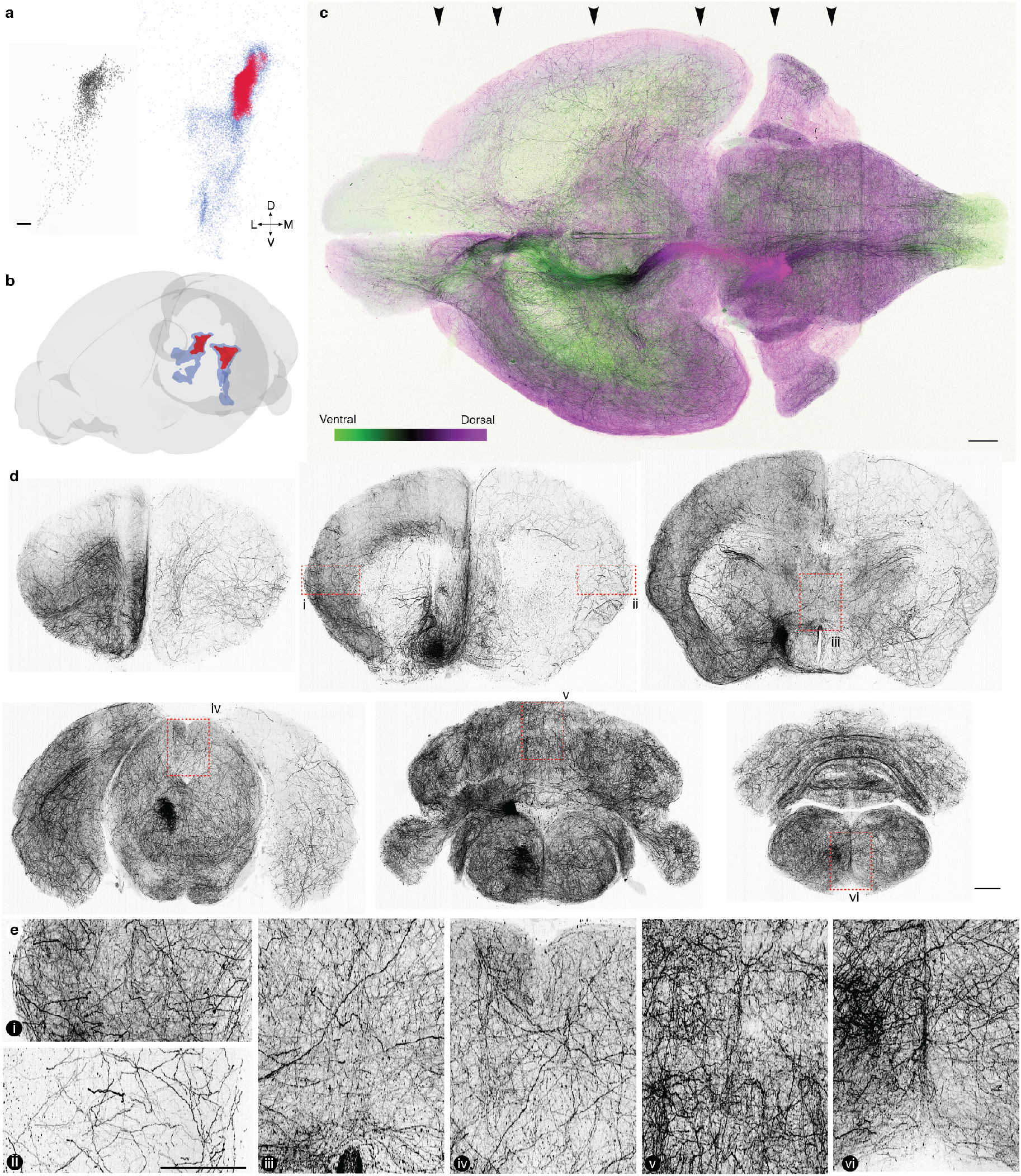
Organization of LC-NE neurons and their brain-wide projections. **a**, Left, coronal view of *Dbh*+ neurons projected across a 500-µm volume. Scale bar, 100 µm. Right, definition of LC-NE neurons in CCF space. Red, region containing 67% of *Dbh*+ cells. **b**, Location of LC in the pons. Blue, region containing 90% of *Dbh*+ cells, including sub-coeruleus. **c**, Maximum intensity projection of LC-NE axons arising from one hemisphere. **d**, Coronal views of axonal innervation across the brain. (maximum intensity projections of 2 mm depth in expanded dimensions). Left hemisphere LC-NE neurons were labeled. Note predominantly ipsilateral innervation of the telencephalon, sparse labeling in the striatum, and bilateral innervation of the thalamus, midbrain, cerebellum, and brainstem. **e**, Detail of labeled axons in regions marked in **d** (scale bar, 2 mm).

We next explored the structure of LC-NE neuron projections (Swanson & Hartman 1975; Nomura et al. 2014). To quantify the distribution of LC-NE axons across the brain, we densely labeled LC-NE neurons unilaterally in *Dbh*-Cre mice. We expanded whole brains and imaged them using a large field-of-view SPIM microscope (Glaser et al. 2025). We found a high density of LC-NE axons in all CNS regions other than the striatum, with varying densities across regions (Figures 1c–1e and S2). Projections to the telencephalon were predominantly ipsilateral, from somata in the same hemisphere. Projections to other regions were both contralateral and ipsilateral (Jones & Moore 1977).

What is the spatial organization of individual LC-NE neuron projections? We reconstructed the complete morphologies of LC-NE neurons (130 cells, 4 mice). We found cells with projections to all regions (defined at level 2 of the CCF ontology) covered by the population (Figures 2 and S3). However, individual LC-NE neurons tended to have axons in a particular region: cortex, thalamus and midbrain, cerebellum, brainstem, or spinal cord (Figure 2b). Many neurons had long axons, 35.14 ± 13.60 cm (mean ± SD; Figure S4a) with the largest neurons reaching 72.57 cm of total length (the longest reported mouse neuron, to our knowledge). Despite having very long axons, these neurons branched little (Figure S4a; 17.8 branches/cm, *R*^2^ = 0.74, *p* < 0.001). Compared to 1,227 other CNS neurons from the MouseLight database (Winnubst et al. 2019) (24.9 branches/cm, *R*^2^ = 0.42, *p* < 0.001), LC-NE neurons branched significantly less (ANCOVA, *p* < 0.001). To quantify the degree to which individual neurons projected to multiple regions, we measured the fraction of each neuron’s axons in each area (Figures 2c and S4b). Most neurons had primary targets, either as part of ascending projections to the forebrain or descending projections to the brainstem and spinal cord. Consistent with largely nonoverlapping pathways, neurons with forebrain projections tended to project to other regions in the forebrain but not the brainstem or spinal cord (Figure 2c). Likewise, neurons with projections to the brainstem or spinal cord tended to co-innervate regions in these descending targets while avoiding the forebrain. Other neurons projected mainly to the cerebellum. We compared these data against multiple models of how axons could fill CNS space. A simple model that assumes axons grow as a random walk with a renewal process that depends on the length of the axons captured some fundamental structural features of LC-NE projections (Figure S4c). However, LC-NE morphologies are likely driven by multiple factors, including developmental programs (Robertson et al. 2013) and adult plasticity (Fritschy & Grzanna 1992).

**Figure 2:**
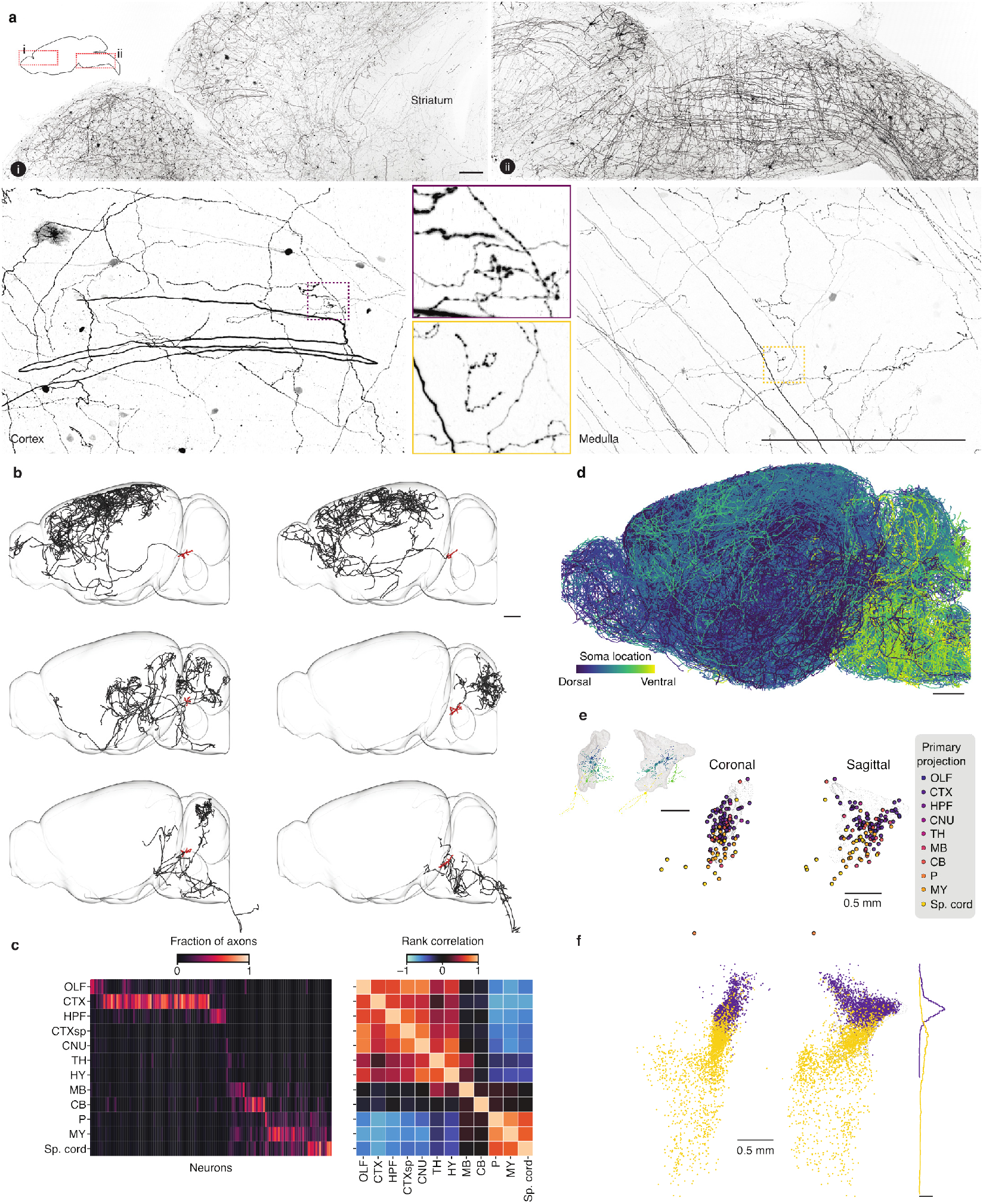
LC-NE projections are spatially organized. **a**, Examples of LC-NE axons labeled in frontal cortex (i) or medulla (ii). Scale bars, 1 mm. **b**, Example morphologies of six fully reconstructed LC-NE neurons. Red, cell bodies and dendrites. Black, axons. Scale bar, 1 mm. **c**, Left, fraction of each neuron’s axons in each region. OLF, olfactory bulb. CTX, isocortex. HPF, hippocampal formation. CTXsp, cortical subplate. CNU, cerebral nuclei. TH, thalamus. HY, hypothalamus. MB, midbrain. CB, cerebellum. P, pons. MY, medulla. Right, rank correlation among neurons projecting to pairs of regions. **d**, Lateral view of single neurons superimposed, colored by locations of somata from dorsal to ventral. Scale bar, 1 mm. **e**, Locations of somata from reconstructed neurons with primary projections to different regions, projected into CCF coordinates in the coronal and sagittal planes. Gray meshes correspond to 67% of *Dbh*+ cells from Figure 1a. Inset, dendritic arbors from the neurons in **b. f**, LC-NE neurons retrogradely labeled from ipsilateral cortex (purple) or spinal cord (yellow). Histogram scale bar, density of 2.

Once a primary target was reached, LC-NE axons occupied large volumes without clear evidence of target specificity. We quantified the extent to which neurons innervating specific regions of the isocortex sent axons to other cortical regions. Individual cells tended to project to nearby regions, without evidence of individual cells consistently targeting spatially separated, but functionally related regions (Figures S4d–S4g).

We compared the morphological reconstructions of individual neurons with multiplexed analysis of projections by sequencing (MAPseq) (Kebschull et al. 2016). We labeled LC cells with sequences of RNA “barcodes,” dissected multiple regions of the brain and spinal cord, and sequenced barcodes from those regions. MAPseq allowed us to identify projections of a large number of single LC-NE neurons in individual brains, albeit with much lower spatial resolution (approximately 1 mm). We found LC-NE neurons with projections across the CNS, with distributions of somata and axons consistent with our single-neuron morphologies (Figure S5).

What is the relationship between locations of LC-NE somata, their dendrites, and their axons? We visualized the projections of neurons (Figure 2d) and their dendrites (Figure S6a) in the dorsal-ventral axis of LC. Dendrites extended from somata along the dorsal-ventral axis (Figure S6). Somata in different locations of LC showed distinct projection targets (Figure 2e). Neurons in more dorsal LC had primary projections to the forebrain (olfactory bulb, isocortex, hippocampus), whereas neurons in more ventral LC had primary projections to the hindbrain (pons, medulla, and spinal cord). We confirmed this observation using retrograde tracing. We injected AAVrg-DIO-FLPo in six regions in *Dbh*-Cre:Ai65 mice: olfactory bulb (4 mice), frontal cortex (5 mice), thalamus (7 mice), midbrain (6 mice), cerebellar cortex (6 mice), and spinal cord (7 mice) (Figure S7). Consistent with reconstructed neurons (Figure 2e), somata of NE neurons with projections to cortex were preferentially in dorsal LC and NE neurons with projections to the spinal cord were in ventral LC (Figures 2f and S7). Thus, LC-NE neurons are organized in space, with neurons spanning the dorsal to ventral LC innervating the anterior to posterior CNS.

### 2.2 LC-NE neurons have graded gene expression in space

We next investigated how the spatial organization of LC-NE neurons relates to its gene expression. We dissected LC and performed single-nucleus RNA sequencing (snRNAseq, 398,912 cells, 40 mice; 4,845 *Dbh*+ LC-NE neurons; Figure S8). We applied a coupled autoencoder-based unsupervised mixture model (Marghi et al. 2024) and Gaussian mixture model clustering in principal component (PC) space. A single cluster best explained the data (Figure S8), although gene expression varied smoothly across the LC-NE neuron population. We thus treated LC-NE gene expression as a continuum. We applied PC analysis (PCA; Figures 3a and S8) and projected each cell onto a polynomial curve fitted to the first ten PCs. This axis, which we label as “pseudospace” (range 0-1), represents the variation in population gene expression. Similarly graded gene expression was seen along one axis in the UMAP projection (Figure S8f). Many individual genes showed smoothly varying gene expression along the pseudospace (Figures 3b and S9). These genes included molecules associated with axon guidance and development (e.g., *Epha6, Robo1*), adhesion molecules (e.g., *Fat3, Lrrtm4, Ncam2*), extracellular matrix related proteins (*Chsy3, Col18a1, Col6a5, Hs6st3*), ion channels (e.g., *Hcn1, Scn9a*), neuropeptide and neuromodulator receptors (e.g., *Trhr, Tacr3, Sctr, Olfr78*), and transcriptional regulation (e.g., *Nrip1, Tshz1*). In some cases, two related genes expressed in opposing gradients: for example, *Epha6* versus *Epha7* and *Col18a1* versus *Col6a5*. Thus, variation in LC-NE gene expression is dominated by smooth variation.

**Figure 3:**
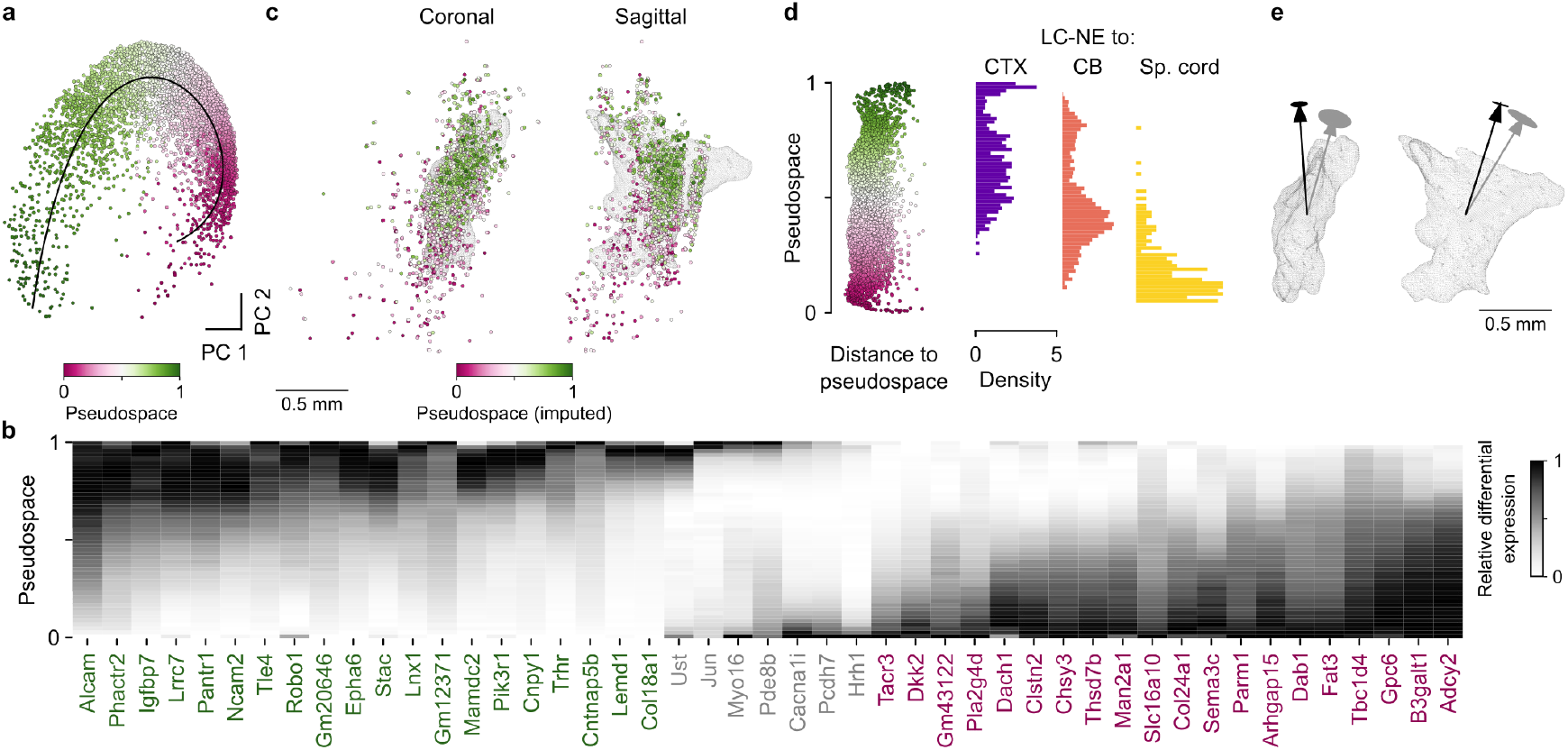
LC-NE gene expression is graded in space and varies by projection target. **a**, First two PCs of batch-normalized scVI counts for 4,845 LC–NE neurons from snRNAseq, colored by “pseudospace.” Pseudospace is defined by projecting each cell onto the polynomial curve fitted (0.47 variance explained) to the first ten PCs (0.80 variance explained). **b**, Heatmap of relative differential expression for a selection of genes along the pseudospace axis. Genes in magenta show highest expression at low pseudospace values. Genes in green peak at high values. Genes in gray are enriched at both high and low pseudospace (but not at intermediate scores). **c**, CCF-aligned mapping of 2,221 MERFISH-labeled LC–NE neurons in the pons (Nardone et al. 2024) in coronal and sagittal planes. Each cell is colored by its imputed location on the pseudospace axis. **d**, Projection target was imputed for each cell in snRNAseq based on the similarity mapping from retroseq data (Kruskal-Wallis, *p* < 0.001). **e**, Primary axes that explain axonal projections (black, from retrograde tracing) and gene expression (gray, from MERFISH) were primarily dorsal-ventral but differed 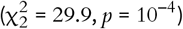. Ellipses: 95% CI.

How does the LC-NE gene expression map onto space in LC? We analyzed spatial gene expression data from multiplexed error-robust fluorescence *in situ* hybridization (MERFISH) in the pons (Nardone et al. 2024). We registered 2,221 LC-NE MERFISH neurons to the CCF and used our snRNAseq data as a reference to impute their pseudospace scores. Gene expression pseudospace aligned with the dorsal-ventral axis of the LC (Figures 3c and S10) and thus with the organization of projection targets observed in our anatomical experiments (Figure 2).

What is the link between spatial variation in projection targets and graded gene expression? We answered this by measuring gene expression in LC-NE neurons retrogradely labeled from neocortex, cerebellum, or spinal cord (“retro-seq”). We embedded the gene expression from cells in each group into the low-dimensional space describing the population from snRNAseq (Figure S11). Neurons projecting to each region occupied different parts of gene expression space (Figure 3d). Cortex-projecting neurons shared gene expression with dorsal LC-NE neurons, spinal cord-projecting neurons with ventral neurons, and cerebellum-projecting neurons were in between.

Finally, we calculated the axes that explained most of the variation in axonal projections or gene expression. Both axes aligned with the dorsal-ventral direction, separated by 22.20◦ (Figure 3e). Thus, LC-NE gene expression correlates with cellular morphologies: NE neurons that project to anterior regions of the CNS lie in the dorsal part of LC, NE neurons that project to posterior regions of the CNS lie in the ventral part of LC, and these cells have different patterns of gene expression.

### 2.3 Spatial variation of LC-NE activity during action-outcome learning

Does the spatial variation in LC-NE neuronal structure map onto its activity during behavior? Is NE released at different times in different places in the CNS? LC-NE neurons have been proposed to modulate sensation, action, and learning. Thus, we focused on dynamic action-outcome learning, which includes all three components.

Head-restrained mice were trained on a dynamic reinforcement-learning task (Grossman et al. 2022) (Figures 4a and 4b). Each day mice performed a few hundred trials (mean, 342, range, 94–558 trials; Figure S12). On each trial, following a “go cue,” mice chose freely between licking a leftward or rightward tube. Each tube probabilistically delivered a water reward following a 300-ms delay after the lick. The outcome (presence or absence of reward) was followed by a variable inter-trial interval before the onset of the next trial.

**Figure 4:**
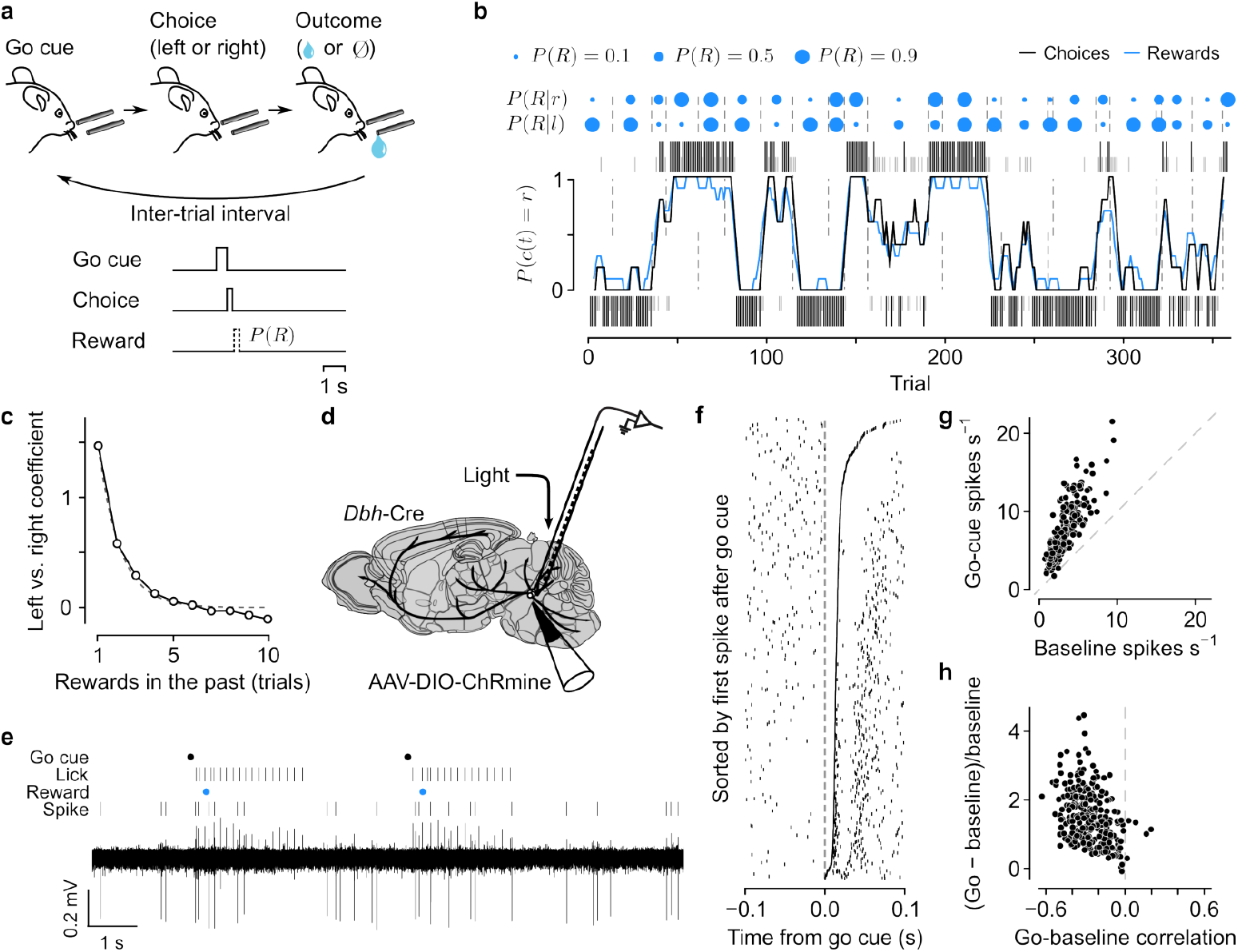
LC-NE neurons are active during action-outcome learning. **a**, Following a go cue, mice chose leftward or rightward licks for probabilistic reward. Reward probabilities (*P*(*R*)) changed over trials. **b**, Example behavior. Tall black (rewarded) and short gray (unrewarded) ticks correspond to left (below) and right (above) choices. Black curve: mouse (smoothed over 5 trials, boxcar filter) choices. Blue curve: rewards (smoothed over 5 trials). Blue dots indicate left/right reward probabilities (*P*(*R*|*l*) and *P*(*R*|*r*), respectively) and dashed lines indicate a change in *P*(*R*). **c**, Logistic regression coefficients for choice as a function of outcome history for all mice (error bars are smaller than points). *R*^2^ = 0.60, successful choice prediction, 0.88. Gray: exponential fit (coefficient of 3.95). **d**, Strategy to record spiking from LC-NE neurons. **e**, Example spiking from an LC-NE neuron around two trials. **f**, Example neuron showing anticorrelation between baseline spike rates and spike rate after the go cue. The raster plot is sorted by the time of the first spike after the go cue. **g**, Spike rates in the 300 ms following go cues relative to the 2 s preceding go cues (241 cells, 19 mice). **h**, Change in go-cue spike rate relative to baseline, normalized by baseline vs. the correlation between go-cue and baseline spike rates (*r* = –0.38, *p* = 8.3 × 10^−11^).

Reward probabilities changed in blocks of trials for leftward and rightward choices, inducing mice to learn from experience. Mice tracked the changing reward probabilities, using reward and choice history to influence future choices (Figure 4c). We compared LC-NE activity to three task events: the go cue (a sensory stimulus), the choice (read out as a directional movement), and a learning signal driven by the presence or absence of reward.

To measure the activity of single LC-NE neurons, we made electrophysiological recordings from LC of *Dbh*-Cre mice. We expressed light-gated ion channels—either channelrhodopsin-2 (ChR2), Chrimson, or ChRmine—selectively in NE neurons. We delivered 473-nm or 640-nm light to the region surrounding the electrodes (Neuropixels probes or tetrodes; Figure 4d). We identified LC-NE neurons as those with short-latency and high-probability responses to light (Figure S13) and estimated their locations in the LC using histological and electrophysiological landmarks. Variation in extracellular action potential waveforms mirrored spatial variation in axonal projections and gene expression (Figure S14): dorsal LC-NE neurons tended to have more symmetric spike waveforms and longer peak-to-trough times than ventral LC-NE neurons.

To test whether extracellular spike waveforms arose from differences in intrinsic biophysical properties of LC-NE neurons, we made whole-cell patch clamp recordings from LC-NE neurons in brain slices (96 cells, 28 mice). We retrogradely labeled neurons from frontal cortex (27 cells, 8 mice), cerebellar cortex (15 cells, 7 mice), or spinal cord (54 cells, 13 mice) (transgenic and viral strategy used for retrograde tracing, Figure 2f ). At the end of the recording, we extracted the nucleus for RNA sequencing (Lee et al. 2021) and confirmed each cell’s identity as an NE neuron. Neurons projecting to spinal cord had significantly different action potential waveforms than those projecting to isocortex or cerebellum. Neurons projecting to isocortex had significantly longer membrane time constants than those projecting to cerebellum or spinal cord (Figure S14). The latter observation may be consistent with the differences in extracellular spike waveforms we observed in spatial coordinates (Pettersen & Einevoll 2008). Thus, intrinsic properties of LC-NE neurons vary along similar spatial dimensions as axonal projections and gene expression.

Nearly all LC-NE neurons showed an increase in spike rate in a 300 ms window following the go cue (paired *t*-tests, 240/241 cells, 19 mice; Figures 4e–4h). Activity during the trial could be due to the go cue, the preparation and execution of the choice movement, or the presence or absence of reward. We analyze each of these below.

On some trials, mice ignored the go cue and did not lick in either direction. A logistic regression predicting whether mice responded to a go cue with a lick revealed a dependence on reward history (Figure 5a). LC-NE neurons had higher spike rates preceding ignored trials relative to those with a lick response (Figures 5b and S15). Across the population of LC-NE neurons, spike rates in the 2 s before the go cue predicted whether the mouse would successfully respond to the go cue with a lick in either direction (46.5% of neurons had *p*-values < 0.05). This effect was strongest in neurons in ventral LC (Figure 5c; *p*-values from permutation test of linear regression between *t*-statistics and spatial location: *R*^2^ = 0.152, 1.4 × 10^−3^). To compare the spatial distribution of activity with our anatomical findings, we projected each neuron onto the axes derived from the retrograde tracing and MERFISH data. We found a significant mapping of cellular activity onto axonal projections and gene expression (Figure 5d).

**Figure 5:**
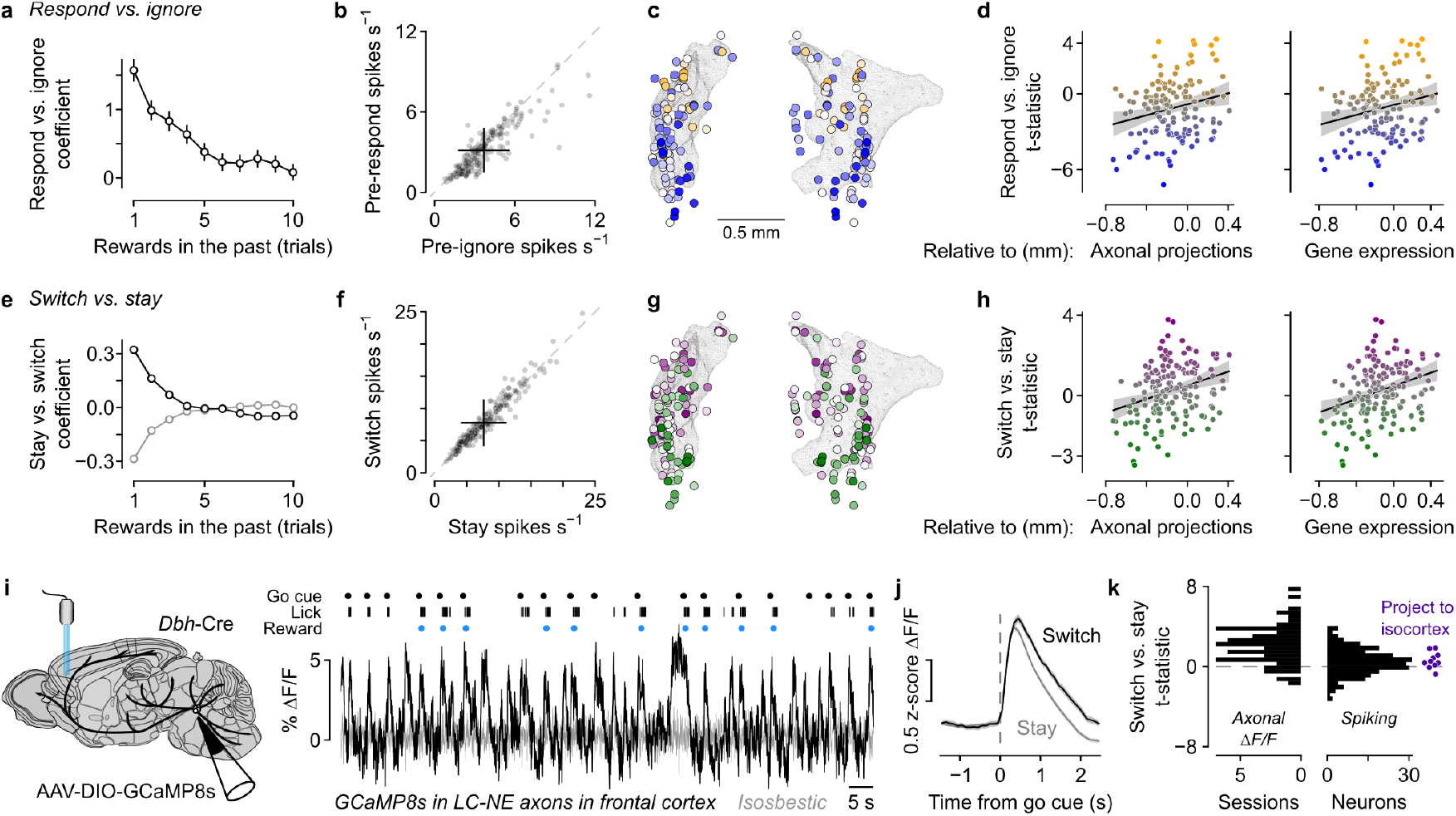
Choice-related LC-NE activity is spatially organized. **a**, Logistic regression coefficients for respond vs. ignore as a function of reward history for all mice. Error bars: 95% CI. **b**, Spike rates in the 2 s before go cues followed by a lick response versus those ignored. Error bars: SD. **c**, Neurons with higher activity preceding ignore vs. respond trials (blue) were located more in ventral than dorsal LC (103 cells, 15 mice; *t*-statistics shown in **d**; kNN cross-validated *R*^2^ = 5.57 × 10^−2^, *p* = 8.4 × 10^−3^; linear model *R*^2^ = 0.152, *p* = 1.4 × 10^−3^). **d**, Respond vs. ignore *t*-statistics projected onto the primary spatial axes from axonal projections and gene expression (Figure 3e; left, *r* = 0.23, *p* = 6.1 × 10^−3^; right, *r* = 0.23, *p* = 5.8 × 10^−3^). **e**, Logistic regression coefficients for switch vs. stay as a function of no-reward (black) and choice (gray) history for all mice (error bars are smaller than points). **f**, Spike rates after go cues during switch vs. stay trials. Error bars: SD. **g**, Neurons with higher activity during switch vs. stay choices (magenta) were located more in dorsal than ventral LC (114 cells, 15 mice; *t*-statistics shown in **h**; kNN cross-validated *R*^2^ = –4.96 × 10^−2^, *p* = 0.15; linear model *R*^2^ = 0.168, *p* = 3.0 × 10^−4^). **h**, Switch vs. stay *t*-statistics projected onto the primary spatial axes from axonal projections and gene expression (left, *r* = 0.32, *p* = 6.1 × 10^−5^; right, *r* = 0.30, *p* = 1.7 × 10^−4^). **i**, Example calcium-dependent ΔF/F of LC-NE axons in frontal cortex. **j**, Mean ± SEM activity from LC-NE axons in frontal cortex during switch and stay trials. **k**, Switch vs. stay *t*-statistics for LC-NE axons in frontal cortex (left), LC-NE spiking (right), and spiking from LC-NE neurons with identified projections to isocortex, using antidromic stimulation and collision tests.

Choices consisted of repeating the same lick direction (“stay”) or making a different one (“switch”). Mice were likely to repeat choices when they received recent rewards from those lick directions (Figure 5e). We measured the responses of LC-NE neurons in a 500-ms window after the go cue and compared switch versus stay choices. Neurons with higher activity during “switch” choices tended to be in dorsal LC, whereas those with higher activity during “stay” choices tended to be in ventral LC (Figures 5f, 5g, S15, and S16; *R*^2^ = 0.168, 3.0 × 10^−4^). Comparing again to the anatomy, we found spatial variation of switch versus stay activity relative to axonal projections and gene expression (Figure 5h).

Given that dorsal LC-NE neurons showed larger responses during switch versus stay choices, and that projections to cortex come from dorsal neurons, we predicted that LC-NE activity in cortex would be higher during switch versus stay choices. We tested this hypothesis by measuring the activity of single LC-NE neurons projecting to the isocortex and the calcium-dependent fluorescence of LC-NE axons innervating frontal cortex. We focused on prelimbic cortex (PL), which has persistent spike rates correlated with relative choice value and is necessary for performance in the present behavioral task (Bari et al. 2019). We recorded spiking from LC-NE neurons with projections to cortex, identified using antidromic stimulation and collision tests (Figure S13). We expressed the calcium sensor jGCaMP8s (Zhang et al. 2023) in LC-NE neurons and recorded fluorescence in PL using an implanted optic fiber (Figures 5i and S17). LC-NE axonal fluorescence changes in PL and single-neuron spiking projecting to frontal cortex were significantly higher during switch versus stay choices (Figures 5j and 5k; Fisher’s exact test, *p* = 7.8 × 10^−5^). We conclude that activity of LC-NE input to frontal cortex, arising primarily from dorsal LC, is higher during switch relative to stay choices.

### 2.4 Activity of LC-NE projections to frontal cortex correlates with reward prediction errors

Do different LC-NE neurons behave differently as mice learn from action outcomes? Across the population, LC-NE neurons showed varied responses to the reward or its absence (76.8% cells with significant regressions; Figures 6a and S15). Theoretical work proposed that RPEs guide decisions by updating the internal models used to generate behavioral policies. We fit reinforcement-learning models to mouse choice behavior (Figure S12) and found that future choices correlated with estimates of RPEs (Figures 6b and 6c), consistent with the observation that mice used reward history to guide choices. Figure 6d shows two examples of LC-NE neuron activity. Following the choice lick, the example neuron on the left showed a monotonic increase in spike rate with increasing RPE. By contrast, the neuron on the right was excited by lack of reward, without modulation by expected reward. Importantly, these responses were evoked without any external stimulus, other than the presence or absence of reward.

**Figure 6:**
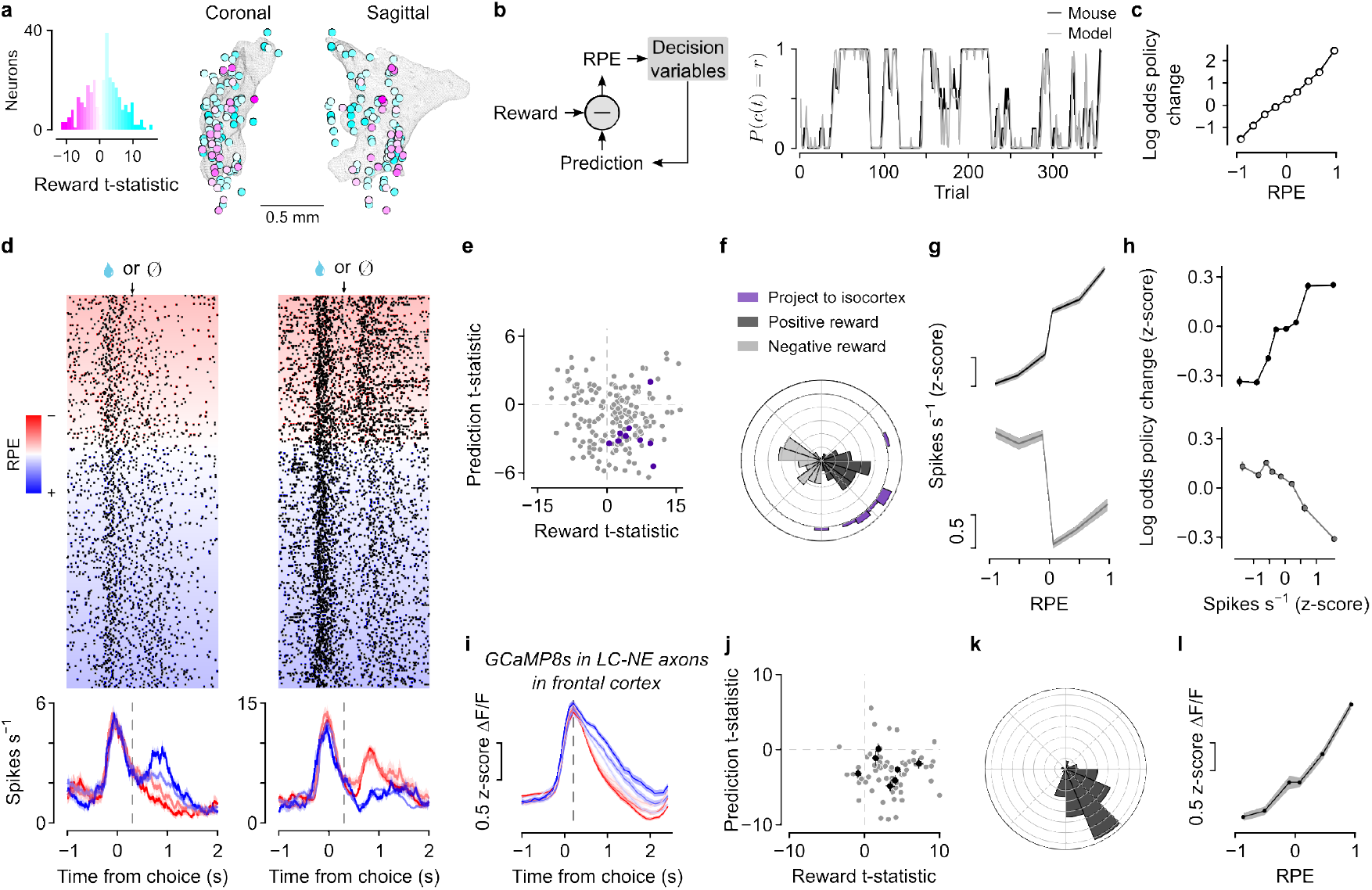
LC-NE projections to frontal cortex correlate with reward prediction error (RPE). **a**, Neurons excited by reward (cyan) or no reward (pink) were distributed nonuniformly in space (*p* = 3.0 × 10^−4^). **b**, Reinforcement-learning model. The predicted value of a choice is compared to reward to generate RPE. Right: model fit to the example in Figure 4b. Success rate of predicting choices, 0.87. Per-trial log likelihood, –0.24. **c**, Relationship between how choices changed on consecutive trials and RPE (rank correlation, ρ = 0.84, *p* = 0.0). **d**, Two example neurons with different outcome-related activity. Upper plots show spikes (ticks) for each trial (rows), sorted by RPE. Lower plots show mean ± SEM spike rates in RPE quartiles. Note that the first neuron’s response scales with RPE, whereas the second neuron is excited by lack of reward regardless of chosen value. **e**, Effects of chosen value (“prediction”) versus outcome (reward or no reward) on spike rates. Each *t*-statistic comes from a regression of a neuron’s spike rates on chosen value and outcome. Purple: neurons projecting to isocortex (antidromic stimulation and collision tests). **f**, Polar histogram of coefficients from regressions in (**e**) shows a nonuniform distribution of activity (Rayleigh test, *z* = 19.53, *p* = 2.3 × 10^−9^). Concentric circles denote 0.2 density. Purple: neurons projecting to isocortex are RPE-biased (Watson-Wheeler test, *W* = 0.04, permutation *p* = 0.01). **g**, Activity of LC-NE neurons excited or inhibited by reward versus RPE. **h**, Higher activity of neurons excited by reward correlated with larger change in choice likelihood. **i**, LC-NE axonal fluorescence in PL correlated with RPE (9 mice). **j**, Effects of chosen value versus outcome (reward or no reward) on axonal activity (compare to Figure 6e). Black: mean and SEM for each mouse. **k**, Polar histogram of regression coefficients of LC-NE axonal activity in cortex as a function of chosen value and outcome (compare to Figure 6f; Watson-Wheeler test, *W* = 0.1, permutation *p* = 2.0 × 10^−4^). Concentric circles denote 0.1 density. **l**, LC-NE axonal activity in frontal cortex correlates with RPE (compare to Figure 6g).

We calculated regressions of spike rates during the outcome (reward or no reward) as a function of the predicted reward value of the choice and outcome (Figure 6e). Neurons clustered in different regions of this plot: some were primarily excited by reward and inhibited by predicted reward (that is, they correlated with RPE), whereas other neurons were primarily excited by lack of reward without strong modulation by reward prediction. Neurons with increased activity in response to reward tended to lie more dorsal and anterior in LC than neurons with increased activity in response to no reward (Figures 6a and S18). To quantify this observation further, we calculated a polar histogram from the regression coefficients for reward prediction and reward in Figure 6e relative to the origin. We observed distinct clustering of the task-related activity of LC-NE neurons (Figure 6f ). Average responses of neurons of each population showed modulation by RPE or lack of reward (Figure 6g). Neurons that correlated with RPE or lack of reward had opposing correlations with choice behavior on subsequent trials (Figure 6h).

Do LC-NE neurons with different reward-related activity project to different regions? We tested this hypothesis by analyzing the activity of single LC-NE neurons projecting to the isocortex and the bulk calcium-dependent fluorescence of LC-NE axons innervating PL. If RPE is used to update decision variables in this task, we expect to observe LC-NE input to PL to correlate with RPE. LC-NE axonal activity in frontal cortex 1.5 s after choice outcome correlated with RPE (Figures 6i–6l). We quantified this observation by calculating regressions of single neurons that project to cortex and axonal ΔF/F as a function of chosen value and outcome. Regression *t*-statistics (Figure 6j) and coefficients (Figure 6k) were more positive than the population of single neurons (Fisher’s exact test, *p* < 0.01) and consistent with modulation by RPE. We thus conclude that frontal cortex receives input from dorsal LC-NE neurons whose activity increases during switch versus stay choices and correlates with RPE at the time of reinforcement.

## 3. Discussion

We discovered a topographic organization of the structure and function of mouse LC-NE neurons, one of the brain’s major neuromodulatory systems. LC-NE neurons project broadly in the CNS, but individual cells reach a limited set of structures. These projections are organized in space in the LC, with a dorsal-ventral distribution of somata that maps to the anterior-posterior distribution of axons. Gene expression is graded and organized in space, matching the patterns of projections to different CNS regions. LC-NE neuronal activity varies along the same spatial dimension, with dorsal cells, including those that project to isocortex, active when mice change a choice or receive a reward during a reinforcement-learning task. These findings add to a growing knowledge of heterogeneity among LC-NE neurons (Poe et al. 2020; Schwarz et al. 2015; Uematsu et al. 2017; Chandler et al. 2014; Breton-Provencher et al. 2022; Hughes et al. 2024; Luskin et al. 2025).

The graded molecular and anatomical organization of LC-NE neurons has parallels in other CNS structures. For example, the isocortex (Tasic et al. 2018), hippocampus (Cembrowski et al. 2016), thalamus (Turner et al. 2025), and striatum (Stanley et al. 2020) show graded gene expression in space, with specific locations on the gradient corresponding to projections to different target areas. Gene expression gradients may reflect an organizing principle of the CNS: continuous, spatial variation within a discrete cell class (Cembrowski & Menon 2018).

How do diverse synaptic inputs and biophysical properties of LC-NE neurons drive diverse choice and outcome activity? Major LC afferents include frontal cortex, central amygdala, cerebellum, hypothalamus, pons, and ventral medulla, as well as other neuromodulatory systems (Schwarz et al. 2015; Luskin et al. 2025; Breton-Provencher & Sur 2019; Cedarbaum & Aghajanian 1978; Aston-Jones et al. 1986; Luppi et al. 1995; Aston-Jones et al. 1991). Inputs from frontal cortex (Jodo et al. 1998) and central amygdala (Bouret et al. 2003) excite LC. Excitatory inputs from the medulla (Ennis & Aston-Jones 1988) may increase LC firing rates following aversive stimuli (Chiang & Aston-Jones 1993). In addition, connections between LC neurons (McKinney et al. 2023; Totah et al. 2018; Breton-Provencher & Sur 2019) may contribute to shaping the distinct outputs we measured here (e.g., increased activity with switch versus stay choices, or RPE versus lack of reward). It will be important to determine how biased inputs to different LC-NE neurons shape their diverse activity patterns.

We observed that LC-NE input to cortex correlates with RPE, a signal thought to be involved in learning. How could NE input to cortex drive learning? The mechanisms likely include multiple cell types and receptors. NE modulates synaptic plasticity across brain areas (Delaney et al. 2007; Seol et al. 2007; Carey & Regehr 2009; Hong et al. 2022). *Ex vivo* electrophysiology indicates different effects of NE on different cell types in cortex (Bergles et al. 1996; Kawaguchi & Shindou 1998; Dembrow et al. 2010; Dembrow & Johnston 2014). NE receptors are expressed nonuniformly across cortical cell types (Scheinin et al. 1994; Rosin et al. 1996; Day et al. 1997; Talley et al. 1996; Santana et al. 2013; Santana & Artigas 2017). In addition, NE may exert its effects on both neuronal and non-neuronal cells (Bekar et al. 2008; Paukert et al. 2014; Mu et al. 2019), providing a rich substrate for broadcasting an ascending learning signal across large volumes of cortex. This learning signal may generalize beyond reinforcement learning: LC-NE axonal activity in visual cortex correlates with sensory-motor errors (Jordan & Keller 2023), and artificial excitation of LC changes sensory-driven learning (Jordan & Keller 2023; Glennon et al. 2019; McBurney-Lin et al. 2022).

We conclude that a major neuromodulatory input to cortex signals RPE, a key variable for reinforcement learning. It is striking that dopamine inputs to basal ganglia carry a similar RPE, yet project comparatively little to cortex (Nomura et al. 2014; Descarries et al. 1987) (for differences between rodents and primates, see (Berger et al. 1991; Lewis et al. 2001)). We propose that cortex and basal ganglia may receive complementary inputs from the two major catecholamines in the brain: NE and dopamine. These two neuromodulators may allow cortex and basal ganglia to operate as hierarchical learning systems (J. X. Wang et al. 2018), allowing the brain to learn over multiple timescales (Murray & Escola 2020).

## 4. Methods

All surgical and experimental procedures were in accordance with the *National Institutes of Health Guide for the Care and Use of Laboratory Animals* and approved by the Animal Care and Use Committees of the Allen Institute or Johns Hopkins University.

### 4.1 Definition of LC-NE neurons

We used *Dbh*-Cre mice (Tillage et al. 2020) (Dbh^tm3.2(cre)Pjen^, The Jackson Laboratory, 033951) backcrossed with C57BL/6J mice (The Jackson Laboratory, 000664). *Dbh*-Cre mice were crossed to RCL-H2B-GFP mice (The Jackson Laboratory, 036761), and the native fluorescence of the labeled nuclei was imaged on a SmartSPIM microscope after tissue clearing using LifeCanvas active delipidation, agar embedding, and refractive-index matching using EasyIndex (SmartSPIM, LifeCanvas Technologies). Images from 8 brains were acquired at 1.8 × 1.8 × 2.0 µm per voxel. Raw image data were passed through a customized 3D UNet (Trailmap) (Friedmann et al. 2020) with a model trained for immunolabeled nuclei (Friedmann et al. 2024). The resulting probability maps were convolved with a spherical kernel approximating a nucleus diameter and then centroids were estimated with 3D local maxima detection. Stitching and registration to the CCFv3 followed the SmartSPIM Pipeline (https://github.com/AllenNeuralDynamics/aind-smartspim-pipeline).

Points were subsequently transferred to the Allen atlas reference space via an ANTS transform (Friedmann et al. 2020). After restricting analysis to caudal midbrain and dorsal pons, and after reflecting all points into a single hemisphere, local densities were estimated via nearest neighbor analysis and points at uniform density levels were used to define meshes. Briefly, normals for points were estimated and used to generate surfels which were wrapped in a watertight mesh (Perens et al. 2021) by isosurface extraction (https://www.github.com/fwilliams/point-cloud-utils). A mesh representing LC-core at the 67th percentile represents the density threshold at which sub-coeruleus cleanly separates from the main portion of LC. Initial mesh estimates were made using dynamic radius and resolution parameters to compensate for point density and mesh size across the displayed percentiles.

### 4.2 Single-neuron reconstructions

Whole mouse brain specimens were prepared for single neuron reconstructions using previously published methods (Glaser et al. 2025) (dx.doi.org/10.17504/protocols.io.n92ldpwjxl5b/v1). Briefly, 2 male and 2 female adult *Dbh*-Cre mice received systemic injections, via the retro-orbital sinus, of a 100 µL mixture of Cre-dependent Tet transactivator (AAV-PHP-eB_Syn-FlexTRE-2tTA, Addgene plasmid id: 191210; dosage range 1.0 × 10^8^ gc/mL – 3.0 × 10^9^ gc/mL) and a reporter virus (either AAV-PHP-eB_7x-TRE-3xeGFP or AAV-PHP-eB_7x-TRE-tdTomato, typical dose 1.8 × 10^11^ gc/mL; 191206 and 191207). Viruses were obtained from either the Allen Institute for Brain Science viral vector core, the University of North Carolina, or the BICCN-Neurotools core and were prepared in an AAV buffer consisting of 1xphosphate-buffered saline (PBS), 5% sorbitol, and 350 mM NaCl. Six to eight weeks after viral transfection, mice were anesthetized with an overdose of isoflurane and transcardially perfused with 10 mL 0.9% saline at a flow rate of 9 mL/min followed by 50 mL 4% paraformaldehyde in PBS at a flow rate of 9 mL/min. Brains were extracted and post-fixed in 4% paraformaldehyde at room temperature for 3–6 hr and then left at 4 ◦C overnight (12–14 hr). The following day, brains were washed in 1xPBS to remove all traces of excess fixative. Subsequent tissue processing including clearing, immunolabeling, and whole brain expansion steps were carried out as previously described (Glaser et al. 2025) (dx.doi.org/10.17504/protocols.io.n92ldpwjxl5b/v1). Gelled brains were soaked in 0.05x saline sodium citrate (SSC) to achieve approximately 3x expansion, 24 h prior to imaging the expanded brains were equilibrated in a solution of 0.05x SSC that contained 10 mM ascorbic acid included as an antifade.

Expanded brains were imaged on a custom SPIM microscope (ExA-SPIM) (Glaser et al. 2025) (dx.doi.org/10.17504/protocols.io.6qpvrwe33lmk/v1). Brains were imaged with both 488 nm and 561 nm excitation and an 8x binned autofluorescence image volume was collected for the “non-signal” channel for registration to the CCF. All subsequent data processing steps including image illumination correction, stitching and fusion into a coherent image volume were carried out via automated cloud pipelines (Glaser et al. 2025).

To generate whole-brain single neuron reconstructions, we used HortaCloud (Rokicki et al. 2025), an open-source streaming 3D annotation platform enabling fast visualization and collaborative proofreading of terabyte-scale image volumes. Using this platform, human annotators proofread individual neurons via a web browser on a personal workstation. Proofreading entailed starting from the soma and tracing out all axonal and dendritic segments using a depth first search tree traversal approach (Winnubst et al. 2019; Economo et al. 2016). Neuronal trees generated by manual placement of control points were refined offline to generate dense (one node per voxel) and sparse (uniform sub-sampling of the dense tree) representations of the reconstruction data before use for subsequent quantitative analysis.

#### Models of single-neuron axonal distributions

We developed two models to generate whole-brain distributions of axons to compare to the distribution of LC-NE axons (Figures 2 and Figure S4c). We first create an axonal projection data matrix where each row is a reconstructed neuron, and the columns are coarse brain regions. Each entry in the matrix is the fraction of that cell’s reconstructed axon that lies within that coarse brain region. The coarse brain regions we used were: olfactory areas (OLF), isocortex (CTX), hippocampal formation (HPF), cortical subplate (CTXsp), cerebral nuclei (CNU), thalamus (TH), hypothalamus (HY), midbrain (MB), cerebellum (CB), pons (P), medulla (MY), cerebellum related fiber tracts (cbf ), lateral forebrain bundle system (lfbs), medial forebrain bundle system (mfbs), and ventricular systems (VS).

The first model (“random projections”) posits that each LC-NE neuron projects randomly across the brain. We assess this model by shuffling the projection matrix across brain regions and comparing the shuffled data with the original. As expected, the shuffled data loses the structured correlation matrix of the original data. This model ignores important anatomical constraints of the brain, namely that an axon cannot go straight from the pons to isocortex.

The second model (“Markov with renewal”) posits that LC-NE neurons distribute their axons by a Markov process walk across the brain subject to anatomical constraints of how brain regions are connected, and constraints on the minimum and maximum axonal length. We first created a transition matrix between coarse brain regions by averaging across all LC-NE neurons in our dataset. Using the graph representation of our axonal reconstructions, we iterate through each segment of each cell’s axon. Each segment of the reconstructions is approximately 10-µm in length. For each segment, we tally the coarse brain region where that segment starts and stops. After tallying across all cells, we normalize the matrix into a transition matrix where each entry represents the probability of transitioning from one coarse brain region into another. An axonal branch that is near a region boundary might artificially inflate the rate of transitions between regions. To avoid this, we only counted transitions between two brain regions if the axon stayed in the new region for a minimum of 2 mm.

Each simulated neuron starts in the LC, which in our coarse brain regions is part of the pons. Our simulations take 10-µm steps. On each step, the axon branch transitions of a new brain region according to the transition matrix, as well as branches or terminates. Branching and termination rates were learned from data and are dependent on axon length from the soma but not the current brain region. Over the first 10 cm from the soma, both rates increase with the branching rates larger than the termination rate. After 10 cm from the soma, both rates are roughly constant and equal. Each axon branch grows independently according to the same Markov process. We add an additional constraint of a minimum axon length (summed across branches) of 10 mm to match the shortest neuron in our dataset. The last axon branch of our simulated neurons was not allowed to terminate until this minimum threshold was met. Finally, we add a Weibull renewal process that stops all active axon processes. The Weibull renewal process has a hazard function that is dependent on the overall axon length of the cell (summed across branches) and then results in a Weibull distribution of axonal length. We fit a Weibull distribution to the LC-NE neurons in our dataset, using neurons with complete reconstructions as uncensored observations, and neurons with incomplete spinal cord projections as censored observations.

This model is not an exact replica of the data, but gets surprisingly close. The model under-projects to the isocortex, while over-projecting to the thalamus, and to the cerebellum. The correlation matrix matches the anterior/posterior block structure observed in the data. Our process for grouping each axon segment into coarse brain regions is imperfect, especially around narrow fiber tracts. This explains why the model creates a stronger correlation between the cerebellum and brainstem structures then is observed in the data.

#### Somatodendritic properties

We quantified somatic bipolarity by fitting a 3D gaussian distribution to a local patch of the image volume around each cell’s soma center. The square root of the largest eigenvalue of the covariance matrix (normalized length of the principal axis) was taken as the somatic bipolarity coefficient. A similar approach was applied to dendrites, by first calculating a unit vector in the direction of each dendritic “stem” point (primary branches within 50 µm or points where each dendrite crosses 50 µm radius from the soma center), then finding the normalized length of the principal axis of the set of dendritic stem direction vectors. We also calculated the angular separation between the somatic and dendritic primary axes, to show that each independently capture the neurons’ bipolarity. The harmonic mean of these bipolarity metrics was used as a combined bipolarity coefficient.

### 4.3 Retrograde tracing

Heterozygous *Dbh*-Cre mice crossed with homozygous Ai65 mice (The Jackson Laboratory, 021875; P40–P50) were anesthetized with 2.0% isoflurane in O_2_ and maintained at 0.7–1.0% isoflurane throughout the surgery. Mice were mounted in a stereotaxic frame (David Kopf Instruments) for injections of AAV pEF1a-DIO-FLPo-WPRE-hGHpA (Addgene 87306-AAVrg, lot v188462; Table 1). Small craniotomies were made in the skull above the injection target for intracerebral injections (dx.doi.org/10.17504/protocols.io.14egn8ewzg5d/v8 for iontophoretic injections and dx.doi.org/10.17504/protocols.io.bp2l6nr7kgqe/v8 for pressure injections). Spinal cord injections were made between C4-C5 vertebrae without drilling (dx.doi.org/10.17504/protocols.io.yxmvm8mq9g3p/v1). Intracerebral injections were in the right hemisphere. Spinal cord injections were bilateral.

**Table 1:** Virus injections for retrograde tracing experiments. Summary of animal and cell counts by projection target and labelled soma location. Spinal cord injections were made bilaterally. All other injections were in the right hemisphere. Injection coordinates are relative to bregma (anterior-posterior, AP; medial-lateral, ML; dorsal-ventral, DV).

| Injection target (mice, cells) | Injection type | Injection volume | Coordinates from bregma (AP, ML, DV) (mm) |
| --- | --- | --- | --- |
| MOB (4, 928) | Pressure | 150 nL per insertion | (4.28, 0.50, -2.30); (4.28, 1.20, -0.70) |
| CTX: PL, MO (5, 2,714) | Pressure | 300 nL per insertion | (2.10, 0.33, -1.50); (1.70, 0.90, {-0.80, -0.40}) |
| TH: VAL, VPM (7, 327) | Iontophoresis | 5–7 min, 3 μA per insertion | (-0.82, 1.13, -3.28); (-1.82, 1.60, -3.30) |
| PAG (6, 1,546) | Pressure | 200 nL | (-4.45, 0.40, -1.40) |
| CB: culmen, anisiform, paramedian (6, 3,829) | Pressure | 200 nL, 300 nL | (-5.75, 0.00, -0.50); (-7.50, 2.00, {-1.75, -0.90}) |
| SP: C4–C5 (7, 4,321) | Pressure | 100 nL per insertion | (N.A., 0.35, -0.90); (N.A., -0.35, -0.90)) |

Mice were euthanized with an isoflurane overdose 4–6 weeks after AAV injection and transcardially perfused with PBS followed by 4% paraformaldehyde (PFA EMS 50-980-495 or equivalent; dx.doi.org/10.17504/protocols.io.8epv51bejl1b/v8). Brains were dissected, post-fixed in 4% PFA overnight, and transferred to PBS. Tissue was cleared using LifeCanvas active clearing, followed by agar embedding, refractive-index matching using EasyIndex solution, and imaging (SmartSPIM, LifeCanvas Technologies).

Retrogradely labeled LC-NE neurons were identified by tdTomato fluorescence in the soma and primary processes. Positions of retrogradely labeled cells in whole-mount stitched and CCF-registered samples were identified either by manual labeling or automated segmentation followed by manual verification. Labeling accuracy was examined in at least two distinct planes (coronal, horizontal).

For automated segmentation, brain volumes were divided into 3D chunks (512 x 512 x 512) optimized for OME-Zarr image processing. Each block underwent background subtraction using the photutils photometry package (Bradley et al. 2025). Processed images were passed to a modified version of the CellFinder algorithm (Tyson et al. 2021), where a median filter followed by a Laplacian of Gaussian (LoG) filter were applied to each slice along the imaging plane. A threshold 6 standard deviations above the mean was used to identify potential cell locations and the image was binarized. 2D regions were merged via an ellipsoid filter and large regions, resulting from the merging of cells, were split using iterative ellipsoid filters. To minimize false positives from fluctuations in tissue autofluorescence, cell proposals from the detection phase were passed to an 18-layer ResNet for classification. For each proposal a small block (28 x 28 x 50) centered on the proposed cell location was taken from the signal, as well as the autofluorescence channel. The input image to the network was downsampled by a factor of 2 across each dimension prior to classification to minimize the contributions of fluorescent signal from axonal and dendritic processes. Interactive Neuroglancer images were created using proposals classified as true cells, and a final round of manual refinement was done to remove any remaining false positives and add false negatives.

Annotations were registered to CCFv3 using the affine and warp transforms calculated during image registration. Annotations related to the LC were identified using a signed distance function comparing point locations within the CCF relative to the exterior mesh of the pons. All annotations were categorized as existing inside, outside, or on the boundary of the mesh. Retained points inside and on the pons mesh boundary were then manually examined to remove false positive cells and add back in false negatives.

### 4.4 BARseq and MAPseq

The sindbis virus barcode library HZ120 (Yuan et al. 2024) was generated by the MAPseq core facility at Cold Spring Harbor Laboratory (CSHL) and used for MAPseq and BARseq experiments as described previously (X. Chen et al. 2019). The HZ120 library exhibits a diversity of approximately 8 million barcodes and was not fully sequenced *in vitro*.

Eight-week-old C57BL/6J mice were anesthetized using oxygenated 4% isoflurane and maintained with oxygenated 0.7–1.0% isoflurane throughout the surgery. We injected 300 nL of HZ120 sindbis virus in LC in each hemisphere at AP –5.2; ML 0.85; DV –3.20, –2.90 using pressure injection (NanoInject III; dx.doi.org/10.17504/protocols.io.bp2l6nr7kgqe/v8). Mice recovered and were euthanized 22–28 h post-injection.

Brains were dissected (dx.doi.org/10.17504/protocols.io.j8nlken35l5r/v6), embedded in OCT and snap-frozen in an ethanol dry-ice bath. Extruded spinal cords were flash frozen, straightened out on a razor blade secured in a conical tube. Samples were stored at –80 ◦C until cryosectioning (Leica 3050S) in the coronal plane (dx.doi.org/10.17504/protocols.io.3byl4p18rlo5/v1)). We cut 300 µm slices for regions outside LC (MAPseq) and 20 µm for tissue containing LC (BARseq). To avoid cross-contamination, we used a fresh unused part of a blade to cut each slice and cleaned the brush and the holding platform with 100% ethanol between slices. Cryostat chamber temperature was allowed to equilibrate for 30 min to account for temperature changes between cutting thick and thin slices.

For MAPseq experiments, 3 × 300 µm coronal sections were mounted in a row onto Superfrost Plus Gold slides. For BARseq experiments, 4 × 20 µm sections were mounted onto Superfrost Plus Gold slides according to microfluidic chamber template. Once all cut sections were melted onto the slide, slides were rapidly frozen on dry ice and stored at –80 ◦C in slide boxes in vacuum sealed bags until microdissection (MAPseq) or library prep (BARseq).

Regions of interest were dissected from each chilled brain slice with microscalpels pre-chilled (Fine Science Tools 10316-14). Glass slides were kept on dry ice and placed on a pre-chilled metal platform mounted above a dry ice/ethanol bath. Sections were first bisected down the midline. Then major anatomical regions (e.g., isocortex, hippocampus, midbrain) were cut along their respective boundaries. Each cut was performed with a cleaned blade. Blades were rinsed in consecutive 4 × 50 ml conical tubes containing ethanol stored on ice. To keep blade temperature consistently low, 3–4 scalpels were rotated and stored on the metal platform. After right hemisphere dissection was completed, a reference image for dissection boundaries was taken again for each of the 3 sections. After that, the left side was dissected along the same boundaries. Individual tissue pieces were then collected into pre-chilled microtubes (Qiagen 19560 tubes, 19566 caps) on dry ice, capped and stored at –80 ◦C. Left and right hemisphere samples were kept separate, and corresponding regions of interest were combined across 3 consecutive 300-µm sections mounted on the same slide. To assess cross-sample contamination, we collected tissue from uninjected brains as negative controls. After all regions of interest were collected, with samples kept on dry ice we added pre-chilled Qiagen bead (69989) to each tube. Spinal cord samples then received 800 µl Trizol, while the rest used 400 µL Trizol (Thermofisher 15596026). Samples were shipped overnight on dry ice to the CSHL MAPseq facility.

RNA extraction and sequencing library preparation were performed following the protocol (dx.doi.org/10.17504/protocols.io.bsm9nc96), as described previously (X. Chen et al. 2019; Han et al. 2018; Kebschull et al. 2016). Briefly, total RNA for each region was extracted using Trizol (Thermo Fisher Scientific, 15596018) and eluted in 13 µL of H_2_O. Before library preparation, a few randomly selected samples were run on a Bioanalyzer (Agilent, 5067-1513) to ensure good RNA quality. To prepare the library for sequencing, we mixed 4 µL total RNA of each sample with 1 µL spike-in RNA (GTC ATG ATC ATA ATA CGA CTC ACT ATA GGG GAC GAG CTG TAC AAG TAA ACG CGT AAT GAT ACG GCG ACC ACC GAG ATC TAC ACT CTT TCC CTA CAC GAC GCT CTT CCG ATC TNN NNN NNN NNN NNN NNN NNN NNN NAT CAG TCA TCG GAG CGG CCG CTA CCT AAT TGC CGT CGT GAG GTA CGA CCA CCG CTA GCT GTA CA), where ATCAGTCA is the barcode tag of the spike-in. Spike-in RNA was transcribed in vitro by T7 RNA polymerase and diluted into1x104 molecules/µL. Barcode mRNA was reverse-transcribed into the first-strand cDNA using gene-specific barcoded reverse transcription primers and SuperScript IV (Thermo Fisher Scientific, 18090010). The barcoded reverse transcription primer sequence was 5’-CTT GGC ACC CGA GAA TTC CAX XXX XXX XXX XXZ ZZZ ZZZ ZTG TAC AGC TAG CGG TGG TCG-3’, where X12 are the barcoded unique molecular identifiers (UMI) and Z8 are barcoded sample specific identifiers (SSI). The synthesized first strand cDNA of each sample was labeled with a 12-nt UMI (unique for each RNA molecule) and 8-nt SSI (unique for each sample). After synthesis of the first strand cDNA, every 7–8 samples of target brain regions were pooled together for clean-up by 1.8 x AMPure XP beads (Beckman Coulter, A63881) and synthesis of the second strand cDNA (Thermo Fisher, A48571). The double-stranded cDNA samples were cleaned up by 1.8 x AMPure XP beads and treated with Exonuclease I (New England Biolabs, M0293S) to remove the single-stranded reverse transcription primers before PCR amplification.

The double-stranded cDNA samples were amplified by two rounds of polymerase chain reactions (PCR) using standard Accuprime Pfx protocol (Thermo Fisher Scientific, 12344032) with 2 min extension for each cycle. Primers 5’-CTG TAC AAG TAA ACG CGT AAT G-3’ and 5’-CAAG CAGAAGACGGCATACGAGATCGTGATGTGACTGGAGTTCCTTGGCACCCGAGAATTCCA-3’ were used for the first PCR reaction, and primers 5’ -AATGATACGGCGACCACCGA-3’ and 5’ -CAAGCAGAAGACGGCATACGA-3’ were used for the second PCR. In the first PCR, the pooled cDNA samples were amplified for 15 cycles in 250 µL of reaction volume per 7–8 samples. After treatment with Exonuclease I (New England Biolabs, M0293S) to remove excess primers, all the PCR1 products of the target areas were pooled together. A quarter of the pooled PCR1 products of target areas were amplified by 10 cycles in 12 mL of PCR II reaction volume. The PCR2 products were cleaned up and concentrated by SV wizard PCR cleanup kit (Promega, A9282) and loaded into a 2% agarose gel for electrophoresis. The 233 bp PCR product was cut from the agarose gel, cleaned up by Qiagen MinElute Gel Extraction Kit (Qiagen 28606) and quantified by Bioanalyzer (Agilent, 5067-4626) and qPCR.

The pooled library was sequenced on two lanes of Illumina Nextseq550 high output at paired end 36 with Illumina compatible Read 1 Sequencing Primer and Illumina compatible small RNA read 2 primer as described previously (Kebschull et al. 2016).

#### MAPseq data processing

The raw MAPseq data consist of FASTQ files, where paired end 1 covers the 30 nt barcode sequence and paired end 2 covers 12 nt UMI and the 8 nt SSI (Kebschull et al. 2016). The raw sequencing data were processed with code available at: https://github.com/ZadorLaboratory/mapseq-processing We first assembled the reads from the paired-end input, trimming each to the length of the sequence of interest. These sequences were then aggregated by exact duplicate and the read count for each unique sequence kept. We removed any data with ambiguous bases or long runs (>7) of the same base. We labeled and categorized all the reads by SSI sequence and expected spike-in sub-sequence. To account for expected replication, amplification, and sequencing errors, we identified all barcodes that differed from each other by an edit distance of 3 or less. This parameter is informed by the barcode sequence length, and the known diversity of the virus library. All groups of barcodes were then collapsed to the most common variant, as measured by read count. We then aggregated by barcode, SSI, and UMI sequence, retaining the number of UMIs in each. To exclude low-confidence projections, we required a barcode to have at least 5 molecule counts in at least one projection region. We normalized the variation in library preparation and sequencing across different samples within each brain based on recovered spike-in RNA as described previously (Han et al. 2018).

#### BARseq library preparation

BARseq samples for *in situ* sequencing were prepared using a protocol described previously (https://dx.doi.org/10.17504/protocols.io.n2bvj82q5gk5/v1) (X. Chen et al. 2019; Sun et al. 2021; X. Chen et al. 2025) with modifications (dx.doi.org/10.17504/protocols.io.kqdg3ke9qv25/v1). Briefly, brain sections were fixed in 4% PFA in PBS for 1 hr, dehydrated through 70%, 85%, and 100% ethanol, and incubated in 100% ethanol for 1.5 hr at 4 ◦C. After rehydration in PBST (PBS and 0.5% Tween-20), slices were incubated in the reverse transcription mix (RT primers, 20 U/µL RevertAid H Minus M-MuLV reverse transcriptase, 500 µM dNTP, 0.2 µg/µL BSA, 1 U/µL RiboLock RNase Inhibitor, 1×RevertAid RT buffer) at 37 ◦C overnight.

On day two, cDNA was cross-linked using BS(PEG)9 (40 µL in 160 µL PBST) for 1 hr, the remaining crosslinker neutralized with 1 M Tris pH 8.0 for 30 min, followed by a wash with PBST. Sections were then incubated with non-gap-filling ligation mix (1xAmpligase buffer, padlock probe mix for endogenous genes, 0.5 U/µL Ampligase, 0.4 U/µL RNase H, 1 U/µL RiboLock RNase Inhibitor, additional 50 mM KCl, 20% formamide) for 30 min at 37 ◦C and 45 min at 45 ◦C. Then sections were incubated in gap-filling ligation mix (same as the non-gap-filling mix with the Sindbis barcode padlock probe [XCAI5] as the only padlock probe, and with 50 µM dNTP, 0.2 U/µL Phusion DNA polymerase, and 5% glycerol) for 5 min at 37 ◦C and 45 min at 45 ◦C. After the second round of ligation, samples were washed with PBST, hybridized with 1 µM RCA primer (XC1417) for 10 min in 2×SSC with 10% formamide, followed by two washes in 2×SSC with 10% formamide and twice in PBST, then incubated in the RCA mix (1 U/µL phi29 DNA polymerase, 1xphi29 polymerase buffer, 0.25 mM dNTP, 0.2 µg/µL BSA, 5% glycerol (extra of those from the enzymes), 125 µM aminoallyl dUTP) overnight at room temperature to complete rolling circle amplification. On day three, rolonies were crosslinked using BS(PEG)9 (40 µl in 160 µL PBST) for 1 hr, the remaining crosslinker neutralized with 1 M Tris pH 8.0 for 30 min, followed by a wash with PBST.

#### BARseq sequencing and imaging

Sequencing was performed using Illumina MiSeq Reagent Nano Kit v2 (300-cycles) and imaged on a Nikon Ti2-E microscope with Crest xlight v3 spinning disk confocal, photometrics Kinetix camera, and Lumencor Celesta laser. All images were taken with a Nikon CFI S Plan Fluor LWD 20X 0.7NA non-immersion objective. For each sample, first seven sequencing cycles probe the selected gene panel, followed by 15 sequencing cycles for barcodes, followed by hybridization cycle. In each imaging round, a z-stack of 10 images centered around the region of interest was captured with a 1.5 µm step size at each field of view (FOV). Adjacent FOVs had an overlap of 24%.

A detailed protocol is available here (dx.doi.org/10.17504/protocols.io.81wgbp4j3vpk/v2). Briefly, for sequencing the first gene cycles, the sequencing primer (YS220) was hybridized in 2×SSC with 10% formamide for 10 min at room temperature. Subsequently, the sample underwent two washes in 2×SSC with 10% formamide and twice in PBS2T (PBS with 2% Tween-20). Following this, the sample was incubated in MiSeq Incorporation buffer at 60 °C for 3 min, followed by a wash in PBS2T. The sample was then exposed to Iodoacetamide (9.3 mg vial in 2.5 mL PBS2T) at 60 ◦C for 3 min, followed by a single wash in PBS2T and twice in Incorporation buffer. It was then subjected to two incubations in IMS at 60 ◦C for 3 min, followed by four washes with PBS2T at 60 ◦C for 3 min. Finally, the sample was washed and imaged in SRE. For sequencing the first barcode cycle, the sequenced products were initially stripped with three 10-minute incubations at 60 ◦C in 2×SSC with 60% formamide. Subsequently, the sample was washed with 2×SSC with 10% formamide, and the barcode sequencing primer (XCAI5) was hybridized. The sequencing process then proceeded similarly to the first gene cycle.

For subsequent sequencing cycles for both genes and barcodes, the sample underwent two washes in Incorporation buffer, followed by two incubations in CMS at 60 ◦C for 3 min. This was succeeded by two more washes in Incorporation buffer, after which the sample was treated with iodoacetamide as in the first sequencing cycle. During hybridization cycles, the sequenced products were stripped with three 10-minute incubations at 60 ◦C in 2×SSC with 60% formamide. The sample was then washed with 2×SSC with 10% formamide. Following this, the hybridization probes (XC2758, XC2759, XC2760, YS221) were hybridized for 10 min at room temperature. The sample was subsequently washed twice with 2×SSC with 10% formamide and incubated in PBST with 0.002 mg/mL DAPI for 5 min. Finally, the sample was transferred to SRE for imaging. For one brain all the above steps were completed manually. For the subsequent samples *in situ* sequencing was performed on an automated microfluidics set up mounted on the microscope using the same reagents, temperatures and incubation durations.

### Analysis of MAPseq/BARseq data

#### BARseq data processing

BARseq data were processed according to previous methods (Sun et al. 2021) with slight adjustments. Max-projections were generated from the image stacks, followed by noise reduction using Noise2Void (Krull et al. 2019). Subsequently, background subtraction, correction for channel shift and bleed-through, and registration of images across all sequencing cycles were applied. Cell segmentation was performed using Cellpose (Stringer et al. 2021), while rolonies were decoded using BarDensr (S. Chen et al. 2021), and then assigned to cells. During BarDensr decoding, negative control GIIs (GIIs not associated with any padlock probe) were utilized to estimate the false discovery rate (FDR), with the decoding threshold automatically adjusted to target approximately 5% FDR. Whole-slice images were generated by stitching individual imaging FOVs. Processing steps were applied separately to each FOV to prevent stitching errors, minimize alignment artifacts, and facilitate parallel processing. Stitched images were solely used to generate a transformation, applied to each rolony and cell after completing all other steps.

Overlapping cells from neighboring FOVs were identified using a custom implementation of sort and sweep collision detection algorithm (Baraff & Witkin 1992). In cases where two or more cells overlapped, cells with higher read counts (indicating higher read quality) were retained. Each stitched slice was registered to Allen CCFv3 using manual registration. QuickNii (v.3 2017) was used to manually select the CCF plane for each slice, followed by alignment of area borders within each slice using Visualign (v. 0.9) using nonlinear adjustments to the coronal plates.

#### BARseq transcriptomic analyses

MATLAB outputs from the BARseq data processing pipeline (.mat, v7.3/HDF5) were imported into R by reconstructing the sparse gene-by-cell count matrix from the stored CSC components and assembling per-cell metadata into a SingleCellExperiment object. Cells were assigned stable unique identifiers by concatenating batch number, slice, and cell ID. Negative control genes were removed from the data matrix. When duplicated gene symbols were present, the duplicate entry corresponding to a hybridization cycle was removed.

To identify transcriptomic types, an iterative clustering pipeline adapted from single-cell RNA sequencing (scRNAseq) studies (Amezquita et al. 2020) was used. Cells were QC filtered prior to normalization and clustering to ≥ 20 total counts and ≥ 5 detected genes to ensure correct transcriptomic identity assignment. Counts were normalized to CP10 (counts per 10) values and then subjected to log1p transformation. As the panel consisted of pre-selected genes, highly variable gene selection was skipped, and PCA was conducted on all probed genes. Unsupervised clustering was performed on log-normalized expression matrix using PCA (30 components), UMAP on the PCA space (100 neighbors), and shared-nearest-neighbor graph clustering (*k* = 50 neighbors; Jaccard weighting) followed by Louvain community detection to assign cluster labels.

LC neuron isolation by iterative clustering and anatomical gating. LC-NE neurons were isolated by iterative clustering. First, all neurons were clustered using the PCA/UMAP/SNN-Louvain workflow described above. For each sample, the cluster enriched for canonical NE markers (*Dbh, Th, Ddc, Slc18a2*) was identified and subset. The subset was then renormalized to CP10, log1p-transformed and reclustered to increase resolution within the LC-enriched compartment. At each iteration, candidate LC/LC-NE clusters were evaluated using (i) marker enrichment and (ii) anatomical localization across slices using CCF coordinates. Clusters whose cells localized clearly outside the LC region were excluded as entire clusters.

As the final cleanup step to obtain high-confidence LC-NE neurons, spatial coherence and density filtering was applied after the last round of LC-NE refinement. Using CCF coordinates (ML and DV converted to µm by ×25) and a depth coordinate defined as slice index × 20 µm, k=10 nearest neighbors were computed per cell in 3D space. Spatial coherence was defined as distance-weighted cluster purity: the fraction of neighbors sharing the same Louvain label, weighted by 1/(distance+ϵ) (ϵ = 10^−6^). Spatial density was defined as 1/(mean kNN distance+ϵ). Cells were retained if spatial coherence ≥ 0.05 and spatial density was at least the 5th percentile of the dataset-wide density distribution. Cells failing either criterion were excluded, yielding the final high-confidence LC-NE cell set used for all subsequent downstream transcriptomic and projectomes analyses.

#### BARseq–MAPseq barcode matching

BARseq 15-cycle barcodes from high-confidence, barcoded LC-NE neurons were converted from numeric base calls to nucleotide strings using a fixed mapping (1=G, 2=T, 3=A, 4=C). MAPseq 32 nt barcodes were truncated to 15 nt to match BARseq length. For each BARseq cell, candidate MAPseq matches were identified by Hamming distance at thresholds of 0–3 mismatches. Matches were resolved conservatively by retaining only the minimum-distance hit per cell and accepting a match only if (i) a single best hit remained and (ii) the matched MAPseq 15-mer was unique in the MAPseq dataset (15-mer collisions excluded). MAPseq projection tables were exported for each mismatch threshold. False-positive match rates were estimated by applying the same matching and resolver rules to (i) random 15-mer barcodes and (ii) shuffled BARseq barcodes (with the 9th nucleotide fixed and the remaining positions permuted), averaging across 20 runs. For each mismatch threshold, the fraction of simulated barcodes that were selected as unique best matches was used as the estimated false-positive rate under the full resolver. All projection analyses used data from one Hamming distance mismatch as FP rate was estimated under 1%.

After matching, we removed duplicate entries arising from segmentation/FOV overlaps or barcode reuse. Duplicate structure was detected in two ways: (i) identical MAPseq profiles across all ROIs and (ii) repeated BARseq barcodes. All identical MAPseq profiles were the result of repeat BARseq barcodes. For these duplicate BARseq groups, segmented cell images were exported and manually reviewed. If the same barcode appeared in different structures (barcode reuse/mixed sources), all entries for that barcode were discarded. If entries were the same biological cell segmented more than once, the better-segmented instance (operationalized by manual review, with transcriptomic QC metrics such as total UMI counts) was retained and the others discarded.

Projection matrix construction and harmonization. Per-brain matched MAPseq projection tables were converted into analysis-ready projection matrices using brain-specific loader functions. First, matched barcodes were filtered using the curated blacklists derived from manual review of duplicate/ambiguous barcoded cells. Next, MAPseq ROIs were renamed using sample metadata to encode target identity, hemisphere it was dissected from and slide number. Then we matched uniquely barcoded MAPseq cells to BARseq metadata, and soma hemisphere was assigned from CCF coordinates.

To enable cross-brain data pooling, per-brain matrices were harmonized to a shared ROI set. This included resolving minor ROI naming inconsistencies, combining predefined subdivisions, and collapsing slide-specific replicates by summing columns that shared the same base ROI and hemisphere. As an internal integrity check, hemisphere assignment for selected targets was evaluated by comparing mean projection strengths to left versus right targets stratified by soma hemisphere. In one instance when a consistent inversion was detected, the corresponding RH/LH target columns were swapped.

Per brain ipsilateral and contralateral matrices were then created by relabeling projection columns relative to soma hemisphere and retaining only common ROIs across RH and LH samples. Projection strengths were row-normalized by each cell’s total projection count, log1p transformed, and scaled by the global maximum value for the given sample. Normalized matrices were concatenated across brains by row-binding. Metadata tables were combined and reordered to match the final matrix row order.

#### Projection pattern analyses

The harmonized ipsi/contra-normalized projection matrix was visualized as heatmaps. Cells were ordered by their top-projection region, defined as the region with the maximum normalized value per cell.

#### Regional module aggregation and coarse projection-pattern visualization

To align single-neuron MAPseq projection profiles with the coarse anatomical summaries used in ExA-SPIM, we mapped fine-grained target ROIs to 12 regions (OLF, CTX, HPF, CTXsp, CNU, TH, HY, MB, CB, P, MY, SP) and summed ipsi/contra-normalized projection strengths within each group to generate a cell × group matrix. Values were then row-normalized to fractions of each neuron’s total grouped projection strength (rows sum to 1) and visualized as a heatmap after ordering neurons by their top (maximum-fraction) group to emphasize coarse projection motifs. To quantify coarse co-variation between region groups, we computed a group-by-group Spearman rank correlation matrix across neurons using the row-normalized grouped profiles (pairwise complete observations) and visualized the resulting 12×12 correlation matrix.

For anatomical plots, per-cell summary annotations were generated from the grouped, row-normalized matrix: (i) the top projection group label and (ii) its corresponding maximum normalized value (“top projection strength”). The distribution of top-group assignments across cells was tabulated. Finally, top-group annotations were merged with BARseq-derived CCF coordinates from the combined metadata, and the merged table was exported for plotting.

### 4.5 Single-nucleus RNA sequencing (snRNAseq)

#### LC dissection

Twenty male and 20 female C57BL/6J mice (age P57–69) were used for two rounds of LC dissection. Half the mice received viral injections that aimed to label LC-NE neurons with YFP. There was no significant labeling, and analyses were aggregated across batches of mice. Each mouse was anesthetized with 2.5–3% isoflurane and transcardially perfused with oxygenated ACSFI (Calcium chloride, 0.5, D-Glucose, 25, HCl, 98, HEPES, 20, Magnesium sulfate, 10, Monosodium phosphate, 1.25, Myo inositol, 3, N-acetylcysteine, 12, N-methyl-d-glucamine, 96, Potassium chloride, 2.5, Sodium bicarbonate, 25, Sodium L-ascorbate, 5, Sodium pyruvate, 3, Taurine, 0.01, Thiourea, 2, in mM, 300-310 mOsm). Brains were immediately dissected, mounted and cut into 350-µm coronal slabs in oxygenated ACSFI with 5 µM anisomycin (A9789, Sigma) and 22.7 µM actinomycin-D (A1410, Sigma) on a Compresstome (Precisionary Instruments). LC with surrounding areas was dissected manually under a dissection microscope using local landmarks under bright field illumination. The dissected tissues were transferred to a 2 mL tube, immediately frozen in dry ice, and stored at –80 ◦C until all the samples in a cohort were collected.

#### Single-nucleus isolation

Nuclei were isolated using the RAISINs method with a few modifications as already described in a nuclei isolation protocol developed at Allen Institute for Brain Science. All reagents and methods are described in (doi:10.17504/protocols.io.4r3l22n5pl1y/v1). Briefly, dissectates were thawed and pooled separately in a 12-well plate containing CST extraction buffer. Tissues were chopped using spring scissors in ice-cold CST buffer for 10 min. The suspension was then transferred to a 50-mL conical tube while passing through a 100 µm filter and the walls of the tube were washed using ST buffer. Next, the suspension was gently transferred to a 15-mL conical tube and centrifuged in a swinging-bucket centrifuge for 5 min at 500 rcf and 4 °C. The supernatant was removed, and pellets were resuspended in 100 µL 0.1× lysis buffer and incubated for 2 min on ice. After addition of 1 mL wash buffer, samples were gently filtered using a 20 µm filter and centrifuged as above. After supernatant was discarded, pellets were resuspended chilled nuclei buffer.

#### Sequencing library preparation

For 10x snRNA-seq, the cell suspensions were processed using Chromium GEM-X Single Cell 3’ Kit v4 (1000691, 10x Genomics). We followed the manufacturer’s instructions for cell capture, barcoding, reverse transcription, cDNA amplification and library construction (doi:10.17504/protocols.io.261ger6njl47/v2).

#### Sequencing data processing and QC

Processing of 10x Genomics snRNA-seq libraries was performed as described previously (Yao et al. 2023). We targeted a sequencing depth of 120,000 reads per cell. In brief, libraries were sequenced on an Illumina NovaSeq system at the Northwest Genomics Center at the University of Washington, and sequencing reads were aligned to the mouse reference transcriptome (M21, GRCm38.p6) using the 10x Genomics CellRanger pipeline (v.8.0.1) with the default parameters. The cell by gene matrix along with metadata was exported as R Data (.rda).

#### Data import, pre-processing and quality control

Single-nucleus RNA-sequencing (snRNA-seq) data were processed in R using Seurat (Seurat v5.4.0; SeuratObject v5.3.0) with batch correction performed using Harmony. Raw UMI count matrices and accompanying metadata were used to generate a Seurat object. Nuclei with fewer than 2,000 or more than 15,000 detected genes were excluded, as were nuclei with >4% mitochondrial or >4% ribosomal transcript content. Genes with <3 cells detected were removed and cells with <200 detected genes were also excluded. Cells annotated with ≥0.4 doublet scores were removed. After quality control, 234,813 nuclei (from 398,912 initial profiles) were retained for downstream analyses. Data were counter-per-10k normalized followed by log-normalizations. Then highly variable genes were identified, and scaled expression values were used for principal component analysis (PCA). Batch effects across samples were corrected using Harmony on PCA embeddings, and the integrated low-dimensional space was used to construct a shared nearest-neighbor graph for community detection. Clusters were identified using graph-based clustering and visualized using UMAP computed on the Harmony-corrected embeddings.

#### Identification of NE cells

To classify NE cells, clusters with significant NE marker gene expression (including *Dbh, Th, Slc6a2*, and *Slc18a2*; Figure S8) were considered. Specifically, we calculated the average expression of each gene for each clusters, and only the cluster with larger than 1 unit of log CP10k expression for all the marker genes were kept. The remaining clusters were subset for further analysis. The LC subset was reprocessed independently, including normalization, identification of 2,000 variable features, scaling, PCA, and Harmony-based integration restricted to LC nuclei. A new neighbor graph, UMAP embedding, and graph-based clustering were computed to resolve LC subpopulations at higher resolution. Low-quality clusters were found by computing the median of the doublet score and total UMI counts, and clusters with the lowest median UMI (corresponding to the lowest detected genes) as well as the cluster with the highest median doublet score were excluded, and the final LC subcluster object was retained for downstream analyses, yielding 4,868 LC-NE cells. This subcluster was further examined for its quality, and we found 24 cells were outliers when re-clustered (detected in UMAP), corresponding to a population with high doublet scores (average 0.178 versus 0.070 for the rest of the populations). We removed these 24 cells, yielding a final number of 4,845 LC-NE cells.

#### LC-NE transcriptomic analysis

Code for snRNAseq, MERFISH, and retro-seq analyses are available in the accompanying GitHub repository and Code Ocean capsules. Analyses were done in Python using the Scanpy package. The selected cluster was saved as a separate AnnData object while retaining all genes. This dataset was used for downstream analyses. Gene chromosomal locations were queried using the mygene Python package (https://pypi.org/project/mygene/). Genes located on chromosomes X or Y were excluded. From the remaining genes, 1,500 HVGs were identified (Seurat v3 implementation in Scanpy).

#### Batch correction and normalization with scVI

We ran batch correction first using scVI (version 1.3.3, https://scvi-tools.org/). Batches were defined as combinations of experimental date (two categories) and sex (two categories), resulting in four batches. Gene expression was modeled using a zero-inflated negative binomial (ZINB) distribution. A single-layer variational autoencoder with four latent dimensions was used to account for the relatively homogeneous cell population. All other parameters were set to default values. Normalized, batch-corrected expression values (ρ) were extracted from the trained model, representing reconstructed gene expression under an assumed library size of 1 (or converted to counts per million, CPM, when specified), with batch effects averaged across all batches.

#### Scaling and dimensionality reduction

Prior to PCA, gene expression values were z-score normalized by scaling each gene to zero mean and unit variance. Extreme standardized values (z-score normalization with scanpy function ‘pp.scale’) were clipped to ±10. PCA was performed on the 1,500 HVGs, and the top 50 PCs were retained. UMAP embeddings were computed from the scVI latent space using default Scanpy parameters for visualization.

#### Pseudospace analysis

To capture continuous transcriptional variation that we observed in PCA space, we developed a metric termed “pseudospace,” representing an ordering of cells along the primary transcriptional axis.

##### 1. Initial cell ordering

Cells were embedded in 2D UMAP space derived from PCA. The UMAP coordinates were centered, and singular value decomposition (SVD) was applied. The first PC of the centered UMAP coordinates defines the primary axis of variation. Cells were projected onto this axis to obtain an initial ordering.

##### 2. Trajectory fitting in PCA space

Using the initial ordering, a smooth trajectory was fitted in 10-dimensional PCA space. Cells were ordered according to the initial ordering, and the first 10 PCs (explaining approximately 79.5% of variance) were extracted. For each PC dimension, a cubic polynomial was fitted as a function of the ordering parameter using least squares. This yielded a parametric trajectory *f* (*t*) = [*f*_1_(*t*), *f*_2_(*t*), …, *f*_10_(*t*)]. Here, subscripts indicate the PC dimensions and t denotes the continuous pseudotime (ordering) parameter. The fitted function *fk*(*t*)(*k* = 1, …10) defines a smooth curve embedded in 10-d space that approximates the underlying cell distribution.

##### 3. Pseudospace score calculations

The parametric trajectory was sampled at 10,000 evenly spaced points. Each cell was projected onto this trajectory by finding the value of t that minimized the squared Euclidean distance between the cell’s PC coordinates *x*_*i*_ and the sampled points (on the fitted trajectory), 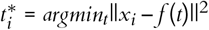. The closest trajectory point was identified, and the corresponding parameter value *t*∗ was assigned to the cell. These values were normalized to the range [0, 1] to produce final pseudospace scores.

#### Validating the robustness of pseudospace scores

Robustness was assessed using bootstrap analysis with 10,000 resamples. For each bootstrap sample, the trajectory was re-fitted and 95% confidence intervals were computed.

#### Identification of U-shaped genes

To identify genes exhibiting U-shaped (non-monotonic) expression patterns along the principal transcriptomic gradient we first fit the 1-d pseudospace scores as described above, where each cell was assigned a pseudospace score. The gene expression profiles were reordered according to the pseudospace score, and the first and second derivatives of each gene’s smoothed expression with respect to pseudospace were computed. Genes with abrupt expression changes were identified by applying dual thresholds on maximum absolute first derivative (>15) and maximum absolute second derivative (>100). From this filtered set, U-shaped genes were further selected based on the criterion that the first derivative at the beginning of the trajectory was negative (<–4) and at the end was positive (>4), indicating a decrease-then-increase (namely U-shaped) expression profiles. Candidate U-genes were ranked by their peak-to-peak expression range (i.e., difference between the maximum and minimum).

#### MMIDAS for identifying the potential clusters

MMIDAS, an unsupervised clustering framework based on coupled autoencoders with GAN-based augmentations (Marghi et al. 2024), was applied as an independent validation approach. A two-arm variational autoencoder architecture was used with 10 latent dimensions, including 4 continuous dimensions and 15 categorical variables. All other parameters followed the implementation provided in the code repository.

#### Cluster quality analysis

Clustering quality was evaluated using silhouette scores across *K* = 1, …, 15 clusters. Analyses were performed on 10-dimensional PCA and scVI representations. Three types of data are used: (1) synthetic data with known ground truth (*K* = 2); (2) synthetic data generated from extreme pseudospace endpoints using a mixture of Gaussian distributions (with *K* = 2); (3) the LC-NE data. For the LC-NE data, we used two methods for the clustering: (a) Gaussian mixture models applied to PCA space; and (b) MMIDAS clustering evaluated in PCA space. For (1) and (2) data we used the same approach as (a). For each method and K, five independent runs were performed with different random seeds, and mean ± SD silhouette scores were reported. Higher silhouette scores (ranging from –1 to 1) indicate well-separated clusters, and they were always highest with a single cluster. As shown in Figure S4, we observed a large decrease when k increased from 1 to 2 regardless of which method was used, indicating that one cluster best described the population.

### 4.6 Analysis of multiplexed error-robust fluorescence in situ hybridization (MERFISH) data

#### Data source and image registrations

MERFISH datasets were downloaded from the original publication repositories (Nardone et al. 2024). We analyzed data from 4 mice (2 female) from the original dataset, which gave 24 slices in total. The original data contains the two-dimensional coordinate and the gene counts for each cell. For each image slice, we ran leiden clustering and colored each cell based on the assigned clusters, before performing image registration manually using QuickNII (https://www.nitrc.org/projects/quicknii) for affine transformations and VisuAlign (https://www.nitrc.org/projects/visualign/) for deformable transformations, to map to CCF coordinates. Transformation matrices were inverted and applied to images, which then yielded the three dimensional spatial coordinates for each cell, which were aligned to anatomical reference frames.

#### Data preprocessing and LC-NE cell selection

MERFISH gene expression matrices and CCF-aligned cellular coordinates were processed independently for each tissue section. For each section, we first removed blank probes and then computed the product of raw counts for the canonical LC-NE markers *Dbh, Th* and *Slc6a2* for every cell, retaining only cells with a marker-product value >10. The filtered cells from all sections were concatenated into a single AnnData object with raw counts preserved and associated metadata (slicename and CCF coordinates). After concatenation, we applied a second filter requiring non-zero expression of all three markers, yielding a high-confidence LC-NE population. To further restrict the dataset to anatomically consistent LC neurons, cells with dorsal-ventral (CCF y-axis) coordinates greater than 220 pixels (5.5 mm) were excluded. The resulting high-confidence LC-NE population was visualized in three-dimensional CCF space to confirm spatial consistency with the LC template.

#### Batch correction, dimensionality reduction, and clustering

The LC-NE MERFISH AnnData object was used as input to scVI, with tissue slice section specified as the batch covariate. A variational autoencoder (one hidden layer, four latent dimensions) was trained under a negative binomial likelihood to jointly perform library size normalization, denoising, and batch correction. The resulting latent representations were used to construct a 30-nearest-neighbor graph, followed by UMAP embedding for visualization and Leiden community detection (resolution = 0.5) to identify transcriptionally distinct subpopulations. Batch-corrected normalized expression values were obtained from the posterior mean for downstream interpretation. For all downstream analysis, library-normalized raw counts were used unless specified to use the batch-corrected counts.

#### Canonical correlation analysis (CCA) for space-gene relationships

CCA was performed between gene expression matrices and 3D spatial coordinates to identify space-gene relationships within the LC. Because the LC is approximately bilaterally symmetric, spatial coordinates were folded across the midline to remove left-right redundancy: coordinates from the right hemisphere were reflected across the midline and mapped onto the left hemisphere. Both gene expression and spatial features were then standardized (zero mean, unit variance) to remove scale differences across features. The top two canonical components were retained. The canonical variables (scalar) were then used as the cell weight for the visualization, termed “weighted gene score.”

#### Gene-space correlation analysis

Rank correlations were computed between individual gene expression levels and projection scores along identified spatial directions. Genes were ranked by absolute correlation values. The top 20 genes were listed in the table and visualized in CCF space.

#### Gene expression imputation

To expand gene coverage beyond the MERFISH panel and enable spatial visualization of genes not directly measured, we performed k-nearest neighbors (kNN) imputation using matched snRNA-seq data as reference. Highly variable genes were identified in the snRNA-seq dataset, and the union of these genes with the MERFISH panel defined the target gene space for imputation. For similarity computation, both datasets were restricted to their shared genes. Expression profiles were normalized using a row-wise rank transformation followed by standardization to improve cross-platform comparability.

For each MERFISH cell, we identified its *k* nearest neighbors (*k* = 200) in the snRNA-seq dataset using Euclidean distance in the normalized expression space. Gene expression for unmeasured genes was imputed as a weighted average of the corresponding snRNA-seq neighbor profiles across the full union gene set. Neighbor weights were derived from a distance-based softmax kernel, *w*_*ij*_ ∝ exp(–*d*_*ij*_)/τ, and normalized to sum to one for each MERFISH cell.

#### Choosing the number of neighbors for gene imputation

To determine the number of neighbors for the imputations, we took 5 genes that were included in the original MERFISH gene panel and we know *a priori* to have large variance, and imputed their expressions from the rest of the genes. We varied the number of neighbors and tested how many neighbors would give the optimal correlations between imputed and the ground-truth expression values. Based on these results, we chose 200 neighbors for the following gene imputations.

#### Pseudospace score imputations

Similar to gene expression imputation, the pseudospace scores derived from snRNAseq analysis were transferred to MERFISH cells (with *k* = 100). For each MERFISH cell, we computed the weighted average pseudospace score from *k* nearest snRNAseq neighbors using the shared genes. To quantify uncertainty, we measured the weighted SD of the neighbors’ pseudospace scores, as well as the confidence intervals. For the confidence intervals, we used weighted sampling of 500 samples among the neighbors of each cell, based on the pre-calculated weights (which sum to one across all the neighbors) to get the re-sampled means. We then took the 2.5th and 97.5th percentiles. The width of the confidence interval (which contains 95% of the means, and the value is constrained between 0 and 1) was computed. The larger width indicates larger uncertainty.

To quantify mapping reliability, we estimated a baseline cross-modal distance distribution from randomly paired cells and compared each MERFISH cell’s mean neighbor distance to this baseline. We derived per-cell confidence metrics, including a normalized distance score and an empirical p-score (fraction of random distances exceeding the observed mean neighbor distance), which were retained for downstream quality control of imputed profiles.

### 4.7 Retrograde snRNAseq (retro-seq)

Experimental mice were generated by crossing homozygous RCFL-H2B-GFP mice (He et al. 2016) (Jackson Laboratory, 028581) to heterozygous *Dbh*-Cre mice. The resulting offspring heterozygous for both *Dbh*-Cre and RCFL-H2B-GFP alleles were used for retrograde injections and subsequent SSV4 snRNA sequencing experiments following FACS sorting for GFP.

*Dbh*-Cre; RCFL-H2B-GFP mice (P40–P50) were anesthetized using oxygenated 4% isoflurane and maintained with oxygenated 0.7–1.0% isoflurane throughout the surgery. Animals were mounted in a stereotaxic frame (David Kopf Instruments), and small craniotomies were performed in the skull above the injection target (dx.doi.org/10.17504/protocols.io.14egn8ewzg5d/v8 for pressure injections using NanoInject III). Spinal cord injections were performed between C4–C5 vertebrae without drilling (dx.doi.org/10.17504/protocols.io.yxmvm8mq9g3p/v1 using NanoInject III). Intracerebral injections were always targeted to the right hemisphere. Spinal cord injections were performed bilaterally. Table 2 lists injection targets, types, volumes and AP/ML/DV coordinates from Bregma. Virus AAV pEF1a-DIO-FLPo-WPRE-hGHpA was procured from Addgene (87306-AAVrg, lot v188462).

**Table 2:**
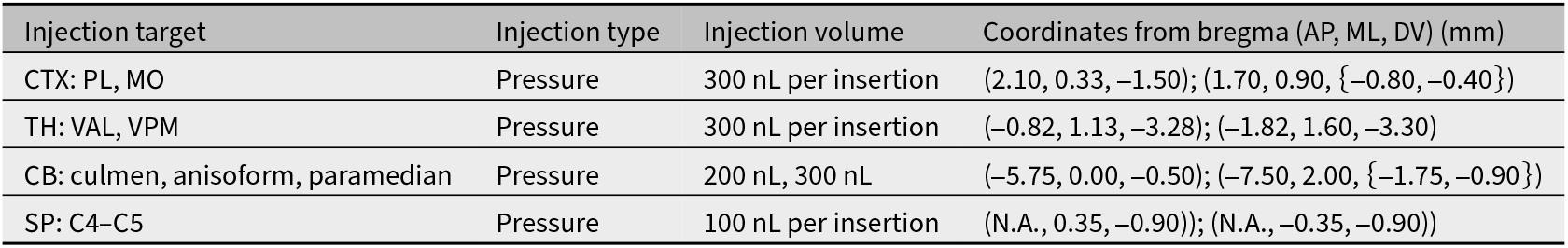
Virus injections for retro-seq experiments. Summary of animal and cell counts by projection target and labelled soma location. Spinal cord injections were made bilaterally. All other injections were in the right hemisphere. Injection coordinates are relative to bregma (anterior-posterior, AP; medial-lateral, ML; dorsal-ventral, DV).

#### Tissue harvest and processing

Mice were euthanized with an isoflurane overdose 4–5 weeks after AAV injection and transcardially perfused with cold, pH 7.4 HEPES buffer containing 110 mM NaCl, 10 mM HEPES, 25 mM glucose, 75 mM sucrose, 7.5 mM MgCl_2_, and 2.5 mM KCl to remove blood from brain. Brains were quickly dissected out and flash frozen in liquid nitrogen. Frozen brains were placed into a 5 mL centrifuge tube containing 1 mL frozen OCT cushion at the bottom and capped.

#### Data preprocessing and quality control

Retro-seq data were filtered to retain only genes present in the reference snRNAseq dataset, resulting in shared gene coverage for integration analysis. Projection site categories were cleaned and organized into three main projection targets: frontal cortex, cerebellum, and spinal cord. Thalamic projecting neurons were kept but the projection information was excluded from downstream analysis to avoid potential confusion with the frontal-cortex-projecting neurons.

#### Gene length normalization (TPM normalization)

SmartSeq2 technology is known to exhibit gene length-dependent bias; therefore, expression values for all samples were normalized by the corresponding gene lengths. Gene lengths were derived from a reference GTF annotation file from GENCODE (version M38, https://www.gencodegenes.org/mouse/) by summing exon lengths per gene. Exons lacking a ‘gene_name’ annotation in the GTF file were excluded. For each gene, exon lengths were summed across all annotated exons to obtain the total exonic gene length in base pairs.

Gene length-normalized expression values were computed by dividing raw transcript counts by the corresponding gene lengths. Counts for each sample were scaled to CPM, then rounded to integers, and used for all downstream analyses.

#### Assessment of batch effects using supervised classification

Residual batch effects were assessed using supervised classification. Following TPM normalization, log transformation, scaling, and PCA, random forest classifiers were trained to predict donor identity, sex, or projection target from gene expression profiles. Model performance was evaluated using 5-fold stratified cross-validation and summarized as mean accuracy ± SD Classification accuracy was compared to the theoretical chance level (1/K where K being number of classes), with above-chance performance indicating detectable batch or covariate structure in the data.

#### Dimensionality reduction and batch correction

As in the snRNAseq analysis, scVI was applied for dimensionality reduction and batch correction across experimental batches. The scVI model was configured with (1) batch key defined as external donor name; (2) a single-layer architecture with four latent dimensions; and (3) ZINB gene likelihood. The other parameters were set to be default. The trained scVI model produced four-dimensional latent representations, which were used for downstream analyses, including UMAP embedding. The batch removal was confirmed by manual inspections.

#### Projection site predictions from gene expression

Projection sites were predicted using supervised classification. The dataset was split into training (60%), validation (20%), and test (20%) sets using stratified sampling to preserve class proportions. Five algorithms were evaluated: logistic regression, random forest, support vector machine, k-nearest neighbors, and gradient boosting. Hyperparameter optimization was performed using 5-fold cross-validation on the training set. Models were first evaluated on the validation set, then retrained on the combined training and validation set, and finally assessed on the test set. Classification performance was quantified using accuracy.

#### Projection site imputations for snRNAseq data

As in the MERFISH analysis, projection site labels were transferred from retro-seq data to snRNAseq data using k-nearest neighbors based on rank-based correlations. For each snRNAseq cell, the 10 nearest neighbors in the retroseq dataset were identified. A projection target was assigned if at least 8 of the 10 neighbors agreed on the projection site. The quality of imputation was computed as disagreement score which is 1-(number of agreement)/(number of valid projections). (Invalid projections indicate thalamus-projecting neurons which is not considered in this case.)

### 4.8 Patch-seq

#### Transgenic mice and labeling projection neurons

Heterozygous *Dbh*-Cre mice crossed with homozygous Ai65 mice (P28–P30) were anesthetized with 2.0% isoflurane in O_2_ and maintained at 0.7–1.0% isoflurane throughout surgery. Mice were mounted in a stereotaxic frame (David Kopf Instruments) for injections of AAV pEF1a-DIO-FLPo-WPRE-hGHpA (Addgene 87306-AAVrg, lot v188462). Small craniotomies were made in the skull above the injection target for intracerebral injections (dx.doi.org/10.17504/protocols.io.bp2l6nr7kgqe/v8). Spinal cord injections were made between C4–C5 vertebrae without drilling (dx.doi.org/10.17504/protocols.io.yxmvm8mq9g3p/v1). Intracerebral injections were in the right hemisphere. Spinal cord injections were bilateral. Mice recovered in their home cage for approximately 21 days before Patch-seq experiments.

#### Tissue processing

For preparation of acute brain slices, adult male and female mice (ages P50–P74) were first fully anesthetized by 5% isoflurane inhalation. A transcardial perfusion was then performed with ∼ 10 mL of ice-cold cutting artificial cerebrospinal fluid (ACSF; 0.5 mM calcium chloride (dehydrate), 25 mM D-glucose, 20 mM HEPES buffer, 10 mM magnesium sulfate, 1.25 mM sodium phosphate monobasic monohydrate, 3 mM myo-inositol, 12 mM N-acetyl-L-cysteine, 96 mM N-methyl-D-glucamine chloride (NMDG-Cl), 2.5 mM potassium chloride, 25 mM sodium bicarbonate, 5 mM sodium L-ascorbate, 3 mM sodium pyruvate, 0.01 mM taurine, and 2 mM thiourea (pH 7.3), which had been continuously bubbling with a mixture of 95%O_2_/5%CO_2_). 350 µm coronal or sagittal sections containing the LC were sliced on a vibrating microtome (VT1200S Vibratome, Leica Biosystems). immediately after slicing, brain slices were placed in warm (34 ◦C) oxygenated cutting ACSF for 10 min, then allowed to further recover in holding ACSF (2 mM calcium chloride (dehydrate), 25 mM D-glucose, 20 mM HEPES buffer, 2 mM magnesium sulfate, 1.25 mM sodium phosphate monobasic monohydrate, 3 mM myo-inositol, 12.3 mM N-acetyl-L-cysteine, 84 mM sodium chloride, 2.5 mM potassium chloride, 25 mM sodium bicarbonate, 5 mM sodium L-ascorbate, 3 mM sodium pyruvate, 0.01 mM taurine, and 2 mM thiourea (pH 7.3)), bubbling with a mixture of 95% O_2_/5% CO_2_ at room temperature until transferred to the microscope for recordings.

#### Patch-seq recordings

Slices were bathed in warm (34 ◦C) recording ACSF (2 mM calcium chloride (dehydrate), 12.5 mM D-glucose, 1 mM magnesium sulfate, 1.25 mM sodium phosphate monobasic monohydrate, 2.5 mM potassium chloride, 26 mM sodium bicarbonate, and 126 mM sodium chloride (pH 7.3)) and continuously bubbled with 95% O_2_/5% CO_2_. The bath solution contained blockers of fast glutamatergic (1 mM kynurenic acid) and GABAergic synaptic transmission (0.1 mM picrotoxin). Thick-walled borosilicate glass (Warner Instruments, G150F-3) electrodes were manufactured (Narishige PC-10) with a resistance of 4–5 MΩ. Before recording, the electrodes were filled with approximately 1.0 to 1.5 µL of internal solution with biocytin (110 mM potassium gluconate, 10.0 mM HEPES, 0.2 mM ethylene glycol-bis (2-aminoethylether)-N,N,N,N-tetraacetic acid, 4 mM potassium chloride, 0.3 mM guanosine 5-triphosphate sodium salt hydrate, 10 mM phosphocreatine disodium salt hydrate, 1 mM adenosine 5-triphosphate magnesium salt, 20 µg/mL glycogen, 0.5U RNAse inhibitor (Takara, 2313A) and 0.5% biocytin (Sigma B4261), pH 7.3). The pipette was mounted on a Multiclamp 700B amplifier headstage (Molecular Devices) fixed to a micromanipulator (PatchStar, Scientifica). Electrophysiology signals were recorded using an ITC-18 Data Acquisition Interface (HEKA). Commands were generated, signals processed, and amplifier metadata were acquired using MIES (https://github.com/AllenInstitute/MIES), written in Igor Pro (Wavemetrics). Data were filtered (Bessel) at 10 kHz and digitized at 50 kHz. Data were reported uncorrected for the measured liquid junction potential of –14 mV between the electrode and bath solutions. Prior to data collection, all surfaces, equipment and materials were thoroughly cleaned in the following manner: a wipe down with DNA away (Thermo Scientific), RNAse Zap (Sigma-Aldrich), and finally with nuclease-free water. After formation of a stable seal and break-in, the resting membrane potential of the neuron was recorded (typically within the first minute). A bias current was injected, either manually or automatically using algorithms within the MIES data acquisition package, for the remainder of the experiment to maintain that initial resting membrane potential. Bias currents remained stable for a minimum of 1 s before each stimulus current injection. To be included in the analysis, neurons needed to have a >1 GΩ seal recorded before break-in and an initial access resistance <20 MΩ and <15% of the *R*_input_. For an individual sweep to be included, the following criteria were applied: (1) the bridge balance was <20 MΩ and <15% of *R*_input_; (2) bias (leak) current within ± 100 pA; and (3) root mean square noise measurements in a short window (1.5 ms, to gauge high frequency noise) and longer window (500 ms, to measure patch instability) <0.07 mV and <0.5 mV, respectively. After electrophysiological recording, the pipette was centered on the soma or placed near the nucleus (if visible). A small amount of negative pressure was applied (approximately –30 mbar) to begin cytosol extraction and to attract the nucleus to the tip of pipette. After approximately one minute, the soma visibly shrank and/or the nucleus was near the tip of the pipette. While maintaining negative pressure, the pipette was slowly retracted; slow, continuous movement was maintained while monitoring the pipette seal. Once the pipette seal reached > 1GΩ and the nucleus was visible on the tip of the pipette, the speed was increased to remove the pipette from the slice. The pipette containing internal solution, cytosol, and the nucleus was removed from pipette holder, and its contents were expelled into a PCR tube containing the lysis buffer (Takara, 634894). Electrophysiological features were computed using the Intrinsic Physiology Feature Extractor (IPFX) Python package.

#### cDNA and sequencing

For Patch-seq experiments, we reverse transcribed the collected nuclear and cytosolic mRNA, and sequenced the resulting cDNA using a the SMART-Seq v4 method (Tasic et al. 2018; Lee et al. 2021). We used the SMART-Seq v4 Ultra Low Input RNA Kit for Sequencing (Takara, 634894) to reverse transcribe poly(A) RNA and amplify full-length cDNA according to the manufacturer’s instructions. We performed reverse transcription and cDNA amplification for 21 PCR cycles in 0.65 mL tubes, in sets of 88 tubes at a time. At least 1 control 8-strip was used per amplification set, which contained 4 wells without cells and 4 wells with 10 pg control RNA. Control RNA was either Mouse Whole Brain Total RNA (Zyagen, MR-201) or control RNA provided in the SMARTSeq v4 kit. All samples proceeded through Nextera XT DNA Library Preparation (Illumina FC-131-1096) using either Nextera XT Index Kit V2 Set A-D (FC-131-2001,2002,2003,2004) or custom dual-indexes provided by IDT (Integrated DNA Technologies). Nextera XT DNA Library prep was performed according to manufacturer’s instructions except that the volumes of all reagents including cDNA input were decreased either to 0.4x or to 0.2x by volume. Each sample was sequenced to approximately 500,000 – 1 million reads.

Fifty-base-pair paired-end reads were aligned to mm10 GENCODE vM23/Ensembl 98 reference genome, downloaded from 10X cell ranger (refdata-cellranger-arcmm10-2020-A-2.0.0). Sequence alignment was performed using STAR aligner (v2.7.1a) with default settings. PCR duplicates were masked and removed using STAR option “bamRemoveDuplicates.” Only uniquely aligned reads were used for gene quantification. Gene counts were computed using the R Genomic Alignments package (Lawrence et al. 2013) summarizeOverlaps function using “IntersectionNotEmpty” mode for exonic and intronic regions separately. Exonic and intronic reads were added together to calculate total gene counts; this was done for both the reference dissociated cell data set and the Patch-seq data set. Data were analyzed as counts per million reads (CPM).

Slices from Patch-seq experiments were mounted on slides and imaged on an upright AxioImager Z2 microscope (Zeiss, Germany) with an Axiocam 506 monochrome camera and 0.63x optivar lens. Two-dimensional tiled overview images were also captured (Zeiss Plan-NEOFLUAR 20X/0.5) in brightfield transmission and fluorescence channels. Light was transmitted using an oil-immersion condenser (1.4 NA). High-resolution, multi-tile image stacks were captured (Zeiss Plan-Apochromat 63x/1.4 Oil or Zeiss LD LCI Plan-Apochromat 63x/1.2 Imm Corr) at an interval of 0.28 µm (1.4 NA objective) or 0.44 µm (1.2 NA objective) along the Z axis. Image tiles were stitched in ZEN software and exported as single-plane TIFF files.

### 4.9 Physiology and behavior

#### Mice and surgery

We used heterozygous *Dbh*-Cre mice for all *in vivo* experiments. Forty-two mice (6 female) were used for behavioral experiments, 28 mice (5 female) were used for electrophysiological recordings during the behavioral task, and 14 mice (1 female) for calcium indicator recordings. Surgery was performed on mice between the ages of 8–16 weeks, under isoflurane anesthesia (1.5–2.0% in O_2_) and under aseptic conditions. During all surgeries, titanium headplates were surgically attached to the skull using dental adhesive (C&B-Metabond, Parkell). After the surgeries, analgesia (ketoprofen, 5 mg kg^−1^ and buprenorphine, 0.05–0.1 mg kg^−1^) was administered to minimize pain and aid recovery. Surgeries performed at the Allen Institute followed this protocol: dx.doi.org/10.17504/protocols.io.eq2lyj72elx9/v1.

For all experiments, mice were given at least one week to recover prior to water restriction. During water restriction, mice had free access to food and were monitored daily in order to maintain 80% of their baseline body weight. Mice were housed in a reverse light cycle (12h dark/12h light, dark from 08:00-20:00) and all experiments were conducted during the dark cycle between 12:00 and 20:00.

#### Behavioral task

Before training on the tasks, water-restricted mice were habituated to head fixation for 1–3 d with free access to water from the two spouts (21 ga stainless steel tubes separated by 4 mm) placed in front of the 38.1-mm diameter acrylic tube in which the mice rested. For LC-NE recordings with tetrodes and calcium indicator recordings in LC axons, the spouts were mounted on a micromanipulator (DT12XYZ, Thorlabs) with a custom digital rotary encoder to measure the position of the lick spouts with 5–10 µm resolution. Each spout was attached to a solenoid (ROB-11015, Sparkfun) to enable movement in the anterior-posterior axis of the mouse. The tones used for the cues (randomly assigned to go and no-go cues per mouse) were 7.5 and 15 kHz square waves generated by microcontrollers (ATmega16U2 or ATmega328), amplified and delivered through speakers (CUI Devices, GF0401M).

Licks were detected using custom circuits (Janelia Research Campus 2019-053). Task events were controlled and recorded using custom code (Arduino or Bonsai: https://github.com/AllenNeuralDynamics/dynamic-foraging-task) written for microcontrollers (ATmega16U2, ATmega328, or Harp boards). Water rewards were 2-3 µL, adjusted for each mouse to maximize the number of trials completed per session and to keep sessions around 60 min. Solenoids (LHDA1233115H or LHDB1233518H, The Lee Co) were calibrated to release the desired volume of water and were mounted on the outside of the sound-attenuated chamber used for behavior. For mice with tetrodes, white noise (2–60 kHz, Sweetwater Lynx L22 sound card, Rotel RB-930AX two-channel power amplifier, and Pettersson L60 Ultrasound Speaker), was played inside the chamber to mask ambient noise.

For LC-NE recordings with Neuropixels probes, behavior was controlled by Harp devices using Python and Bonsai (https://github.com/AllenNeuralDynamics/dynamic-foraging-task). Lick spouts were mounted on a New Scale microcontroller (M3-LS-3.4-15) to track and control lick spout locations across days. The tone used for the cue was 7.5 kHz at 70 dB, generated by a Harp Soundcard. Licks were detected using a custom Harp device (Lickety-split, https://github.com/AllenNeuralDynamics/harp.device.lickety-split).

Water rewards were 2-3 µL, adjusted for each mouse to maximize the number of trials completed per session and to keep sessions around 60 min. Solenoids (LHDA1233115H or LHDB1233518H, The Lee Co) were calibrated to release the desired volume of water and were mounted on the outside of the sound-attenuated chamber used for behavior.

During the 1–3 days of habituation, mice were trained to lick both spouts to receive water. Water was delivered following a lick to the correct spout at any time. Reward probabilities were chosen from the set {0, 1} and reversed every 20 trials. In the second stage of training (5–12 d), the trial structure with tone presentation was introduced. Each trial began with the 0.5 s delivery of either an auditory “go cue” (*P* = 0.95) or a “no-go cue” (*P* = 0.05) in tetrode and GCaMP experiments. Following the go cue, mice could lick either the left or the right spout. If a lick was made during a 1.5 or 1.8 s response window (in separate experiments), reward was delivered probabilistically from the chosen spout. In tetrode and GCaMP experiments, the unchosen spout was retracted when the tongue contacted the chosen spout to prevent mice from sampling both spouts within a trial. The unchosen spout was replaced 3.1 s after cue onset (or 3.0 s in experiments with axonal GCaMP). Following a no-go cue, lick responses were neither rewarded nor punished. Reward probabilities during this stage were chosen from the set {0.1, 0.9} and reversed every 20–35 trials. During this period of training only, water was occasionally manually delivered to encourage learning of the response window and appropriate switching behavior. During this second stage of training, we introduced a 100-ms delay between choice and outcome. This delay was gradually increased to 300 ms for electrophysiological experiments and 200 ms for axonal GCaMP. If a directional lick bias was observed in one session, the lick spouts were moved horizontally 50–300 µm in the opposite direction prior to the following session.

After the 3.0 or 3.1 s trial duration, inter-trial intervals (the times between consecutive cue onsets) were generated as draws from an exponential distribution with a rate parameter of 0.3 and a maximum of 20 s. This distribution results in a flat hazard rate for inter-trial intervals such that the probability of the next trial did not increase over the duration of the inter-trial interval. Inter-trial intervals were 4.83 s on average (range, 1.1–20 s, including a no-lick window; see below). In more than 90% of all well-trained sessions, mice made a leftward or rightward choice in greater than 90% of trials.

In the final stage of the task, the reward probabilities assigned to each lick spout were drawn pseudorandomly from the set {0.1, 0.5, 0.9}. The probabilities were assigned to each spout individually with block lengths drawn from a uniform distribution of 20–35 trials. To stagger the blocks of probability assignment for each spout, the block length for one spout in the first block of each session was drawn from a uniform distribution of 6–21 trials. For each spout, probability assignments could not be repeated across consecutive blocks. To maintain task engagement, reward probabilities of 0.1 could not be simultaneously assigned to both spouts. If one spout was assigned a reward probability greater than or equal to the reward probability of the other spout for 3 consecutive blocks, the probability of that spout was set to 0.1 to encourage switching behavior and limit the creation of a direction bias. If a mouse perseverated on a spout with reward probability of 0.1 for 4 consecutive trials, 4 trials were added to the length of both blocks. This prevented mice from choosing one spout until the reward probability became high again.

To minimize spontaneous licking, we enforced a 1 s no-lick window prior to tone onset. Licks within this window were punished with a new randomly-generated inter-trial interval, followed by a 1 s no-lick window. Implementing this window significantly reduced spontaneous licking throughout the entirety of behavioral experiments.

#### Electrophysiology: viral injections

To express ChR2 (Boyden et al. 2005), Chrimson (Klapoetke et al. 2014), or ChRmine (Marshel et al. 2019) in LC-NE neurons, we pressure-injected rAAV5-EF1a-DIO-hChR2(H134R)-EYFP (3 × 10^13^ GC mL^−1^), AAV5-Syn-FLEX-rc[ChrimsonR-tdTomato] (2.2 × 10^13^ GC ml^−1^), or pAAV-Ef1a-DIO-ChRmine-eYFP-WPRE (9.69 × 10^12^ GC ml^−1^, Addgene; packaged in house, lot number VT8453G) into the LC of *Dbh*-Cre mice at a rate of 1 nL s^−1^ (MMO-220A, Narishige). pAAV-EF1a-double floxed-hChR2(H134R)-EYFP-WPRE-HGHpA was a gift from Karl Deisseroth (Addgene viral prep 20298-AAV5; RRID:Addgene_20298). pAAV-Syn-FLEX-rc[ChrimsonR-tdTomato] was a gift from Edward Boyden (Addgene plasmid 62723; http://n2t.net/addgene:62723; RRID:Addgene_62723). pAAV-Ef1a-DIO ChRmine-eYFP-WPRE was a gift from Karl Deisseroth (Addgene plasmid 130996; http://n2t.net/addgene:130996; RRID:Addgene_130996).

For ChR2, we made three injections of 200 nL at the following coordinates: 0.28–0.35 mm anterior to the junction of the inferior colliculus and cerebellum, 0.85–0.90 mm right from the midline, and {2.95, 3.15, 3.35} mm ventral from the pial surface. Before the first injection, the pipette was left at the most ventral coordinate for 5 min. Before each injection, the pipette was withdrawn 50 µm and left in place for 5 min after injection. For electrophysiology experiments with rAAV5-EF1a-DIO-hChR2(H134R)-EYFP injections, the microdrive was implanted through the same craniotomy.

For Chrimson and ChRmine, we injected 300 nL at the following coordinates: from bregma, 5.4 mm posterior for male mice, 5.2 mm posterior for female mice; 0.85 mm lateral; and (2.9, 3.1) ventral from surface. Details can be found here: dx.doi.org/10.17504/protocols.io.eq2lyj72elx9/v1.

#### Electrode recording

We used two types of electrodes for extracellular recordings: tetrodes and Neuropixels probes. For electrophysiological experiments with tetrodes, we implanted a custom microdrive targeting right LC, entering through a craniotomy at 0.28–0.35 mm anterior to the junction of the inferior colliculus and cerebellum and 0.85–0.90 mm lateral from the midline (identified using the vasculature as a landmark). For recordings with tetrodes, we sampled at 32 kHz (Digital Lynx 4SX, Neuralynx, Inc.). The recording system was connected to 8 implanted tetrodes (32 channels, nichrome wire, PX000004, Sandvik) fed through guide tubes that could be advanced with the turn of a screw on a custom, 3D-printed microdrive. The impedances of each wire in the tetrodes were reduced to 180–250 kΩ by gold plating. The tetrodes were adhered to a 200 µm optic fiber used for optogenetic identification. After each recording session, the tetrode-optic-fiber bundle was driven down 30–75 µm. Channels were bandpass-filtered between 0.3–6 kHz. The bandpass-filtered signal, *x*, was thresholded at 2.5σ_*n*_ where 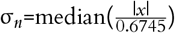. Detected peaks were sorted into individual unit clusters offline (Spikesort 3D, Neuralynx Inc.) using peak waveform amplitude, minimum waveform trough, and waveform PCA. We used two metrics of isolation quality as inclusion criteria: L-ratio (<0.05) and interspike interval (ISI) violation ratio <0.1%.

For electrophysiological experiments with Neuropixels probes, we made acute craniotomies at 0.28–0.35 mm anterior to the junction of the inferior colliculus and cerebellum and 0.85–0.90 mm lateral from the midline prior to recording (details at https://www.protocols.io/view/acute-lc-craniotomy-and-skull-thinning-for-neuropi-eq2ly6o6qgx9/v1) and inserted the probe either vertically or with a 6-degree anterior angle. A Kilosort pipeline (Buccino et al. 2026) was used for spike detection and sorting into single units. Signals adjacent to laser stimulation (0–4 ms) were removed and smoothed. Signals were filtered between 0.3–6 kHz, with phase shift correction and common reference removal. Probe displacement was estimated using the DREDge algorithm (Windolf et al. 2025). For some sessions, we also made a second recording at the brain surface without moving the probe to estimate the brain’s surface location along the probe.

The following were applied as initial inclusion criteria for single units unless further specified: ISI-violation ratio <0.1 (unless identified to project to cortex, where we used 0.2 as a threshold); presence ratio >0.8, amplitude cut off <0.1. Unit drift detection was performed by checking for timepoints where abrupt spike rate changes could be predicted by changes of probe location (estimated by DREDge) and changes of waveform peaks. Only periods without abrupt changes were used for future analysis. See details in the “Drift control” section below.

#### Spike waveform extraction

To minimize distortion of extracellular spike waveforms by filtering, raw traces were Butterworth filtered between 50–8,000 Hz before computing spike waveforms. In waveform analysis, only neurons recorded with tetrodes or Neuropixels 2.0 were included. To optimize the data for unit waveform characterization, unit inclusion criteria were modified as follows: ISI violation ratio bar was raised to 0.5, and a peak threshold of lower than –50 µV was applied. Each spike waveform was aligned to the time of its peak and normalized so that baseline started at zero and had peak values of 1. Six features were used to characterize each waveform: post-peak trough, post-peak time, post-slope, pre-slope, slope symmetry, trough distance, and trough symmetry (Figure S14a). All features were z-scored before further analysis.

#### Optical identification of LC-NE neurons

In tetrode recordings, light was delivered through a fiber attached to the tetrode at 5 or 10 Hz with pulse width of 10 or 20 ms. Laser irradiance was 0.2 mW, measured from the patch cord. In Neuropixels 2.0 recordings, light was delivered from brain surface above the inferior colliculus at 5 Hz with pulse width 4 or 5 ms. Laser irradiance was 10, 20, 30, 40, or 50 mW, measured at the collimator that focused the laser onto the brain surface.

Individual neurons were determined to be identified LC-NE neurons using the following criteria: (1) Responses to brief pulses (5, 10 or 20 ms) of laser stimulation (473, 560 or 640 nm wavelength, depending on the opsin) with significant increase of firing rate compared to baseline on both early and late pulses in a train, with short latency (<20 ms but >2 ms) and with higher probability than sham (>0.55; Figure S13f ). (2) Light-evoked spike waveforms were similar to spontaneous waveforms, with correlation coefficient greater than 0.95 and Euclidean distance (normalized to spike amplitude) less than 0.35.

In addition to observing responses to light stimuli, in tetrode experiments, LC targeting was confirmed by performing electrolytic lesions of the tissue (25 s of 20 µA direct current across two wires of the same tetrode) and examining the tissue after perfusion; in Neuropixels recordings, probe tracks were reconstructed (details below).

#### Antidromic stimulation

To identify if an LC-NE neuron projected to cortex, light was delivered above multiple sites in frontal cortex where the skull was thinned and cleared using cyanoacrylate, covered by Kwik-Cast or Kwik-Sil to avoid damage. Light (640 nm) was delivered at 5 Hz with a pulse width of 4 ms.

For each neuron, we defined “antidromic response time” τ_anti_ as the peak time of the peristimulus time histogram (PSTH) aligned to antidromic stimulation; “response jitter” as the half-peak-width of the PSTH around τ_anti_; “collision window” as the time window τ_anti_ before and τ_anti_ after antidromic stimulation (Figure S13j). The collision window is the time during which if a spontaneous spike occurred, it would collide with the backpropagating antidromic spike.

We fit a regression to the spike rate λ_resp_ around τ_anti_ after real or sham (equal number of random timepoints as real) antidromic stimulations. Regressors were as follows: sham or real laser stimulation *L* ∈ (0, 1) (0 for sham), spike rate in the collision window λ_col_, interaction between the presence of laser stimulation and presence of spikes in the collision window, such that λ_resp_ ∼ 1 + *L* + λ_col_ + *L* × λ_col_. A neuron was identified as projecting to cortex if it responded to laser stimulation consistently (*t*-statistic of laser stimulation *L* > 0 and *p* < 0.005) unless there were spontaneous spikes in the collision window (*t*-statistics of interaction between laser stimulation *L* and spike rate in the collision window λ_col_ less than –3.5 and *p* < 0.005) with low response jitter (< 20 ms). Jitter threshold was applied here to ensure accuracy of τ_anti_ estimation.

#### Histology

After experiments were completed, mice were euthanized with an overdose of isoflurane, exsanguinated with saline, and perfused with 4% paraformaldehyde. For mice with tetrodes, the brains were cut in 100-µm-thick coronal sections and mounted on glass slides. We validated expression of rAAV5-EF1a-DIO-hChR2(H134R)-EYFP with epifluorescence images of LC (Zeiss Axio Zoom.V16) and confirmed targeting of the optic-fiber-tetrode bundle to LC by colocalization of the electrolytic lesion with immunostaining against TH (rabbit anti-TH, ab152, Millipore, followed by donkey anti-rabbit Alexa 488, Life Technologies, or goat anti-rabbit Alexa 594, Life Technologies). In electrophysiological experiments with Neuropixels probes, probe tracks were reconstructed as described below.

#### Probe reconstruction

In Neuropixels recordings, probes were dipped into florescence dye before penetrating the brain. After 1 week of recording (3–5 recording sessions), mice were perfused with PFA, brains were extracted and cleared with a LifeCanvas protocol, and imaged using SPIM. Images were processed through a SmartSPIM pipeline (github.com/AllenNeuralDynamics/aind-smartspim-pipeline), where images were stitched and registered to CCF.

In stitched images, probe tracks were manually labeled and transformed into CCF coordinates. To further refine the location of the probe (Liu et al. 2021), electrophysiological features along the probe were aligned to brain regions along the probe track using a modified version of International Brain Laboratory software (github.com/AllenNeuralDynamics/ibl-ephys-alignment-gui). Key features for identifying the locations of recorded cells dorsal to LC include the discontinuity of spatial cross correlation of local field potentials at the dorsal edge of the pons, change of signal amplitude among cerebellum layers, and transient signal change at the brain surface.

#### Drift control

In recordings with Neuropixels probes, spike detection and clustering are subject to movement artifacts when the brain moves relative to the probe. We controlled for both abrupt movement and slow drift of the probe.

To remove the effects of abrupt displacement, we restricted our analyses to time windows without such events. To identify when abrupt changes occurred, we tested whether sudden changes in spike rate could be predicted by concurrent changes in spike waveform amplitude and probe position. Specifically, we estimated the first derivative of spike rate λ′(*t*), probe displacement *d*′(*t*) and spike waveform peak amplitude *p*′(*t*) from one channel for each neuron, at each time point *t*. For a generic signal *v*(*t*) at time *t*_0_, the first derivative was estimated using a windowed contrast measure defined as follows:

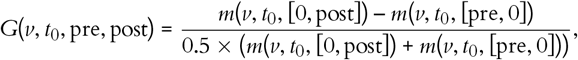

where *m*(*v, t*_0_, [*a, b*]) denoted the windowed mean of signal *v* in the interval [*t*_0_ + *a, t*_0_ + *b*), defined as

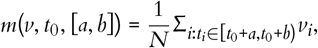

where *N* = |{*i* : *t*_*i*_ ∈ [*t*_0_ + *a, t*_0_ + *b*)}|. The first derivatives were estimated every 100 s. A regression model was fitted to predict |λ′(*t*)| using |*p*′(*t*)| and |*d*′(*t*)| as, |λ′| ∼ 1 + |*d*′| + |*p*′|. The estimated absolute value of the first derivative of firing rate is 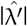. This analysis was performed at two different time scales, short and long, with pre = –300 s for both time scales, post = 100 s for short time scales, and post = 300 s for long time scale calculations. Abrupt drift events were detected at time *t* when both λ′(*t*) and 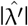 exceeded 0.5, with either short or long time-scale calculations.

We also estimated slow drifts of the probe over long timescales. A neuron’s overall relative spike rate change over a whole session was calculated as the ratio of standard deviation of spike rate divided by mean spike rate, calculated by binning the session into non-overlapping 300-s bins. NE neurons with this ratio higher than 0.3 (98th percentile) after being restricted to periods without abrupt drift events were excluded from analysis.

To avoid potential effects of slow electrode motion on spike sorting, one drift regressor was added to all single neuron analyses with linear regression. The drift regressor was the deviation of mean waveform amplitude on its peak channel for each trial from the mode of waveform amplitudes.

#### Axonal calcium indicator measurements

To express GCaMP6s, or jGCaMP8m (Zhang et al. 2023) in LC-NE neurons, we pressure-injected within LC AAV-hSyn-FLEX-axon-GCaMP6s (1.0 × 10^13^ GC mL^−1^) or AAV-hSyn-FLEX-Axon-jGCaMP8s (1.0 × 10^13^ GC mL^−1^) into the LC of *Dbh*-Cre mice at a rate of 1 nl s^−1^. pAAV-hSynapsin1-FLEx-axon-GCaMP6s was a gift from Lin Tian (Addgene viral prep 112010-AAV5; RRID:Addgene_112010) (Broussard et al. 2018) For Axon-jGCaMP8s, based on the Axon-GCaMP6s sequence (Addgene: 112010), we PCR-amplified AvrII-Axon-jGCAMP8s-NheI. Then, the construct was ligated into the hSynapsin1-FLEX pAAV backbone cut from pAAV-Syn-FLEX-rc[ChrimsonR-tdTomato] (Addgene: 62723) with AvrII/NheI.

Viruses were injected at following locations: from bregma, 5.4 mm posterior for male mice (400 nL), 5.2 mm posterior for female mice (300 nL); 0.85 mm bilateral; and (2.9, 3.1) mm from surface.

#### Fiber implantation

We implanted 200-µm-diameter optic fibers (0.39 NA) above prelimbic cortex (2.0 mm anterior to bregma, 0.5 mm lateral, 1.5 mm ventral from the pial surface). Detail can be found here: https://dx.doi.org/10.17504/protocols.io.6qpvr3dqovmk/v1. We measured GCaMP6s and GCaMP8s fluorescence using 60 µW irradiance of 488-nm (to excite GCaMP6s and GCaMP8s) and 40 µW 405-nm (as an isosbestic control) light using a custom photometry system (dx.doi.org/10.17504/protocols.io.261ge39edl47/v2). Signals were sampled at 60 Hz, alternative sampling from 405 nm, 488 nm and 560 nm (0 irradiance from 560 nm in these experiments) channels, resulting in 20 Hz for each channel. Example raw traces are in Figure S17c.

#### Data preprocessing

Raw photometry traces from both channels (488 nm and 405 nm) were first de-trended by fitting and removing a tri-exponential curve, each modeling signal decay with different timescales. ΔF/F was computed as the ratio of the detrended signal and the trend. After this, we generated a low pass filtered baseline at 0.05 Hz for both channels’ ΔF/F and fitted the isosbestic channel’s (405 nm) baseline to the 488 nm channel’s baseline. The fitted baseline was treated as a movement artifact estimated by the isosbestic channel. This baseline was removed from ΔF/F from the 488 nm channel, generating a motion corrected detrended signal. This signal was then low pass filtered at 3 Hz before further analysis. Code can be found here: https://github.com/AllenNeuralDynamics/aind-fip-dff.

#### Comparison between single neuron activity and axonal GCaMP

To test whether LC-NE neurons projecting to cortex differed from the whole distribution of neurons, we compared task responses of single LC-NE neurons identified as projecting to cortex versus those that were not. To compare *t*-statistics from regression models between these two groups of neurons, we used Welch’s two-sample *t*-test (two-sided), which allows unequal variances. In addition to the asymptotic *p*-value, significance was also assessed using a label-permutation test in which pooled observations were randomly reassigned to groups while keeping group sizes. Effect size was quantified using Cohen’s *d*, and 95% confidence interval was estimated from bootstrapping by resampling each group with replacement across bootstrap iterations.

Comparisons between single-neuron spiking and photometry-derived measures, are sensitive to modality-dependent differences in effect magnitudes or signal-to-noise ratios. Thus, we used a sign-based comparison to compare single-neuron spiking to photometry. To compare the sign of effects (*t*-statistics from regression models), we computed the proportion of positive values in each group and tested for differences between groups. Group differences were quantified using the risk difference, risk ratio, and odds ratio. Statistical significance was evaluated using a two-proportion *z*-test, and a label-permutation test on the difference in positive rates obtained by shuffling group labels while keeping group sizes.

We tested whether the sample of LC-NE neurons projecting to cortex were biased relative to the distribution across all neurons. We compared *t*-statistics values from regression models between these two groups of neurons, we used Welch’s two-sample *t*-test (two-sided), which allows unequal variances. In addition to the asymptotic p-value, significance was also assessed using a label-permutation test in which pooled observations were randomly reassigned to groups (single neuron activity or calcium indicator dynamics) while keeping group sizes. Effect size was quantified using Cohen’s *d*, and 95% confidence intervals were estimated from bootstrapping by resampling each group with replacement across bootstrap iterations. The permutation test served as the primary significance test because sample size of neurons identified to project cortex is small.

#### Data preprocessing

Raw photometry traces from both channels (488 nm and 405 nm) were first de-trended by fitting and removing a tri-exponential curve, each modeling signal decay with different timescales. ΔF/F was computed as ratio of detrended signal over the trend. After this, we generated a low pass filtered baseline at 0.05 Hz for both channels’ ΔF/F and fitted isosbestic channel’s (405 nm) baseline to 488 nm channel’s baseline. The fitted baseline was treated as a movement artifact estimated by the isosbestic channel. This baseline was removed from ΔF/F from the 488 nm channel, generating a motion corrected detrended signal. This signal was then low pass filtered at 3 Hz before further analysis. Code can be found here: github.com/AllenNeuralDynamics/aind-fip-dff.

#### Circular distribution comparison

When comparing differences between outcome-related activity of neurons projecting to cortex vs. not and neuronal spiking vs. calcium indicator dynamics in PL, differences between circular distributions were assessed using a permutation-based implementation of the Mardia–Watson–Wheeler test. Angular observations were first converted to floating-point values. All angles were then wrapped to the interval [0, 2π) using a modulo 2π transformation to ensure a common circular reference frame.

For two samples containing *n*_1_ and *n*_2_ observations, respectively, angles from both groups were pooled and assigned ranks across the combined dataset using average ranks for tied values. The pooled ranks *r*_*i*_ were then mapped onto the circle as ϕ_*i*_ = 2π*r*_*i*_/*n*, where *n* = *n*_1_ + *n*_2_. For each group *g*, the summed cosine and sine components of the transformed angles were computed as *C*_*g*_ = Σ_*i*_∈_*g*_cos(ϕ_*i*_), *S*_*g*_ = Σ_*i*_∈_*g*_sin(ϕ_*i*_).

The Mardia–Watson–Wheeler statistic was calculated as

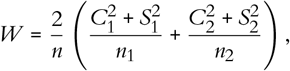

which tests the null hypothesis that the two samples are drawn from the same circular distribution.

Statistical significance was evaluated using a permutation procedure. Group labels were randomly permuted across the pooled angles while preserving the original group sizes, and the test statistic was recomputed for each permutation. This procedure was repeated 5,000 times using a pseudo-random number generator. The *p*-value was estimated as *p* = (*k* + 1)/(*N*_perm_ + 1), where *k* denotes the number of permutations yielding a statistic greater than or equal to the observed value and *N*_perm_ is the total number of permutations. Analyses were performed in Python using custom code based on the rank-based formulation of the Mardia–Watson–Wheeler statistic. Note that because angles compared here are independent of the magnitude of *t*-statistics, they are not subject to differences of signal-to-noise ratios from different data modalities.

#### Pupil tracking

Pupil video was acquired at 22–24 frames per s with a telecentric lens (Edmund Optics 58-430). Time synchronization was performed by adding an LED reflection into mice’s eye, next to pupil, which turns on at the start of each trial. Synchronization signal and pupil edges were detected using a DeepLabCut model (version 2.1.6.2). Two models were trained using 314 and 208 frames from 9 and 13 mice, respectively, and were further refined based on tracking performance. Pupil boundaries were labeled on the left-most and right-most edges. Edge detection confidence lower than 0.9 was excluded from analysis. Signals were low-pass filtered with a 2nd-order Butterworth filter (cutoff, 5 Hz) using zero-phase forward–backward filtering.

#### Pose estimation

We collected video during task performance at a framerate of 500 Hz and resolution of 720 × 540 pixels using FLIR cameras (Blackfly S BFS-U3-04S2M). We trained a Lightning Pose model (Biderman et al. 2024) to detect the distal tip of the tongue at the midline (mean error of 4.47 pixels). The model was trained on 1200 labeled frames across 12 behavioral sessions and 8 mice. Tongue tip location was predicted across all analyzed sessions, and subsequent analysis was limited to sessions with high model performance as measured by >90% mutual agreement with the capacitive lick sensor and > 60 ms median tongue movement duration, which excluded sessions with poor model generalization.

For signal processing, kinematic data (tongue tip position over time) was masked for model confidence at 0.90. Data were then filtered using a 50 Hz, low-pass, symmetric, fourth-order Butterworth filter. Reaction time was defined per trial as the time elapsed from the go cue until the first timepoint in which the tongue was detected, with maximum cutoff set at 1.0 s.

Correlations between spike counts and reaction time were nonparametric. Baseline window was defined as 1.0 s prior to the go cue. Response window was defined as 0.2 s following the go cue, prior to the median reaction time across sessions. *t*-statistics were calculated from linear regression of reaction time to spike counts in specified windows on each trial.

#### Lick bout detection

Licks recorded from lick sensor contact were first processed to remove lick detection noise either from signal rebound or crosstalk between two lick sensors. Licks detected from post tracking in videos were constrained to licks with a) existing time longer than 0.02 s and shorter than 95th percentiles; b) end of lick movement location, peak velocity, and total distance all smaller than 95th percentiles. Curated licks detected by lick sensors or by video were segmented into lick bouts where inter-lick-interval was longer than 0.5 s.

#### Data analysis

All data are presented as mean ± SEM unless reported otherwise. All statistical tests were two-sided. For all analyses, no-go cues (when presented) were treated as part of the inter-trial interval. Code is available at github.com/AllenNeuralDynamics/aind_stan_fit_sim, github.com/AllenNeuralDynamics/aind-beh-ephys-analysis, github.com/JeremiahYCohenLab/sueAnalysis/tree/master/python/pupillometry.

#### Analyses and models of behavior

We fit logistic regression models to predict choices and engagement as a function of outcome history for each mouse. To predict choices, we used this model:

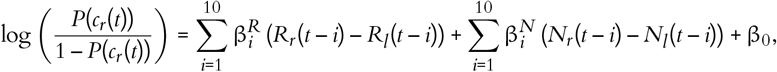

where *c*_*r*_ (*t*) = 1 was a rightward choice and 0 for a leftward choice. *R* = 1 was a rewarded choice and 0 for an unrewarded choice, and *N* = 1 was an unrewarded choice and 0 for a rewarded choice.

To predict engagement, we used this model:

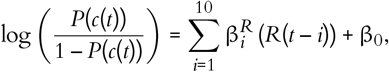

where *c*(*t*) = 1 was a response to the go cue and 0 for ignoring a go cue, and *R* = 1 was a rewarded choice regardless of direction.

To predict switch versus stay choices, we used this model:

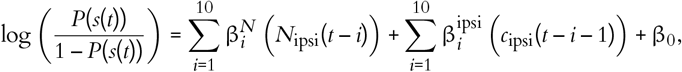

where *s*(*t*) is a switch choice, *N*_ipsi_ = 1 was an unrewarded choice ipsilateral to the choice at *t* – 1, –1 for an unrewarded choice contralateral to the choice at *t* – 1, and 0 for a rewarded choice.

We applied a family of reinforcement-learning models of behavior used previously (Grossman et al. 2022; Bari et al. 2019). This model estimates action values (*Q*_*l*_ (*t*) and *Q*_*r*_ (*t*)) on each trial to generate choices. Choices are described by a random variable, *c*(*t*), corresponding to left or right choice, *c*(*t*) ∈ *l, r*. The value of a choice is updated as a function of the RPE, and the rate at which this learning occurs is controlled by the learning rate parameter α. To account for asymmetric learning from rewards and no rewards, we used separate learning rates for each outcome. For example, if the left spout was chosen, then

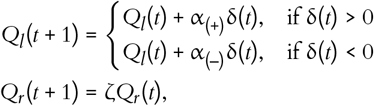

where δ(*t*) = *R*(*t*) – *Q*_*l*_ (*t*) and ζ represents the forgetting rate parameter. The forgetting rate captures the increasing uncertainty about the value of the unchosen spout.

The *Q*-values were used to generate choice probabilities through a softmax decision function:

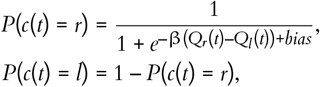

where β, the “inverse temperature” parameter, controls the steepness of the sigmoidal choice function. In other words, β controls the stochasticity of choice. This model was used for experiments with electrophysiology and GCaMP measurements.

To compare change in choice dynamics against RPE, we estimated the change of choice likelihood in consecutive trials. This quantity, defined as

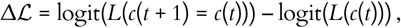

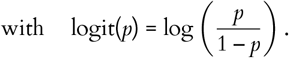

determines how future behavior is altered following RPE. ℒ (*c*(*t* + 1) = *c*(*t*)) was the likelihood of a mouse choosing current choice in the future, estimated from the sequence of mouse choices using logistic regression model. ℒ (*c*(*t*)) represents animals’ choice policy before outcome, it was the likelihood estimated from the *Q*-learning model. Change of likelihood was centered in each session to account for choice autocorrelations.

#### Behavioral model fitting

We fit and assessed models using python and the probabilistic programming language, Stan (https://mc-stan.org/) with the PyStan interface. Stan was used to construct hierarchical models with mouse-level hyperparameters to govern session-level parameters. This hierarchical construction uses partial pooling to mitigate overfitting to noise in individual sessions (often seen in the point estimates for session-level parameters that result from other methods of estimation) without ignoring meaningful session-to-session variability. For each session, each parameter in the model (for example, α) was modeled as a draw from a mouse-level distribution with mean µ and variance σ. Models were fit using noninformative (uniform distribution) priors for mouse-level hyperparameters (Table 3).

**Table 3:** Model parameters.

| Parameter | Prior distribution |
| --- | --- |
| $\alpha_{(-)}$ | $[0, 1]$ |
| $\alpha_{(+)}$ | $[0, 1]$ |
| $\alpha$ | $[0, 1]$ |
| $\zeta$ | $[0, 1]$ |
| $\beta$ | $[0, 10]$ |
| bias | $\mathcal{N}(0, 1)$ |

Weakly informative priors were used for session-level parameters. Mouse-level hyperparameters were chosen to achieve model convergence under the assumption that individual mice behave similarly across days. The parameters were sampled in an unconstrained space and transformed into bounded values (for those parameters that were bounded) by a standard normal inverse cumulative density function. Stan uses full Bayesian statistical inference to generate posterior distributions of parameter estimates using Hamiltonian Markov chain Monte Carlo sampling (Carpenter et al. 2017). The default no-U-turn sampler was used. The Metropolis acceptance rate was set to 0.85–0.9 to force smaller step sizes and improve sampler efficiency. The models were fit with 5,000 iterations and 2,500 warmup draws run on each of 16 chains in parallel. Default configuration settings were used otherwise (https://github.com/AllenNeuralDynamics/aind_stan_fit_sim).

#### Extracting model parameters and variables, behavior simulation

For extracting model variables (like RPE), we took at least 1,000 draws from the Hamiltonian Markov chain Monte Carlo samples of session-level parameters, ran the model agent through the task with the actual choices and outcomes, and averaged each model variable across runs. For comparisons of individual parameters across models, we estimated *maximum a posteriori* parameter values by approximating the mode of the distribution: binning the values in 50 bins and taking the mean value of the most populated bin.

#### Linear regression models of spike rates

To characterize neurons’ correlation with task components, we performed linear regression in different time windows. To account for the potential contribution of slow drift of the probe, mean spike amplitude around each trial was included as a regressor and contributed to prediction of spike counts in on average 25.4% of all neurons across different regression analysis.

To determine how neurons responded to different outcomes (presence or absence of reward) from different choices (ipsilateral or contralateral to the recording site), we regressed firing rates on outcome (*R*), choice side (*c*(*t*), 1 for the direction ipsilateral to the recording site, –1 for contralateral to the recording site), reward prediction (*Q*_*c*_), and their interaction, using the Python package scikit-learn. Because there was no external sensory information to indicate that the mouse successfully made a choice and that water would or would not arrive, we defined separate windows for analysis of reward and no reward. Regressions were performed for each neuron in both a reward response window and a no-reward response window and the one with larger absolute *t*-statistics were used for each neuron.

To quantify how each neuron responds to outcome, the area under the receiver operating characteristic curve (auROC) was computed in a sliding window of 1 s, from response time to 2 s after the response time (Figure S15). Each neuron’s peak “outcome response time” was when the auROC values were furthest from 0.5. A neuron’s outcome response sign was defined as positive if this auROC value was greater than 0.5 and negative if it was less than 0.5. Across the population, we estimated the distribution of “outcome response time” in neurons with positive and negative response signs. The modes of the distributions were taken as the positive and negative outcome response windows.

We quantified regressions using both *t*-statistics of regressors and the angle in Cartesian coordinates of the estimates of coefficients for *R* and *Q*_*c*_ for each neuron. The latter was plotted as a polar histogram (Figure 6).

To quantify each neuron’s relationship to task engagement, we performed linear regression on the baseline firing rate 2 s before each go cue with presence or absence of a lick response to the go cue as a regressor.

To quantify each neuron’s relationship to choice (switch versus stay), we performed linear regression on the go cue period’s spike rate (0.5 s after go cue) with choice as a regressor.

#### Linear regression models of axonal GCaMP

To compare axonal activity correlation with behavior, we performed linear regression in different time windows. To address movement artifacts, the isosbestic channel’s signal from same time windows were included as additional regressor.

We applied a quality control threshold on the LC-NE axon dynamics, including data with signal increases (z-scored across the whole recording) higher than 0.5 z-score ΔF/F.

To determine how LC-NE axons responded to different outcomes, we calculated regressions as for the single neuron analysis (presence or absence of reward, ipsilateral or contralateral choice relative to the recording site, their interactions, and chosen value). Because axonal ΔF/F has lower temporal resolution than single neuron spiking, a single larger time window (2 s) after outcome was used to compute the regression model for outcome encoding.

To quantify how LC-NE axons correlated with task engagement, we calculated linear regression on the baseline firing rate 2 s before each go cue with presence or absence of a lick response to the go cue as a regressor.

To quantify how LC-NE axons correlated with choice (switch versus stay), we performed linear regression on the go cue period’s firing rate (0.75 s after go cue) with choice as regressor.

We quantified regressions using both *t*-statistics of regressors and the angle in Cartesian coordinates of the estimates of coefficients for *R* and *Q*_*c*_ for each neuron. The latter was plotted as a polar histogram (Figure 6).

#### Pupil diameter analysis with behavior

To test how pupil diameter changes with behavior events. A linear regression model was fitted to a sliding window of 1 s across the trial time, aligned to the start of go cues. Regressors included choice outcome, choice side (ipsilateral or contralateral to pupil recording side), and whether mice changed choice or repeated the previous choice. Distribution of *t*-statistics are reported in Figure S16f.

#### Pupil auto-correlogram

To capture temporal statistics of pupil diameter, auto-correlogram were computed across the whole session of pupil diameter recording with a sliding window of 0.5 s, with period of poor time alignment or poor pupil edge detection removed. An exponential curve was fitted to the auto-correlation curve: pupil = *Ae*^−*t*/τ^ + *C*. τ had a mean of 3.37 and SD of 1.03.

#### Pupil correlation with neuron activity

To analyze how LC-NE activity correlated with pupil diameter on different timescales, we performed three types of correlation analysis focusing on different time windows. First, to have a general estimate of task-independent correlation between pupil diameter and neuron activity on long timescales, we calculated cross-correlations between the two. Mean spike rate and mean pupil diameter were computed using a sliding window of 5 s. Cross-correlations were estimated with a maximum lag of 20 s, advancing in steps of 0.1 seconds. Pupil diameter was first de-trended using a linear fit of time to remove the effects of slow pupil diameter changes on the timescale of the whole session. This correlation was computed both including and excluding the trial period (0-3 s from go cues).

Second, to quantify spike-pupil relationships during iter-trial-intervals before go cues, we computed the trial-wise correlation between baseline spike rate and baseline pupil diameter 1 s before the onset of the go cue. Third, to compare baseline neuronal activity to pupil dilation, we computed trial-wise correlations of spike rates 1 s before go cues with pupil dilation (maximum pupil increase from baseline pupil diameter, 1 s before go cue).

#### Comparison of phasic and tonic responses

To quantify how tonic activity (pre-trial spike rate) correlated with phasic responses to the go cue, we calculated correlation coefficients 0.5 s before the go cue 0–0.3 s after the go cue. We compared against a null model where we computed this correlation with “sham” windows, randomly sampled from a uniform distribution between the first and last go cue.

#### Analyzing relationships with space

We analyzed the relationship between space and both one-dimensional and high-dimensional data. To test whether a particular feature (or group of features) was randomly distributed in space or instead shows spatial structure, we quantified two complementary forms of spatial dependence using the corresponding CCF coordinates *X* = (*x, y, z*). The analysis assessed (i) the presence of a global linear trend across space and (ii) the extent to which values were more similar to their neighbors than randomly assigned neurons. All significance testing was performed using permutation tests that preserve the spatial sampling geometry by keeping coordinates fixed while randomly reassigning values across locations. When assessing a significant linear trend, we also estimated the primary spatial vector and its confidence interval for each feature (or group of features). Any neuron containing non-finite entries in either its location or feature values were removed. Analyses were only performed when at least 100 valid spatial samples remained after filtering. Medial-lateral (ML) coordinates were mirrored to the left hemisphere by taking the absolute value of ML, so that left and right locations contributed to the same axis estimate.

#### Global linear trend test for single features using linear regression

We first asked whether *y* exhibits a global linear gradient across space by fitting an ordinary least squares (OLS) regression model, *y* = β_0_ + β*X* + ϵ, where β_0_ is an intercept and β ∈ *R*^*D*^ are slope coefficients for each spatial dimension. The model fit was summarized using the coefficient of determination 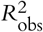, which quantifies the fraction of variance in *y* explained by a linear function of space. β was normalized and used as the primary axis of the spatial trend.

To determine whether the observed linear trend exceeded what would be expected by chance given the same spatial sampling, we constructed a null distribution by repeatedly permuting *y* while leaving *X* fixed. For each permutation *b* = 1, …, *B* (*B* = 5000), we computed 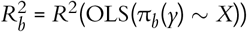, where π_*b*_(·) denotes a random permutation. A one-sided permutation p-value was then computed as 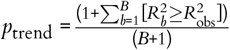, testing whether the observed explained variance is larger than expected under spatial randomness. Reported outputs include unit primary spatial axis, 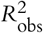, F-statistic, permutation *p*-value, and the mean and SD of the permuted *R*^2^ distribution.

#### Local spatial predictability test for single features using k-nearest neighbors (kNN) test

Global linear trends can miss nonlinear or locally structured spatial patterns (e.g., patches or clusters). Therefore, we additionally quantified local spatial dependence by testing whether values at a location were predictable from nearby locations using distance-weighted *k*-nearest neighbors’ regression. We performed *K*-fold cross-validation (default *K* = 5), shuffling samples with a fixed random seed for reproducibility. For each fold, a kNN regressor was trained on the training set using the spatial coordinates *X*, and predictions were generated for held-out points. The number of neighbors was set to *k* = min(*k*_neighbors_, *n*_train_), (default *k*_neighbors_ = 20) unless noted otherwise) with inverse-distance weighting so that closer neighbors contribute more strongly.

Predictive performance was summarized as the cross-validated coefficient of determination, 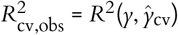, where ŷ_cv_ denotes concatenated out-of-fold predictions across all folds (default, 5). To evaluate whether local predictability exceeded chance, we generated a permutation null distribution by permuting *y* across locations and recomputing the full cross-validated kNN *R*^2^ for each permuted dataset: 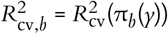.

A one-sided *p*-value was computed analogously: 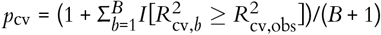.

Reported outputs include 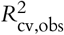, permutation *p*-value, and the mean and SD of the null distribution.

#### Global linear trend test for single features using linear discriminant analysis (LDA)

To identify whether categorical labels exhibited a structured spatial organization, we estimated a primary spatial axis that best separated labels using LDA. Specifically, we modeled the relationship between class labels *y* and spatial coordinates *X* ∈ ℝ^*n*^×^3^ by fitting an LDA model, which finds a linear projection of the coordinates that maximizes between-class variance relative to within-class variance.

Given coordinates *X* and categorical labels *y*, LDA estimates a set of discriminant directions β ∈ ℝ^3^ such that the projected values *X*β optimally separate the label classes. For multiclass settings, the number of discriminant components is bounded by min(*D, K* – 1), where *D* = 3 is the number of spatial dimensions and *K* is the number of unique classes. In all analyses, we retained only the first discriminant component, corresponding to the direction of maximal class separability. The resulting discriminant vector β was extracted from the fitted model (using the feature-space projection matrix when available), and normalized to unit length to define the primary spatial axis as 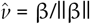.

The unit vector 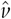 represents the dominant spatial direction along which class labels were most strongly differentiated. For consistency with linear regression-based analyses, we also report the raw coefficients β, while the intercept term was set to zero as it is not meaningful in this geometric interpretation. Reported outputs include the unit primary spatial axis 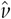, the unnormalized discriminant vector β, and model validity checks based on sample counts and class balance.

#### Global linear trend test for groups of features

To test whether a group of related variables (e.g., different waveform features or expression of multiple genes) had a linear trend in space, we applied CCA. Values were z-scored prior to analysis. CCA was performed using the scikit-learn implementation (sklearn.cross_decomposition.CCA), extracting canonical variates that maximize correlation between linear combinations of functional and anatomical variables. The strength of association was quantified as the linear correlation between paired canonical projections. The weight from the spatial location was then normalized and used as the primary axis of spatial variation.

To assess statistical significance, we performed a permutation test in which anatomical labels were randomly shuffled across units (10,000 permutations). For each shuffle, CCA was recomputed, and the maximum canonical correlation was recorded to generate a null distribution. Empirical p-values were computed as the proportion of shuffled correlations exceeding the observed correlation.

#### Bootstrap estimation of spatial vector confidence intervals

To estimate confidence intervals for primary spatial vectors inferred from the methods mentioned above, we performed bootstrap resampling with replacement across units (5,000 iterations). In each iteration, rows of the original dataset (one row for each unit) were resampled with replacement, and spatial vectors were re-estimated using corresponding methods. Because the sign of axis direction was arbitrary, each bootstrapped axis was aligned to the observed axis by flipping its sign whenever its inner product with the observed axis was negative. This ensured that all bootstrapped axes were in the same hemisphere and prevented artificial bimodality in the bootstrap distribution.

To visualize uncertainty, bootstrapped axis distributions were plotted in azimuth-elevation space and summarized as a 95% confidence cone around the observed axis (Figure S18). The cone half-angle was defined as the 95th percentile of the angular deviation between the bootstrapped axes and the observed axis. For plotting in 2D anatomical planes, the boundary of this 3D cone on the unit sphere was projected into each plane and displayed as a shaded confidence region around the projected axis arrow (Figure S18).

#### Comparison of two primary spatial axes

To compare whether two inferred primary spatial axes differed in direction, each pair of unit vectors from bootstrapping was represented in a local tangent-plane coordinate system. Given a reference axis *b*_*x*_ (one of the primary spatial axes to compare), we constructed an orthonormal basis (*e*_1_, *e*_2_, *u*_0_), where *u*_0_ = *b*_*x*_/||*b*_*x*_|| is the normalized reference axis. The first tangent direction *e*_1_ was defined as the normalized projection of the second axis onto the plane perpendicular to *u*_0_,

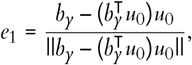

and the second tangent direction was defined as *e*_2_ = *u*_0_ × *e*_1_. This basis defines a two-dimensional plane orthogonal to the reference axis. The observed directional deviation of *b*_*y*_ from *b*_*x*_ was represented by projecting the second axis into this plane, *d*_obs_ = *Ab*_*y*_, where *A* is a 2 × 3 projection matrix,

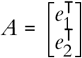

Bootstrap uncertainty for the difference between axes was estimated from the corresponding bootstrap axis estimates 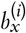 and 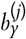. Each bootstrap vector was first aligned to its observed axis by flipping its sign when the inner product with the observed axis was negative. Bootstrap vectors were then projected into the same tangent plane, 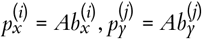.

Since bootstrap samples were not paired across datasets, the sampling distribution of directional differences was approximated by randomly sampling combinations of bootstrap projections from the two datasets and computing 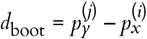. This procedure generates a bootstrap cloud that approximates the sampling distribution of the two-dimensional directional difference. In the case where there are 2,000 bootstraps, 10,000 pairs of bootstrapped vectors were randomly sampled to approximate the distribution.

The magnitude of the observed directional difference was quantified using the Mahalanobis distance of the observed deviation from zero, 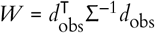, where Σ = *Cov*(*d*_boot)_ is the covariance matrix of the bootstrap difference cloud.

To compute a bootstrap-based *p*-value, the bootstrap difference distribution was first recentered to estimate the null distribution, 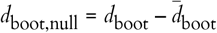. For each recentered bootstrap sample, a quadratic form was computed 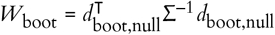. The bootstrap p-value was then defined as

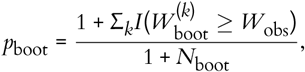

where *N*_boot_ is the number of bootstrap difference samples and *I*(·) is the indicator function.

The bootstrap difference cloud was also used directly to visualize uncertainty and to assess whether the 95% confidence ellipse included the origin, corresponding to no directional difference. Finally, the angular separation between axes was reported as 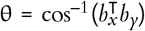, which provides an intuitive summary of directional difference between the two spatial axes.

#### Relating behavioral features to spatial axes

To assess how behavioral encoding varied along each primary spatial axis inferred from other data modalities, each neuron’s anatomical coordinate was projected onto each axis by taking the dot product between its 3D coordinate and the unit spatial vector: 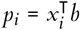. This yielded a one-dimensional coordinate along the waveform axis, the MERFISH axis, and the retrograde-label axis.

For each behavioral feature, the projected coordinate was related to the feature value using Pearson correlation. Correlation coefficients and corresponding *p*-values were calculated using only samples with non-missing values. For visualization, a simple linear regression line was fit to the relationship between projected coordinate and behavioral feature, and a 95% confidence band for the fitted line was computed from the standard linear regression uncertainty estimate. Each behavioral feature was therefore compared across multiple spatial embeddings by repeating the same projection-and-correlation analysis for each inferred axis (Figure S18).

#### Spike count cross-correlations

To estimate spike rate correlations among simultaneously recorded LC-NE neurons, we counted spikes in non-overlapping windows of 50 ms and calculated cross-correlations between them. To extract the main features of cross-correlograms, PCA was applied.

## Data and code availability

Code is available at https://github.com/AllenNeuralDynamics/locus_coeruleus_norepinephrine_neurons and code and data are available at https://codeocean.allenneuraldynamics.org/collections/9cf044ce-93c7-4c7e-bfa1-5d8c37aa42ec.

## Acknowledgments

We thank A. Amaya, J. Baka, B. Bari, A. Bawany, D. Bertagnolli, Y. Browning, A. Cahoon, J. Friedrich, J. Goldy, C. Grasso, A. Grim, C. Grossman, J. Guzman, W. Han, R. Howard, K. James, T. Johnson, J. Jung, M. Kannan, J. Kenney, A. Leon, H. Loeffler, F. Lucantonio, Y. Marghi, N. Mars, R. Naidoo, K. Nguyen, B. Ouellette, C. Rimorin, D. Rocha, H. Rodrigues, A. Sridhar, A. Stearns, L. Suarez, U. Sümbül, J. Swapp, M. Tom, A. Torkelson, J. Wang, J. Weber, J. Wilkes, A. Williford, J. Wong, and B. Wynalda. This publication was supported by and coordinated through the Brain Initiative Cell Atlas Network (BICAN). Research reported in this publication was supported by the National Institutes of Health BRAIN Initiative under award numbers R01MH134833, R01NS104834, and RF1NS131984. This work was supported by the Allen Institute.

## Author contribution matrix

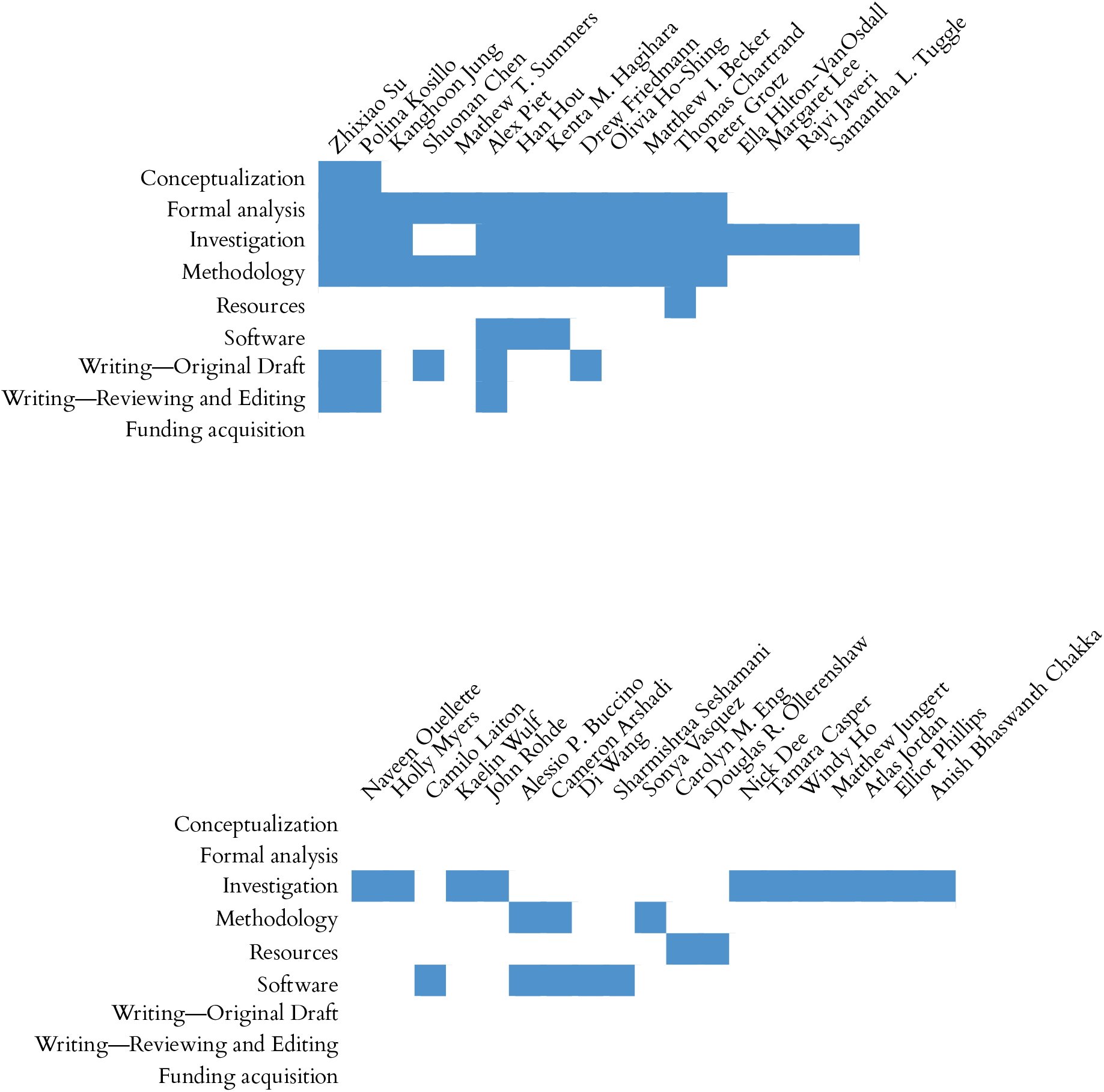
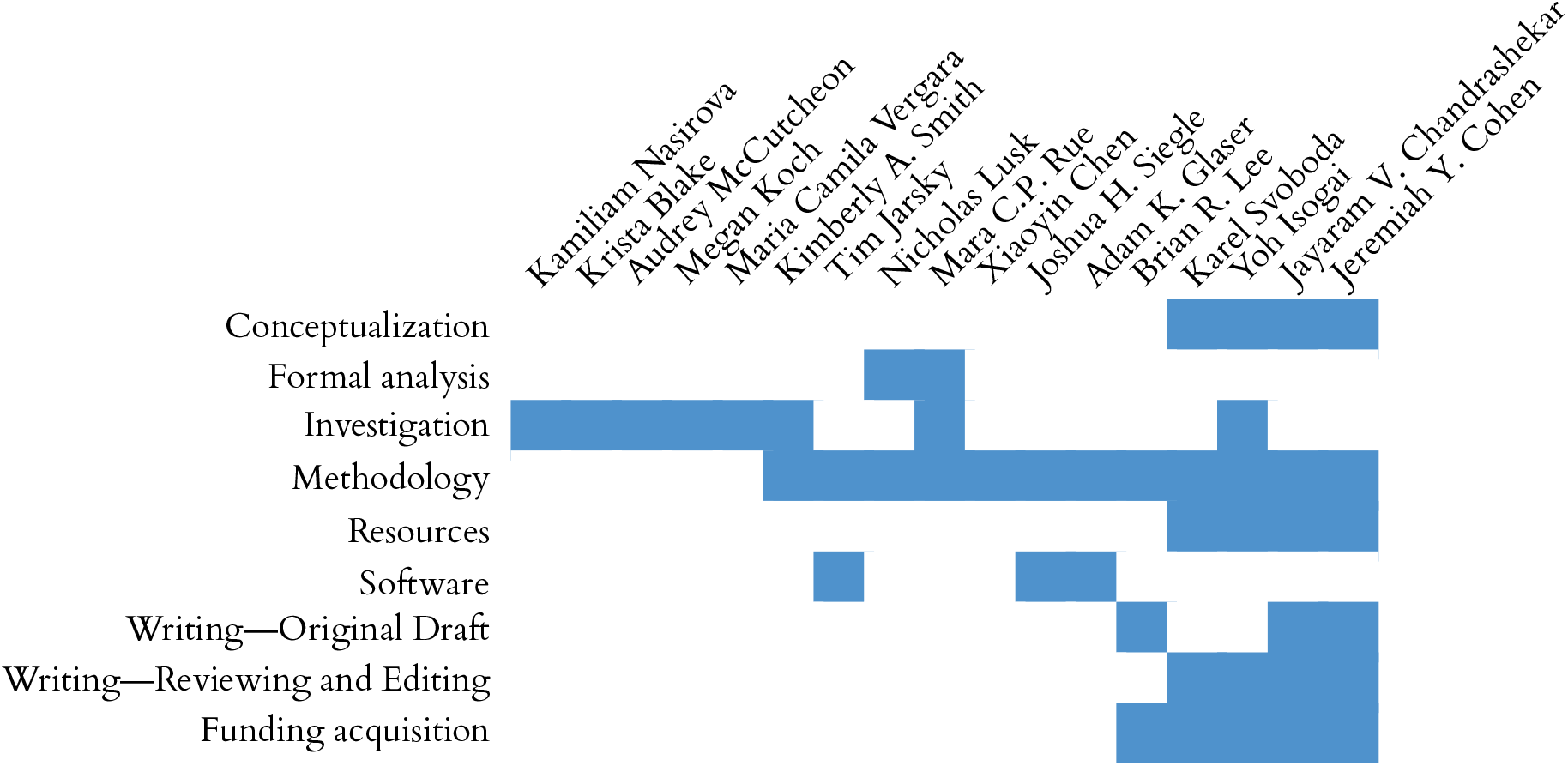

---

**Figure S1:**
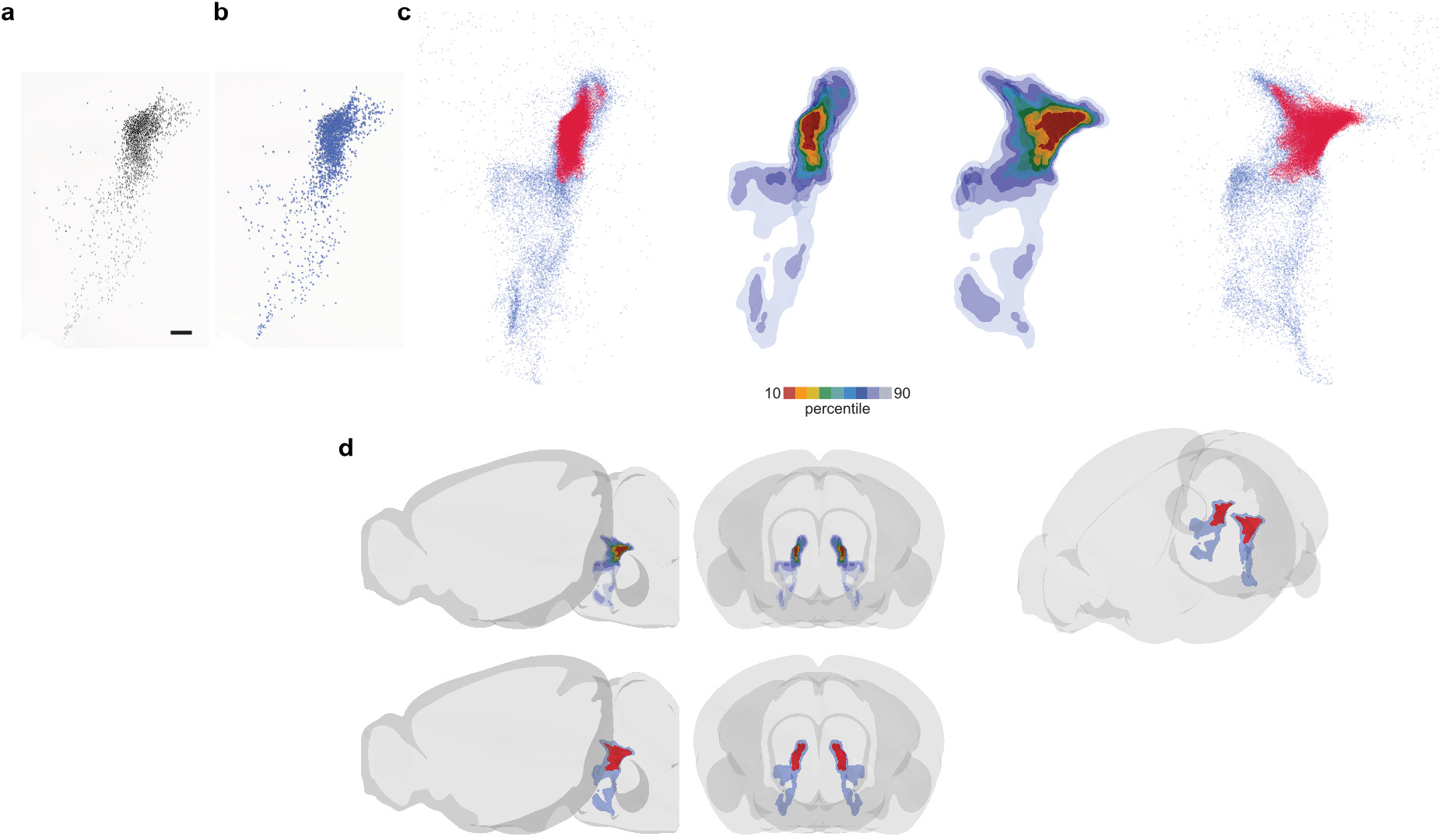
Spatial distribution of LC-NE neurons. **a**, Projection of H2B+ neurons from a 500-µm volume in the coronal plane (reproduced from Figure 1). Scale bar, 100 µm. **b**, Localization of H2B+ neurons from (a). **c**, CCF-localized *Dbh*+ cell bodies projected in the coronal (left) or sagittal (right) planes across mice. Color scale indicates percentage of cells that lie inside the labeled boundaries. **d**, LC atlas in the pons depicted in three planes.

**Figure S2:**
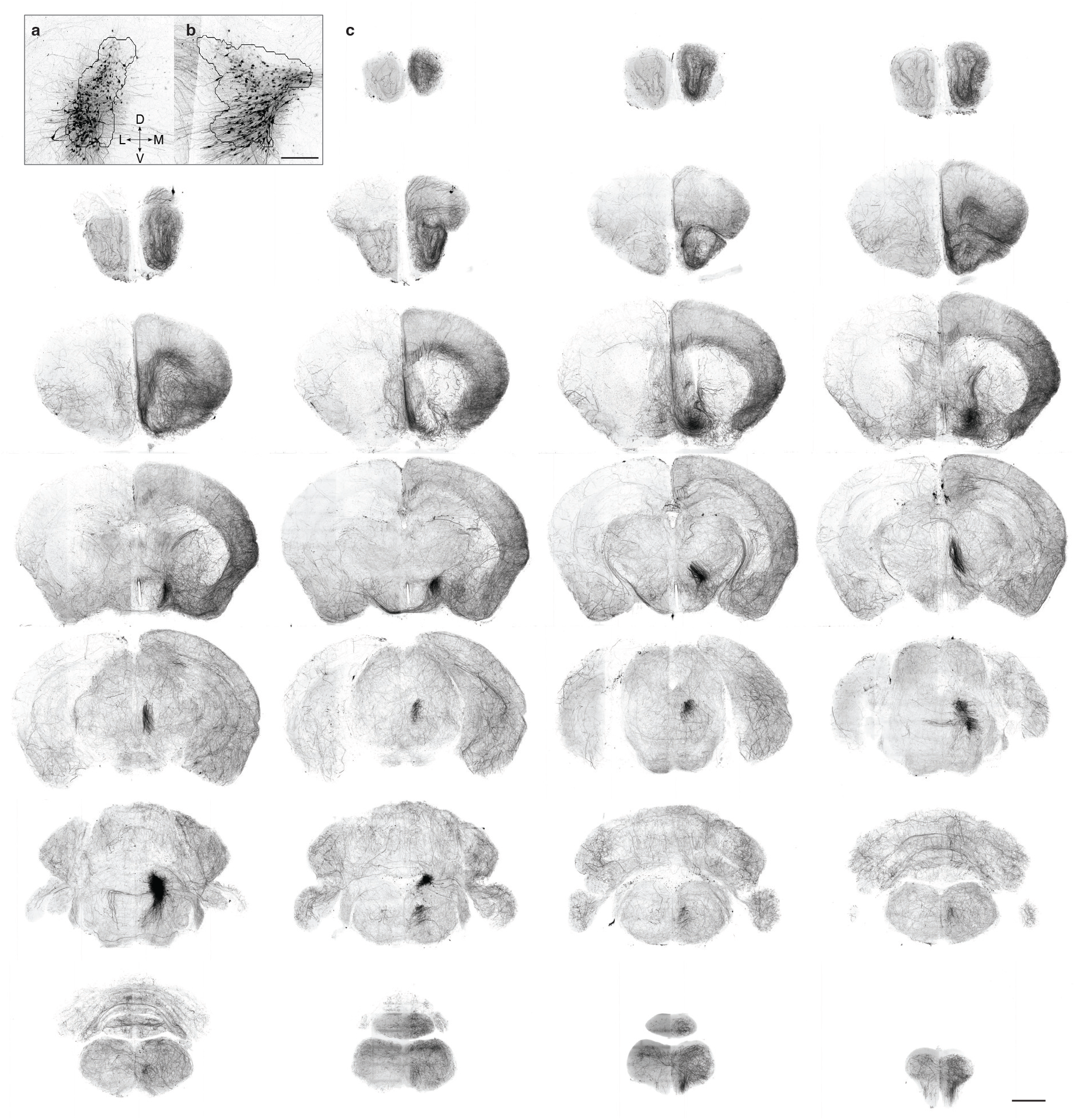
Brain-wide projections of LC-NE neurons in the right hemisphere. **a, b**, Coronal and sagittal views of dense unilateral labeling of LC-NE neurons visualized with a rendering of the specimen space “reverse-morphed” LC mesh. Scale bar, 2 mm. **c**, Montage of maximum intensity projections of serial 2 mm coronal slabs showing LC-NE innervation across the entire sample. Scale bar, 5 mm. See supplementary video for a full volume.

**Figure S3:**
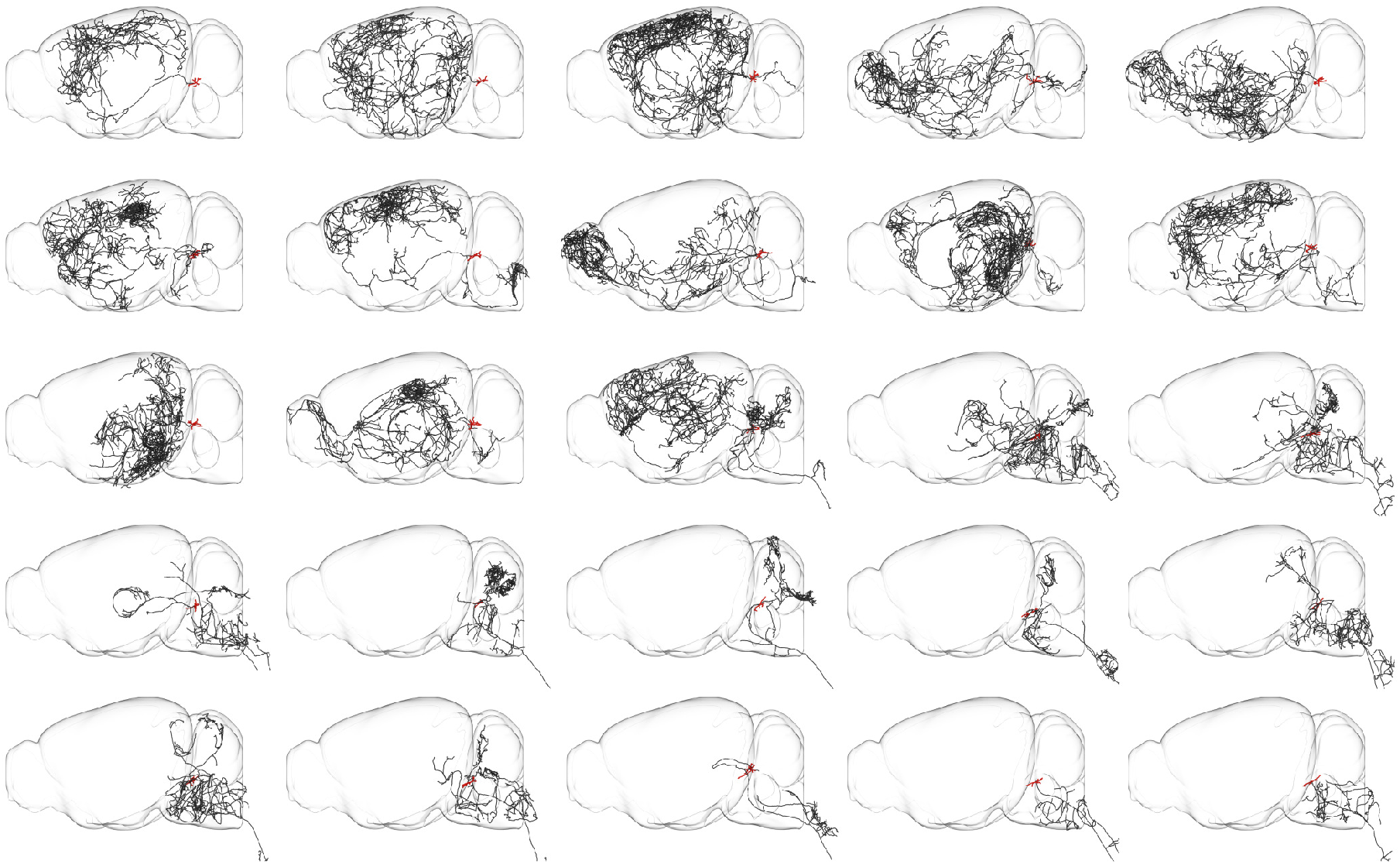
Individual LC-NE morphologies. Examples of reconstructions of LC-NE neurons, illustrating a range of primary projection targets across the brain (see Figure 2). Red, somata and dendrites. Black, axons. See https://morphology.allenneuraldynamics.org/.

**Figure S4:**
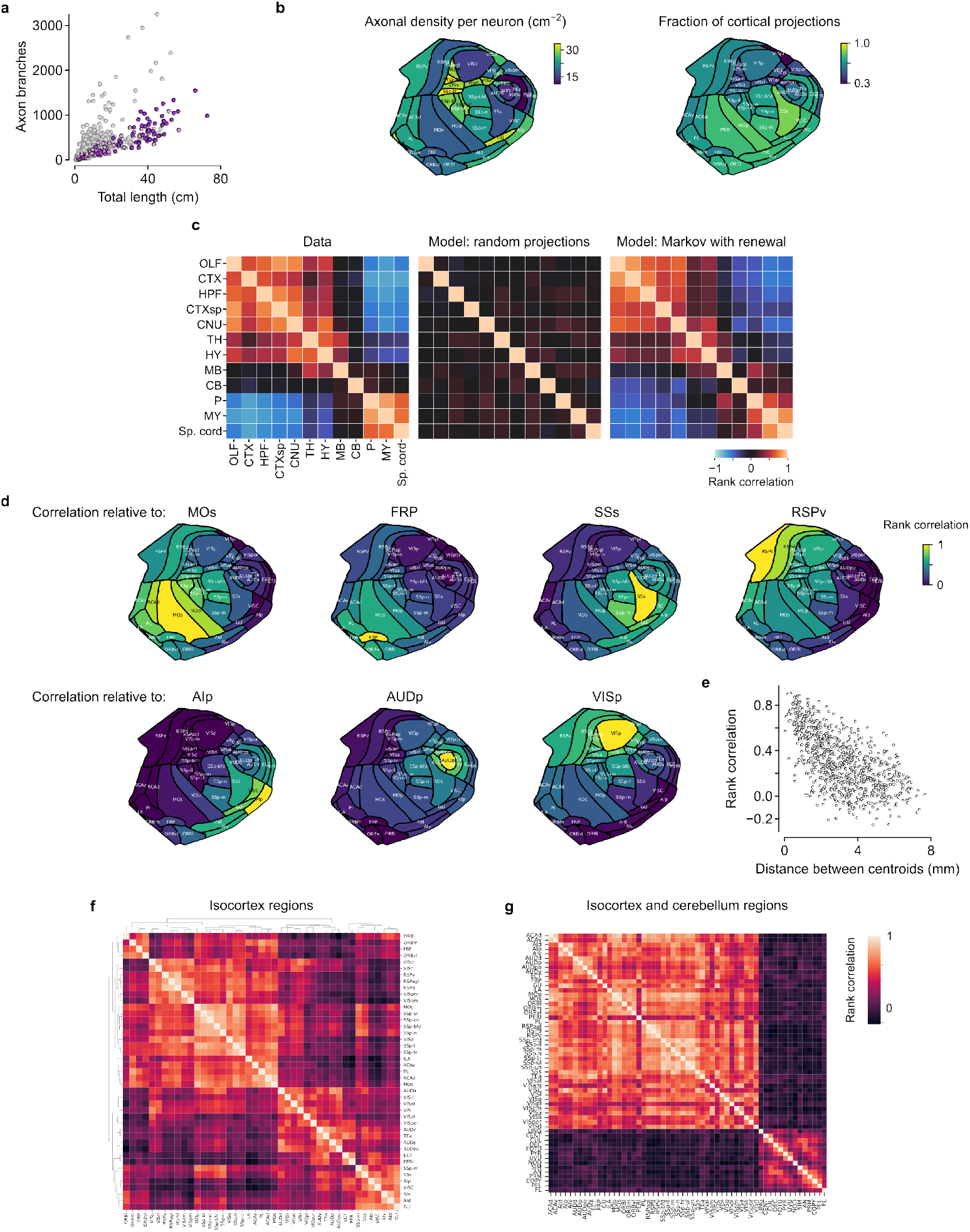
Spatial distribution of LC-NE axon innervation. **a**, Number of axon branches versus total axon length for LC-NE neurons traced to completion (dark purple) or cut off at the spinal cord (light purple), together with data from the MouseLight database (Winnubst et al. 2019). **b**, Axonal density and fraction per neuron innervating different cortical regions, represented as flatmaps. **c**, Comparison of correlations of single-neuron projections to different regions from the data (left, reproduced from Figure 2c) and models with different assumptions about axonal growth patterns. A random walk with simple assumptions is qualitatively similar to the data (with many quantitative differences). **d**, Correlation of axon length in seven cortical regions relative to others. Note high spatial correlations among adjacent structures that decreases with distance in cortex. **e**, Correlation between axon length innervation among pairs of cortical regions as a function of centroid distance between regions. **f**, Rank correlation of axonal length between cortical regions (defined at https://atlas.brain-map.org/). Regions are sorted by hierarchical clustering on the heatmap. **g**, Correlation between cortical and cerebellar regions, for neurons with either a cortical or cerebellar projection.

**Figure S5:**
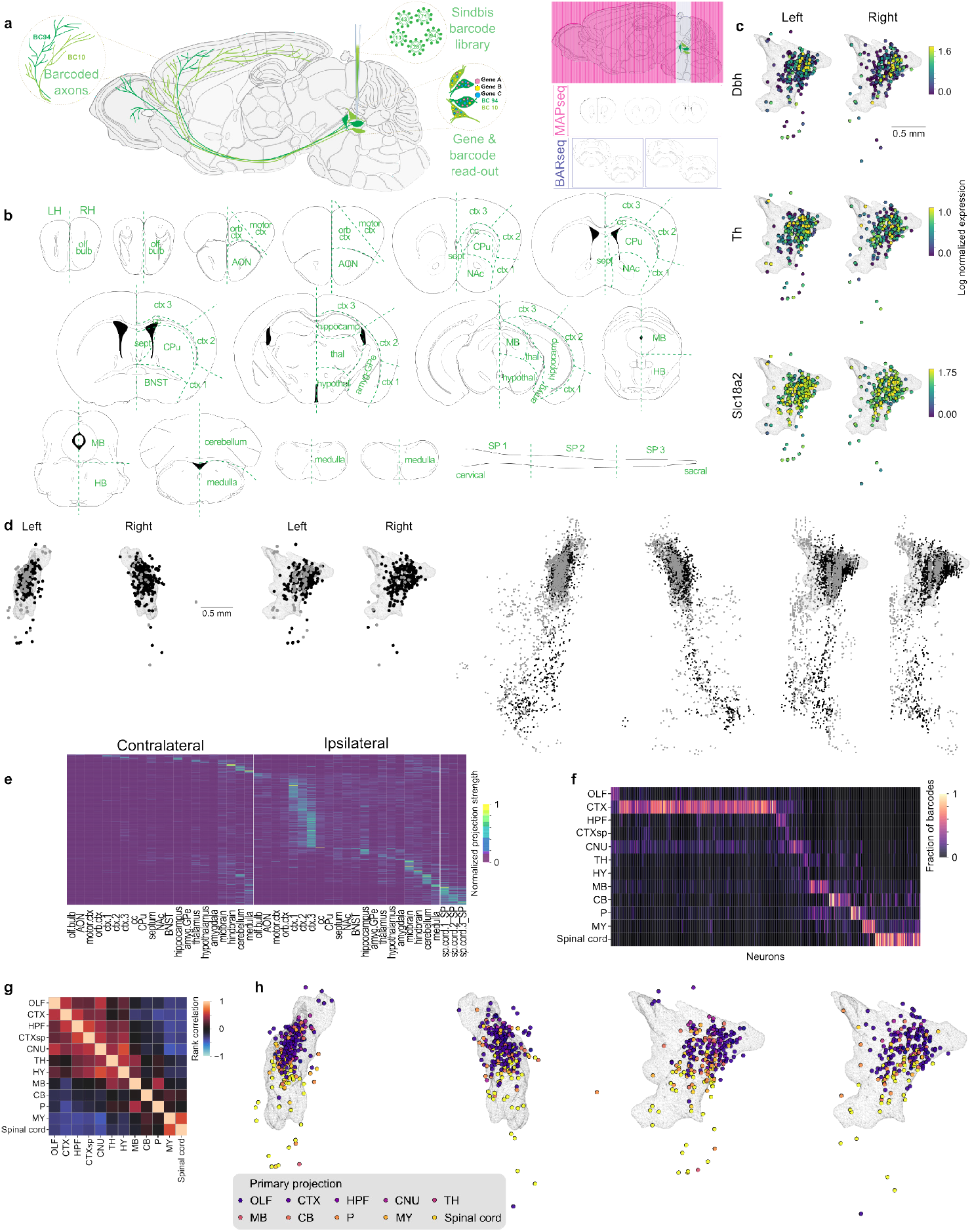
LC-NE projections measured with MAPseq and BARseq. **a**, Schema of experiment. **b**, Dissection boundaries for MAPseq. **c**, Expression of three LC-NE marker genes in cells analyzed in MAPseq/BARseq. **d**, Left: locations of LC-NE neurons analyzed in MAPseq/BARseq for each brain (black or gray). Right: locations of NE neurons present in each of the two samples (including those not labeled with barcodes). **e**, Normalized projection strengths ipsilateral and contralateral to each cell’s soma, for each dissected region. **f**, Fraction of barcodes detected across neurons for each CNS region of interest. Compare to Figure 2c. **g**, Rank correlation of shared projections. Compare to Figure 2c. **h**, Spatial distribution of LC-NE neurons with primary projections to ten regions. Compare to Figure 2e.

**Figure S6:**
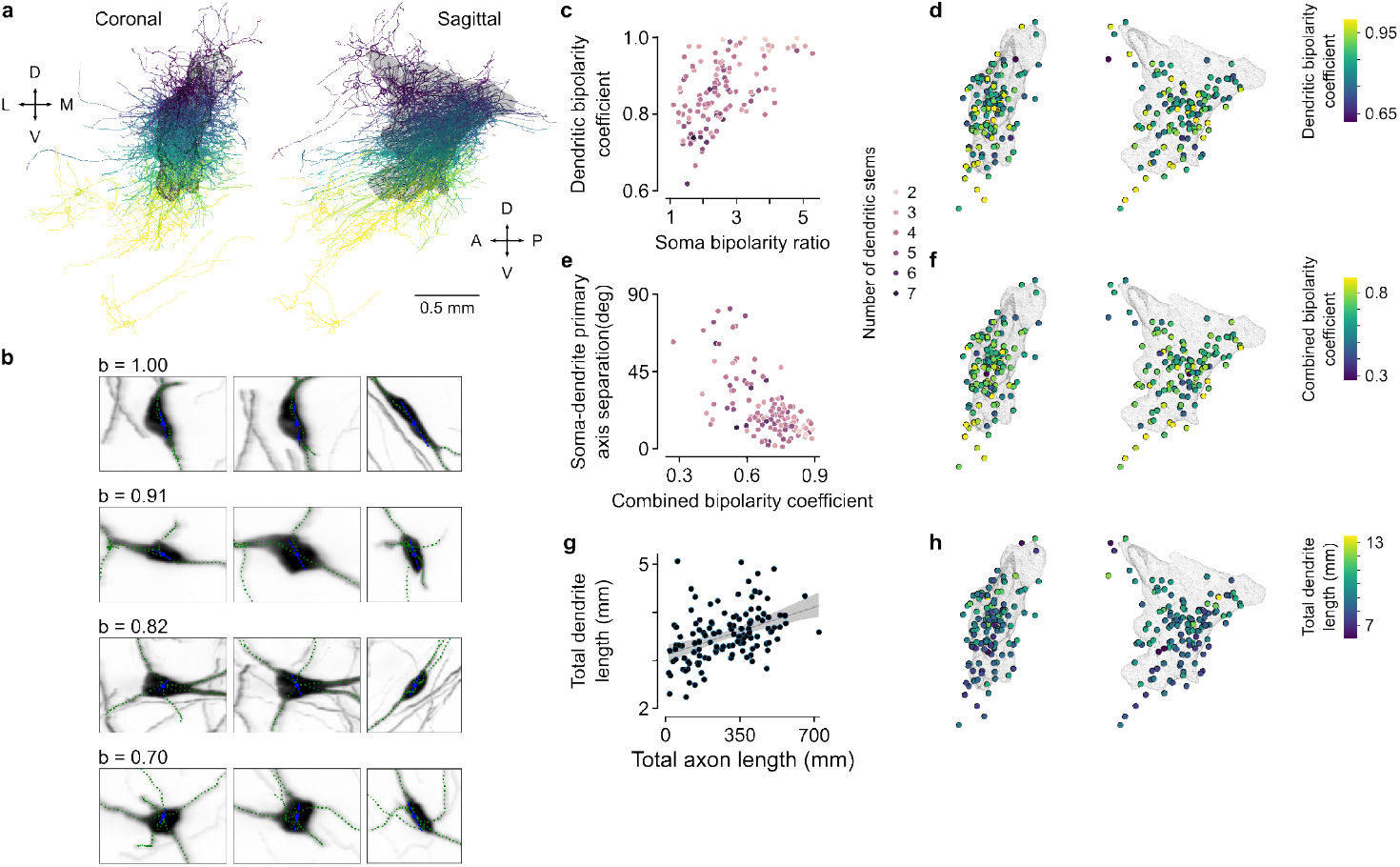
Somatodendritic morphologies of LC-NE neurons vary continuously. **a**, Coronal and sagittal projections of dendrites. Color corresponds to each cell’s location in the dorsal-ventral axis. **b**, Four example neurons sorted by dendritic bipolarity coefficent b. Soma center and primary axis are shown in blue, dendrites in green, overlaid on maximum intensity projections of the soma and proximal dendrites. Each set of images shows projections in the coronal, horizontal, and sagittal planes, respectively. **c**, Relationship between somatic and dendritic bipolarity metrics, showing a continuum from multipolar to fusiform phenotypes (Swanson 1976; McKinney et al. 2023). **d**, Spatial distribution of dendritic bipolarity coefficients in LC. **e**, Relationship between angular separation between primary axes of soma and dendrites and somatodendritic bipolarity (harmonic mean of two bipolarity coefficients). Fusiform cells (high bipolarity) have similar somatic and dendritic primary axes. **f**, Combined bipolarity in CCF space. **g**, Relationship between total length of dendrites and axons. **h**, Total dendritic length in CCF space.

**Figure S7:**
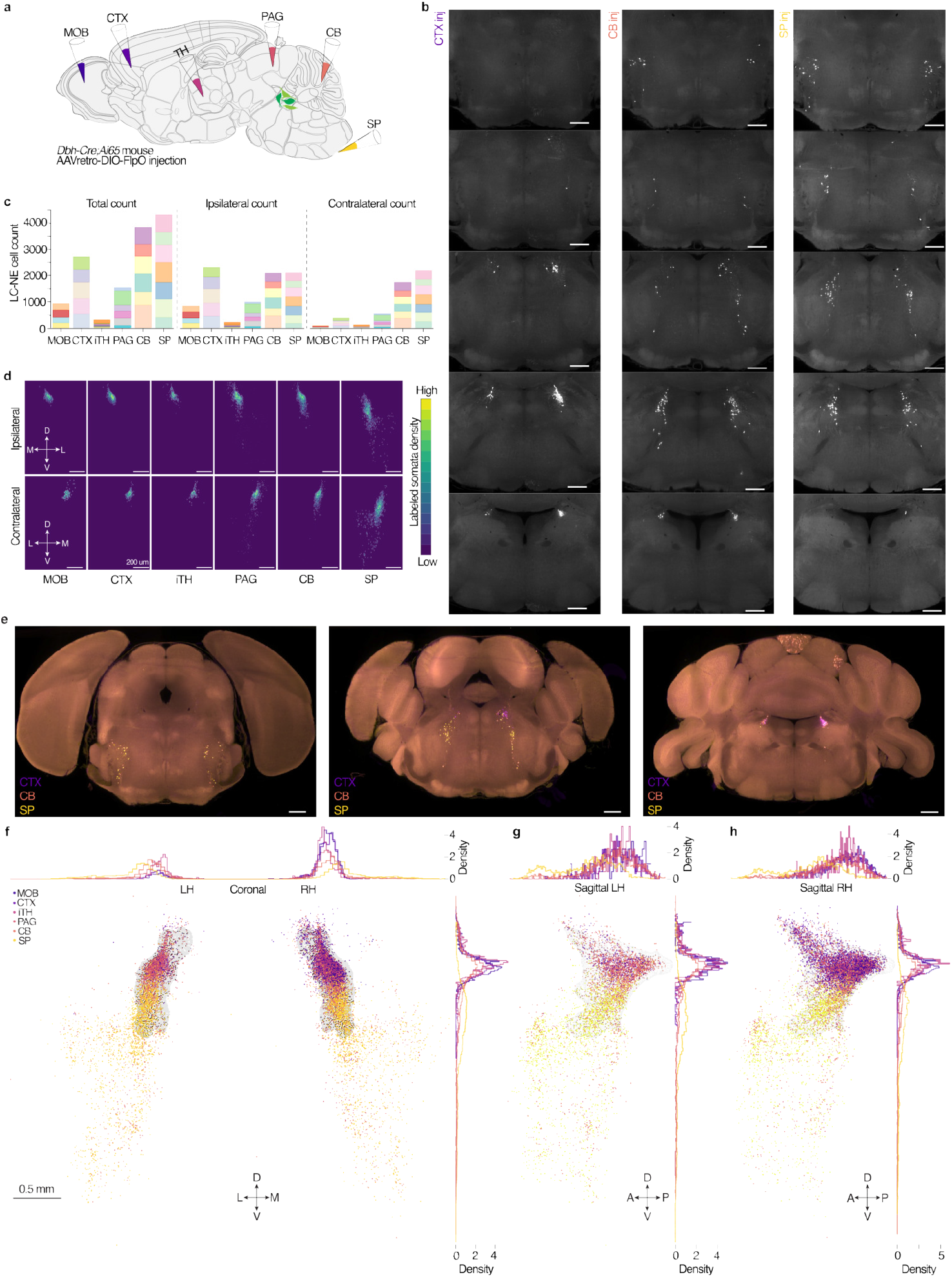
Retrograde tracing to quantify locations of somata with projections to olfactory bulb, frontal cortex, thalamus, periaqueductal gray, cerebellar cortex, or spinal cord. **a**, Schematic of injection sites for retrograde tracing. **b**, Examples of CCF-registered images at five coronal planes for three injection sites. Note labeling in LC. **c**, Counts of retrogradely-labeled LC-NE neurons for each mouse (colors) and region. **d**, Density of retrogradely-labeled cells in the coronal planes, by projection target. **e**, Overlay of retrogradely-labeled neurons from each target in the coronal plane. **f**, Retrogradely-labeled neurons in the coronal plane. LH, left hemisphere. RH, right hemisphere. **g** and **h**, Sagittal planes for the data in **f**.

**Figure S8:**
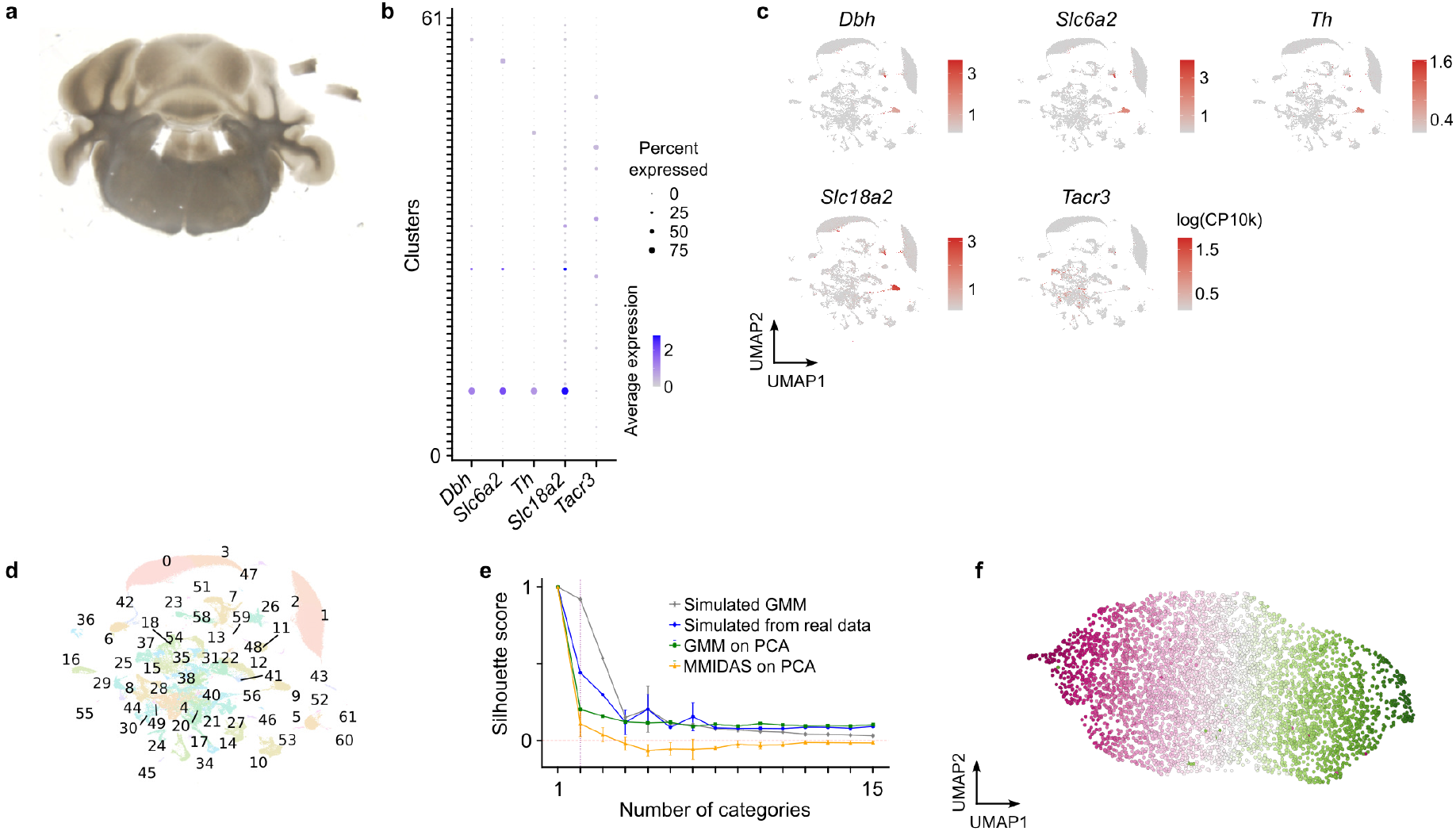
LC-NE neurons have distinct gene expression from surrounding cells. **a**, Example dissection of the region around LC for snRNAseq analysis. **b**, UMAP of gene expression clusters across all cells from the dissection (398,912 total cells, 234,813 total cells that passed quality control, 4,845 LC-NE neurons). **c**, Markers for LC-NE neurons occupy a distinct cluster that we define as LC-NE neurons: *Dbh*, dopamine beta hydroxylase. *Th*, tyrosine hydroxylase. *Slc6a2*, norepinephrine transporter. *Slc18a2*, vesicular monoamine transporter 2. **d**, Clusters defined from UMAP applied to all cells. **e**, MMIDAS model indicates a single cluster best explains the data. **f**, UMAP representation of the data. Color scale as in Figure 3.

**Figure S9:**
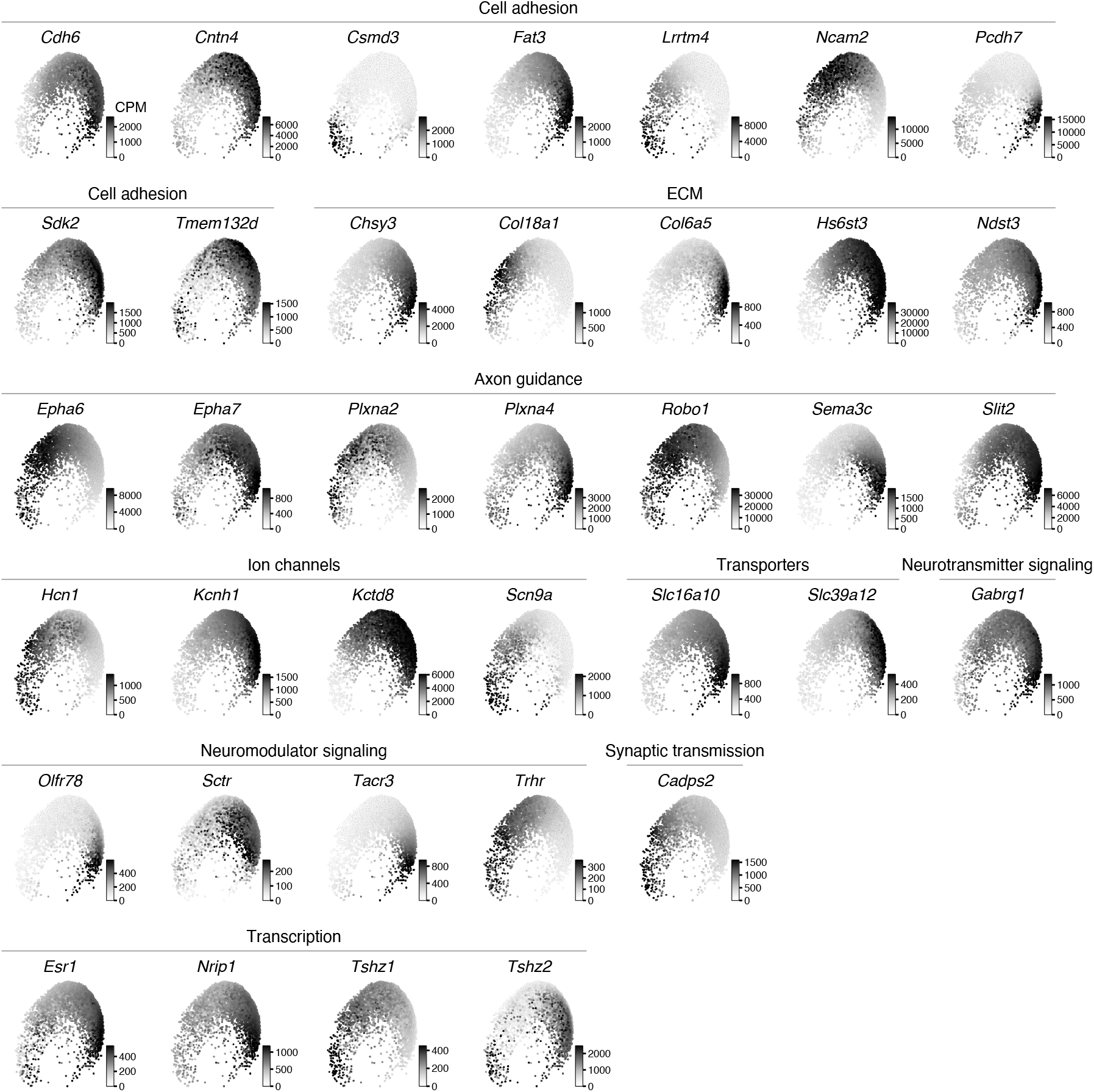
Expression levels of genes in different categories of putative functions. Examples of graded expression of genes related to cell adhesion, extracellular matrix, axon guidance, ion channels, transporters, neurotransmitter and neuromodulator signaling, synaptic transmission, and transcription.

**Figure S10:**
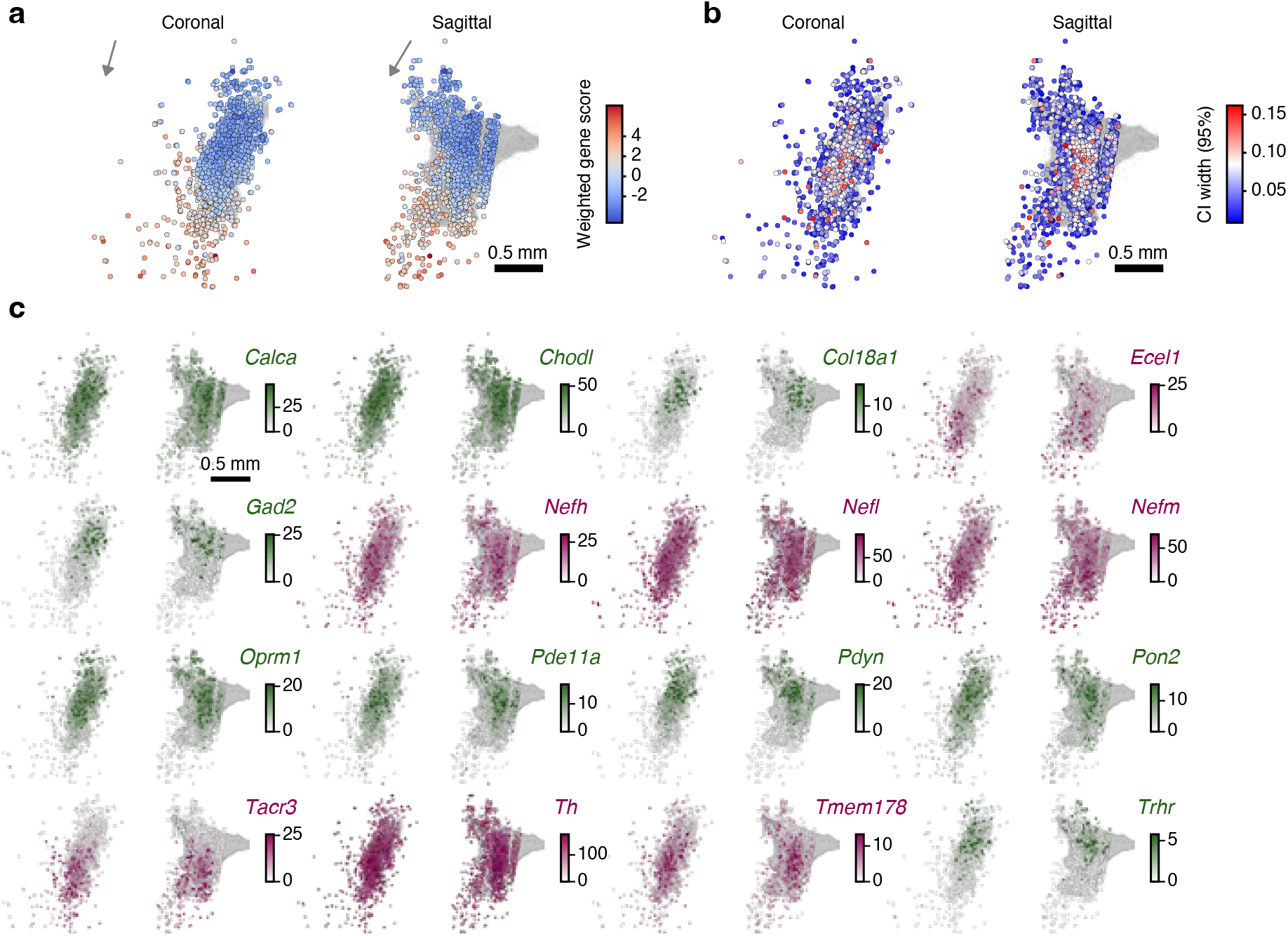
MERFISH analysis. **a**, CCA analyses of population gene expression. **b**, CI around CCA analysis of population gene expression. **c**, Example genes with spatial gradients from MERFISH data.

**Figure S11:**
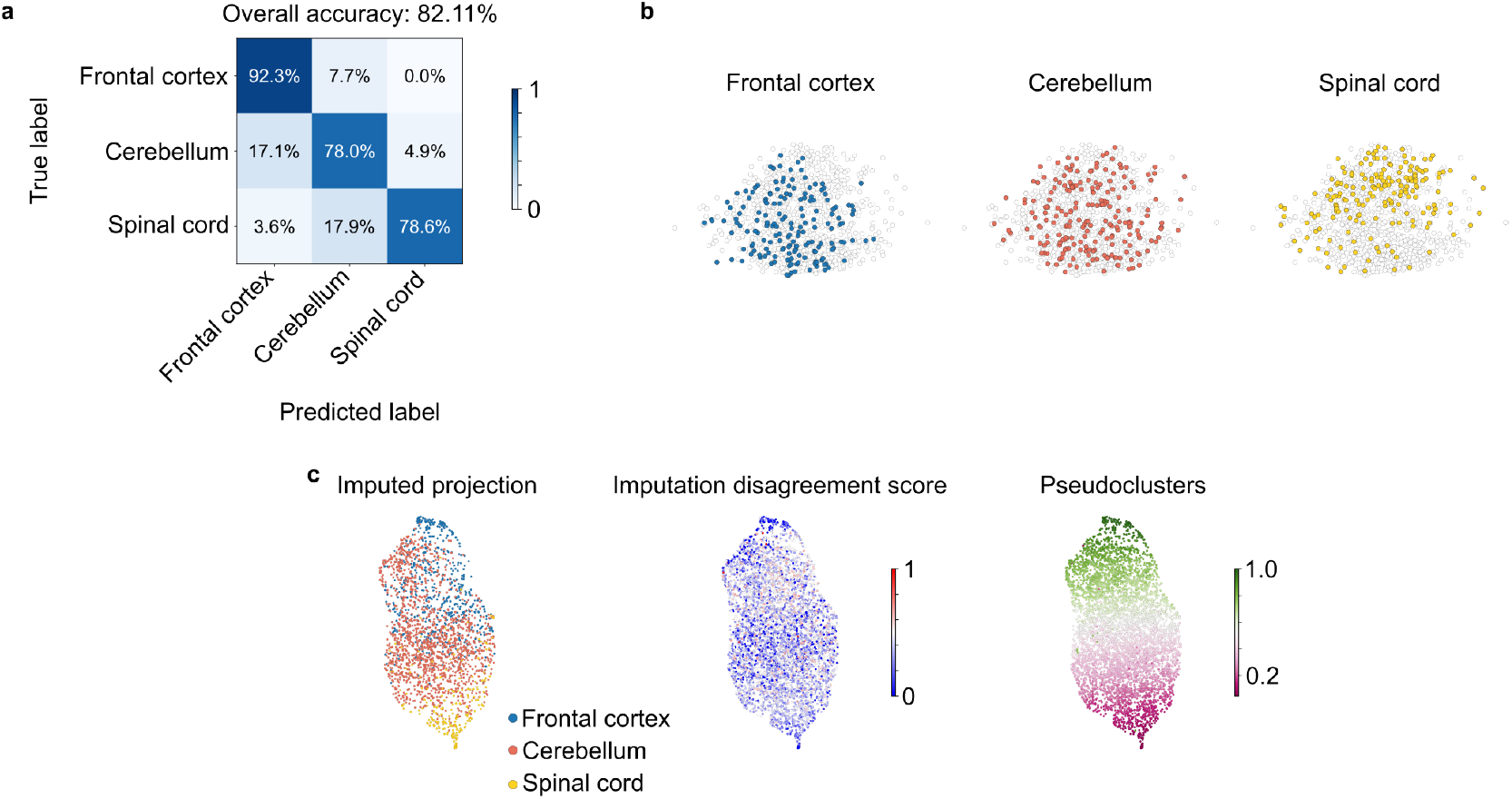
Retro-seq. **a**, GradientBoosting classifiers were trained on the retro-seq dataset (raw counts) to predict projection targets. **b**, Retro-seq cells embedded by PCA using the scVI normalized counts. Projection target labels were restricted to frontal cortex, cerebellum, and spinal cord. **c**, Right: projection targets were imputed onto snRNA-seq cells by k-nearest neighbor matching (k = 10). Cells were assigned by majority vote among labeled neighbors with ≥80% agreement; otherwise left unassigned. Left, inferred projection target; middle, imputation disagreement score (1 – nmax/nvalid,), where (nmax) is the count of the most frequent neighbor label and (nvalid) is the number of labelled neighbors (range 0–1). Only the first two subpanels of the three-panel display are shown.

**Figure S12:**
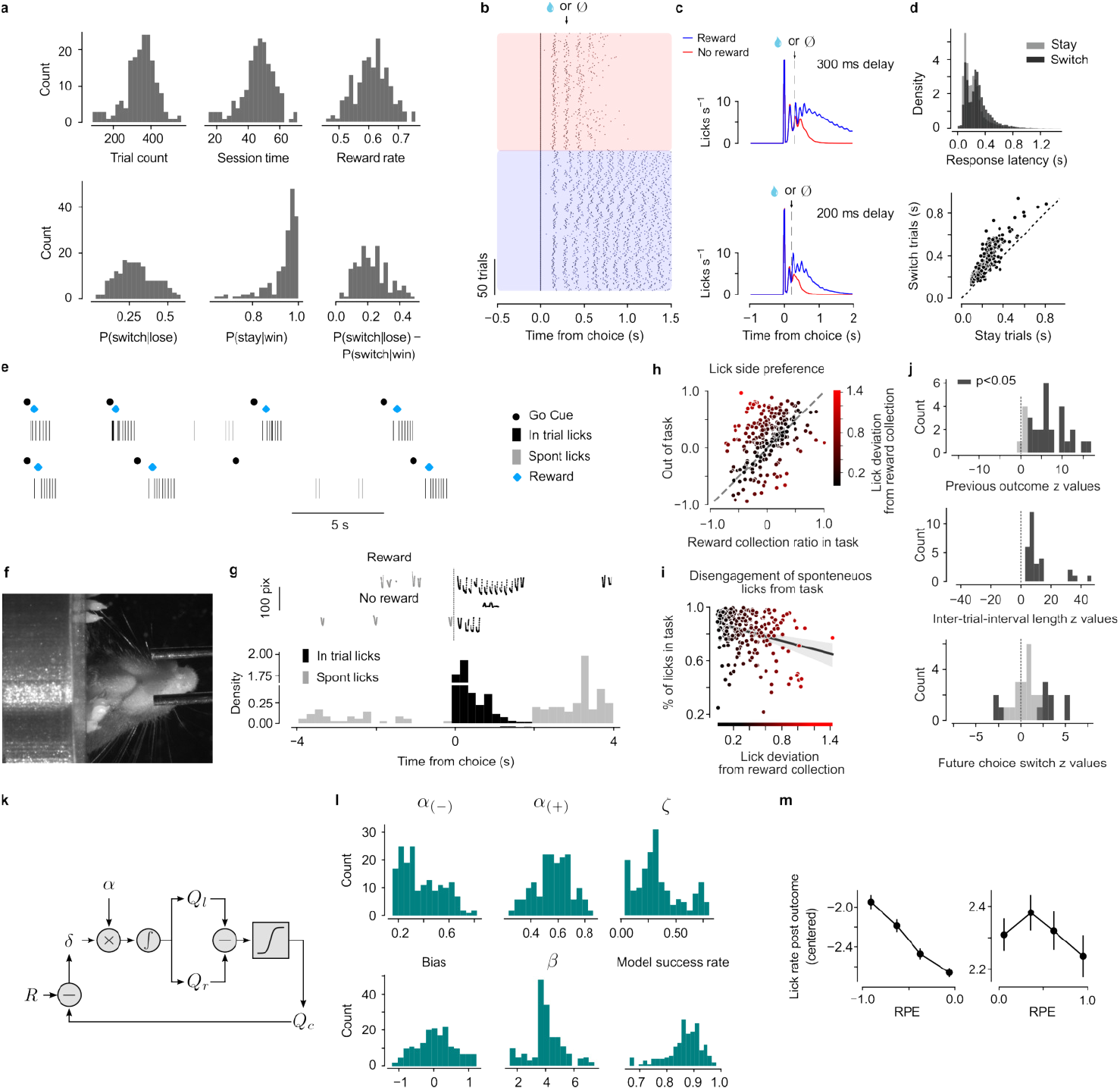
Statistics of behavior. **a**, Distributions of behavioral performance statistics. **b**, Example lick raster aligned to start of choice. Blue: rewarded trials. Red: unrewarded trials. **c**, Example mean licking rate of sessions with reward latency of 200 ms and 300 ms. **d**, Top: distribution of lick latencies of stay and switch trials. Bottom: lick latency of switch vs. stay choices. Each dot is one session. **e**, Example in-trial licks (black) and spontaneous licks (gray). **f**, Example video frame. **g**, Licks detected by video aligned to go cue. Top: example rewarded and unrewarded trials. Bottom: example session’s lick time distribution. **h**, Lick preference (left/right) in spontaneous period vs. normalized reward ratios (left/right) in the task (Pearson’s *r* = 0.39, *p* = 1.54 × 10^−10^). **i**, % of licks in task vs. deviation of lick preference from reward ratio shown in (**h**; Spearman’s *r* = –0.26, *p* = 3.40 × 10^−5^). **j**, Distribution of *z*-statistics of logistic regression models where previous outcomes, length of ITI, and further choice switches are used to predict if a spontaneous lick happens. **k**, Reinforcement model schema. **l**, Distribution of fitted model parameters. One data point is one mouse. **m**, Lick rate changes with negative RPE.

**Figure S13:**
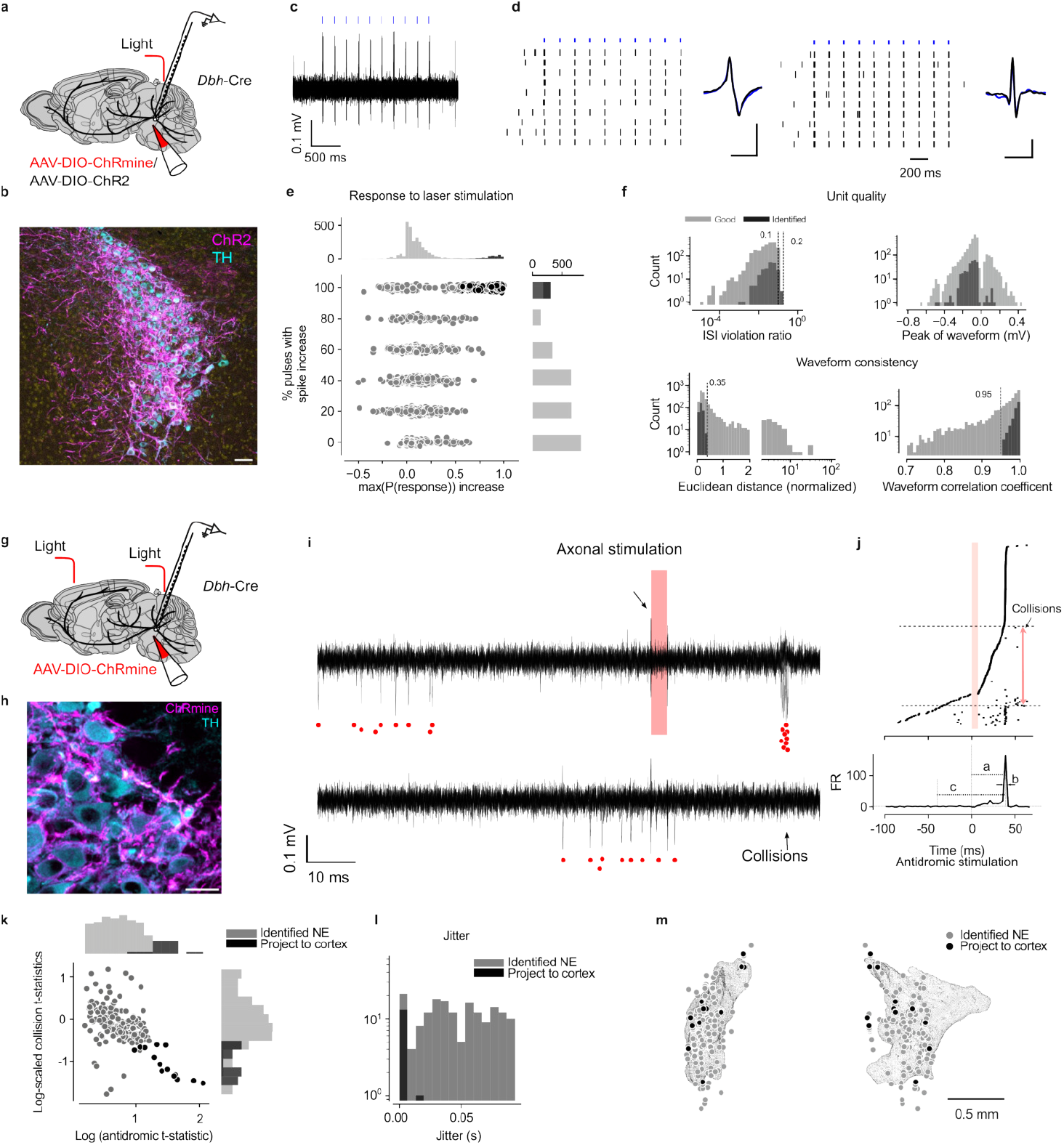
Identifying LC-NE neurons *in vivo*. **a**, ChRmine, Chrimson, or ChR2 were injected into LC of *Dbh*-Cre animals for optical identification with Neuropixels probes or tetrodes. **b**, Expression of ChR2 in LC-NE neurons. Scale bar: 50 µm. **c**, Example raw trace of a tetrode recording with spikes following laser stimulation (blue ticks). **d**, Two example identified LC-NE neurons in response to multiple trains of light stimuli. **e**, Quality metrics of LC-NE neurons. Gray: single units. Black: identified as NE neurons. Top row: basic quality metrics of neurons; bottom row: waveform similarity between stimulated and spontaneous spikes. **f**, Distribution of laser-evoked response properties for identified NE neurons and unidentified neurons. **g**, Schema for antidromic stimulation: ChRmine was injected into LC of *Dbh*-Cre animals. To perform antidromic stimulation, light was delivered to the brain surface through the skull over frontal cortex. **h**, Expression of ChRmine in LC-NE neurons. Scale bar: 20 µm. **i**, Example antidromic stimulation. Top row: antidromic spikes; Bottom row: collision when a spontaneous spike preceded the antidromic spike. **j**, Example antidromic stimulation across trials. Top: Rows are sorted by the time of first spike. Bottom: mean spike rate aligned to the time of antidromic stimulation. a is antidromic latency, defined as the time of peak mean spike rate aligned to time of laser stimulation; b is jitter, defined as the half peak width of mean spike rate aligned to laser stimulation; c is antidromic window, during which if a spontaneous spike occurred, a collision occurred. **k**, Distribution of antidromic response properties of all identified LC-NE neurons, identified to project to cortex (black) or not (gray). *t*-statistics are log-scaled for ease of visualization. **l**, Distribution of light-evoked response jitter of LC-NE cells, identified to project to cortex versus not. **m**, Distribution of all identified NE neurons in space, highlighting cortex-projecting NE neurons.

**Figure S14:**
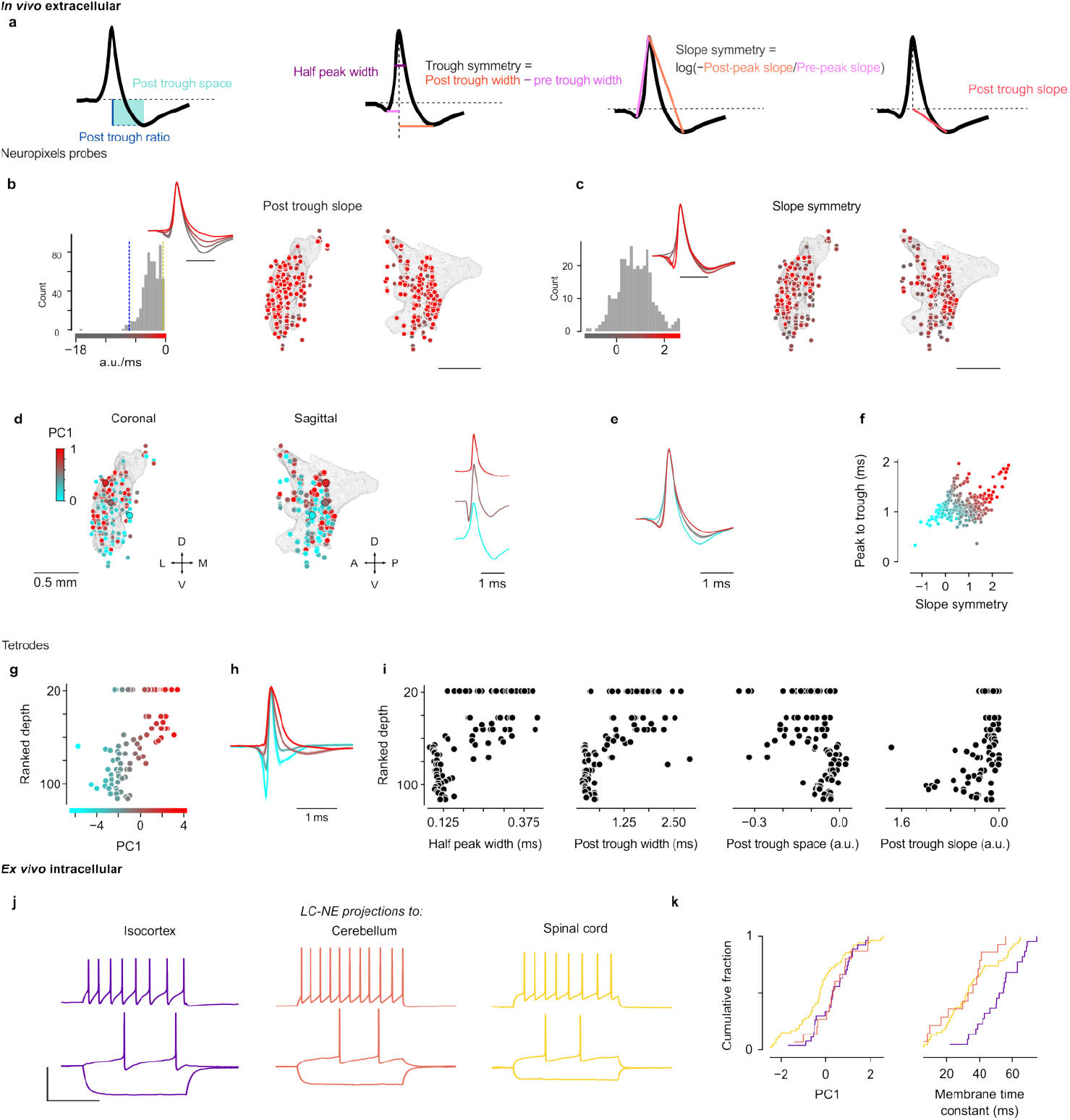
LC-NE spike waveforms are spatially organized. **a**, Illustration of waveform features. **b**, Distribution of post trough slope in space (kNN cross-validated *p* = 0.59; linear model, *p* = 4.0 × 10^−3^). **c**, Distribution of slope symmetry in space (kNN cross-validated *p* = 5.0 × 10^−4^; linear model *p* = 5.0 × 10^−4^). **d**, The first PC (PC1) of features extracted from extracellular action potential waveforms were distributed in space. **e**, Mean normalized waveforms from five quintiles of PC1. **f**, Two features of the waveform co-varied: the symmetry of the rising and falling slopes and the time from peak to trough. **g**, Ranked depth of spike waveforms from tetrode recordings as a function of PC1. **h**, Quantiles of mean spike waveforms from tetrode recordings. **i**, Ranked depth versus spike waveform features. **j**, Examples of *ex vivo* whole-cell patch clamp recordings from LC-NE neurons with projections to either isocortex, cerebellum, or spinal cord. Top row: current injected at rheobase plus 30 pA. Bottom row: depolarizing current injected at rheobase. Scale bars: 50 mV, 500 ms. **k**, Intracellular action potential waveforms (left; paired *t*-tests, spinal cord vs. cortex, *p* < 0.01, spinal cord vs. cerebellum, *p* < 0.05) and membrane time constants (right; paired *t*-tests, spinal cord vs. cortex, *p* < 0.001, cerebellum vs. cortex, *p* < 0.001) showed differences by projection target.

**Figure S15:**
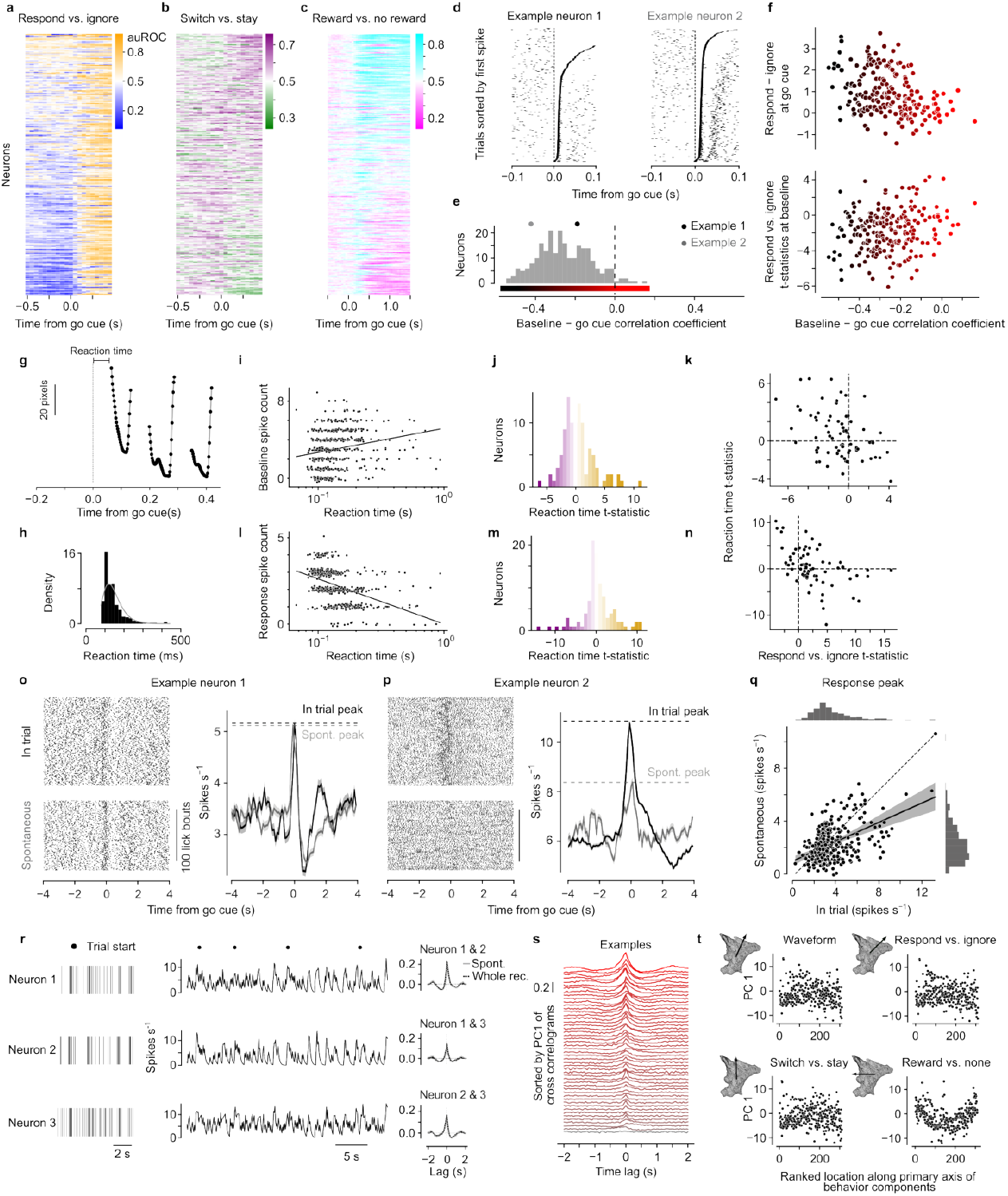
Firing properties of LC-NE neurons during behavior. Area under the receiver operating characteristic (auROC) curve of response vs. ignore (**a**), switch vs. stay (**b**), and reward vs. no reward (**c**). **d**, Example neurons, aligned to time of go cue, with trials sorted by time of first spike after go cue. **e**, Distribution of correlation between baseline activity (0.5 s before go cue) and activity 0.3 s after go cue. Values for examples in (**d**) are marked as black and gray dots. **f**, Top, increase of spike rate in trials with lick responses vs. those without against tonic modulation strength from **e** (*r* = –0.59, *p* = 4.63 × 10^−21^). Bottom, correlation between *t*-statistics of linear regression of baseline spike rate as a function of respond vs. ignore on the next trial against tonic modulation strength from **e** (*r* = –0.59, *p* = 4.63 × 10^−21^). **g**, Example tongue position relative to go cue. Reaction time is defined as the time from the go cue until the first video frame in which the tongue was detected. **h**, Distribution of reaction times for an example session (lognormal fit in gray). **i**, Example neuron’s spike count in the baseline window (1 s before go cues) vs. reaction time. Trend line shows linear fit on log scale for reaction time. Points are jittered for display. **j**, Distribution of *t*-statistics predicting reaction time in the baseline window (104 neurons, 15 mice). **k**, *t*-statistics predicting respond vs. ignore trials (see Figure 5c) against those predicting reaction time in both the baseline window. **l**, Example neuron from (**i**) in the response window. **m**, Distribution of *t*-statistics in the response window as in (**j**). **n**, Respond vs. ignore *t*-statistics against those predicting reaction time in the response window. **o, p**, Two example neurons’ activity aligned to lick bout starts, for licks during the task (“in trial”) and for spontaneous licks. **q**, Peak spike rate around spontaneous vs. in-trial licks (*r* = 0.49, *p* < 0.001). **r**, Three simultaneously recorded neurons with correlated spike rates. Right: cross-correlations across the whole experiment and during spontaneous spiking are similar. **s**, Example cross-correlograms across pairs of neurons (gray to red indicates low to high). **t**, Spatial analysis shows PC1 of cross-correlogram changes along axes of outcome, but not other task responses (kNN cross-validation bootstrapped *p*-values: waveform: 0.1529, respond vs. ignore: 0.7166, switch vs. stay: 0.5787, reward vs. no reward: 0.0005).

**Figure S16:**
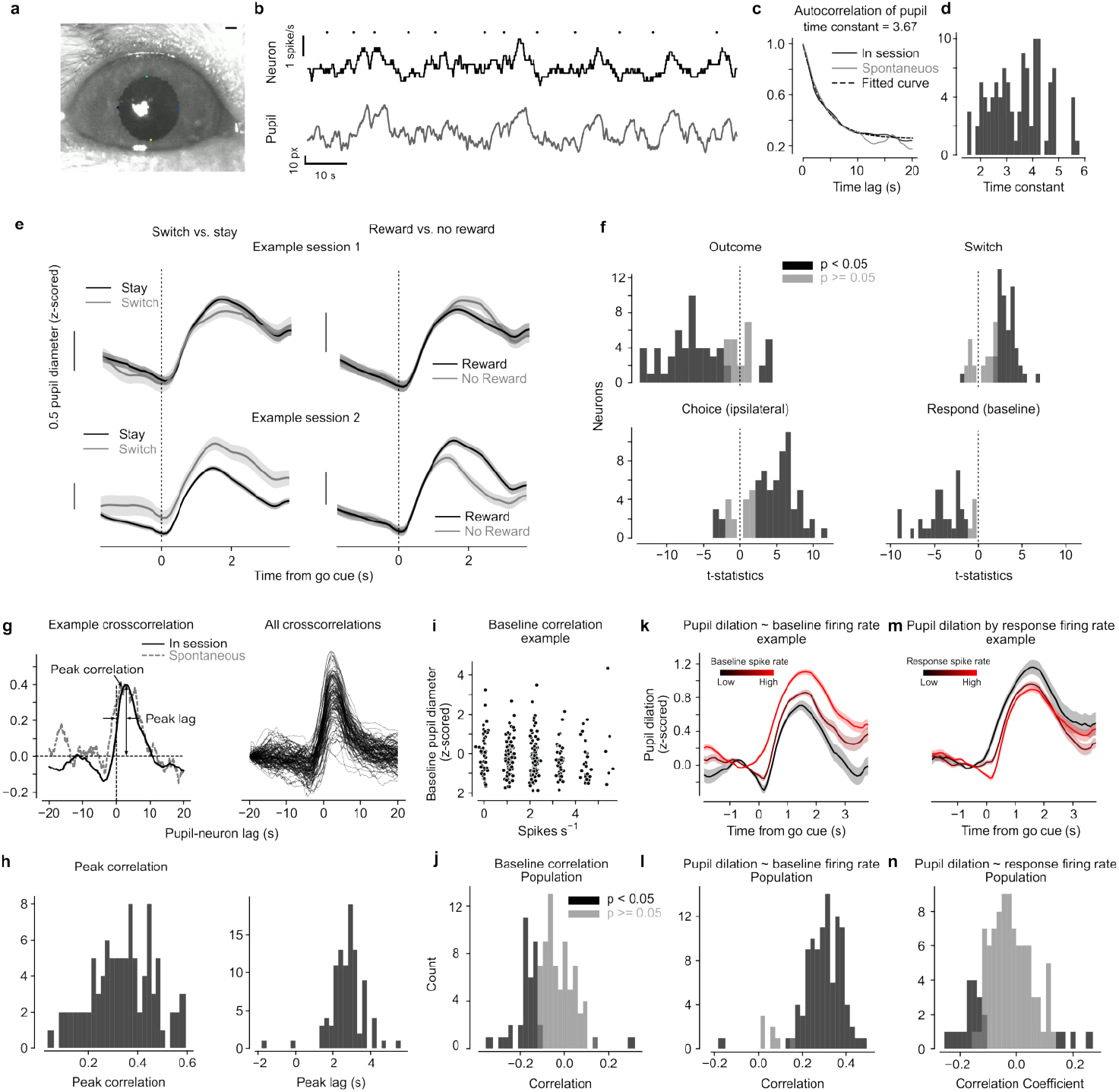
LC-NE neuron spiking correlates with pupil size. **a**, Example video screenshot. Scale bar is 20 pixels. **b**, Example simultaneously recorded neuron activity and pupil diameter. Go cues are marked as black dots. **c**, Autocorrelation of pupil diameter and fitted exponential decay curve (τ = 3.67) for one example neuron. Black and gray curves are for autocorrelations computed in whole session and baseline period. Distribution of τ across all sessions’ pupil recording fitted for whole session autocorrelation shown in **d**,. **e**, Two example sessions with mean spike rate aligned to start of a trial, separated by if a mouse got rewarded and whether they switched choice. **f**, Distribution of t-statistics for regressions on pupil diameter with different task components in different periods of time: top left: reward or not in 1.5 s after go cue; top right: switch or not in 1.5 second after go cue; bottom left: choice side (ipsilateral is 1 and contralateral is 0) in 1.5 s after go cue. **g**, Left: example cross-correlation between simultaneously recorded LC-NE neuron activity and pupil diameter. Solid black line: whole session. Gray dashed line: spontaneous period outside of the task. Peak lag is defined at the time of peak correlation. Positive values indicate pupil dilation after spike rate increase. Right: overlayed cross-correlation across neurons. **h**, Distribution of peak values and peak lag defined in (**g**). **i**, Example baseline pupil diameter vs. baseline spike rate (linear correlation, *r* = 0.46, *p* < 0.001). **j**, Distribution of correlation coefficient between baseline pupil diameter and baseline firing rate across LC-NE neurons. **k**, Example pupil dilation aligned to trial start, binned by baseline spike rate of a LC-NE neuron. **l**, Distribution of correlation coefficient between peak pupil dilation and baseline spike rate across LC NE neurons. **m**, Example pupil dilation aligned to trial start, binned by response to go cue of a NE neuron (linear correlation, *r* = 0.17, *p* < 0.01). **n**, Distribution of correlation coefficients between peak pupil dilation and response to go cue across LC-NE neurons. Black, *p* < 0.05. Gray, *p* ≥ 0.05.

**Figure S17:**
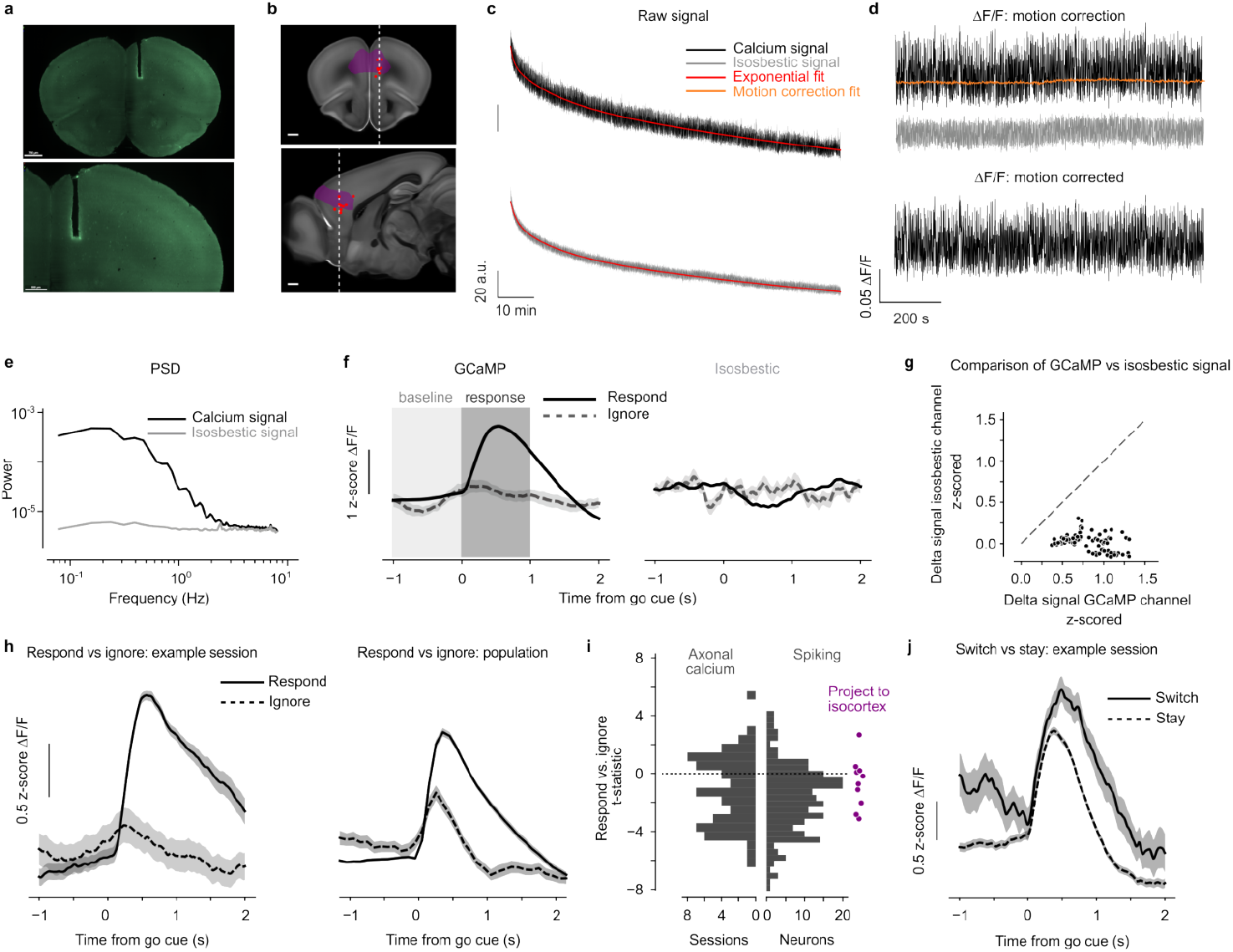
LC-NE axonal calcium activity in frontal cortex. **a**, Example reconstruction of an optic fiber implanted in PL. **b**, Optic fiber termini across mice. **c**, Example raw data from one experiment, illustrating data processing steps. **d**, Example motion-corrected traces of ΔF/F. **e**, Power spectra of GCaMP and isosbestic channels from one experiment. **f**, Example experiment showing GCaMP and isosbestic signals around go cues during trials with or without lick responses. **g**, Change in isosbestic vs. GCaMP signals after the go cue across experiments, defined as change of mean signal in the response window relative to the baseline window. **h**, Example and population data showing larger activity during trials and smaller activity before trials with lick responses. **i**, Comparison of activity after during baseline on trials with versus without lick responses. Right histogram shows single-neuron electrophysiological data. Axonal fluorescence vs. single-neuron spiking, Fisher’s exact test with permutation, *p* = 4.2 × 10^−4^. Single neurons projecting to cortex compared to the population of single neurons, Welch test with permutation, *p* = 6.7 × 10^−2^; Fisher’s exact test with permutation, *p* = 8.8 × 10^−2^. **j**, Example activity during switch versus stay trials.

**Figure S18:**
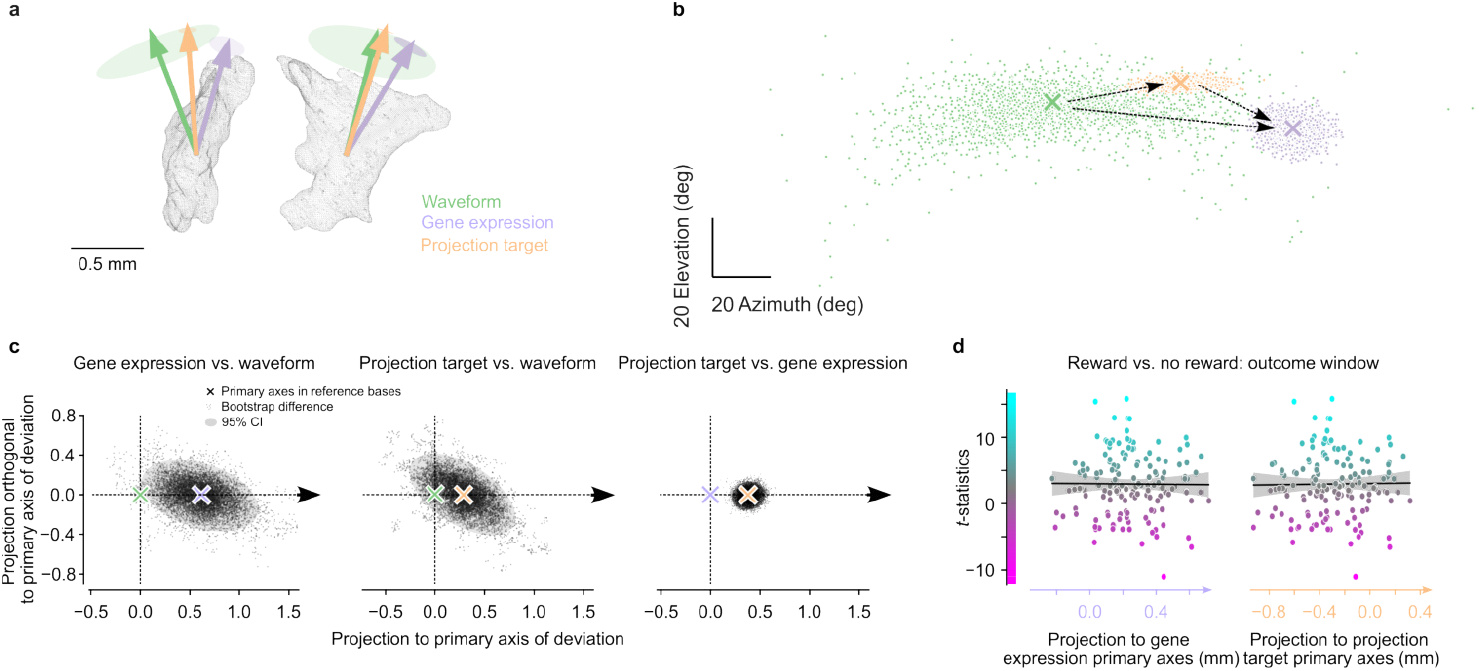
Spatial comparisons between gene expression, retrograde tracing data, and extracellular spike waveforms. **a**, Primary axis of gene expression, projection target and spike waveform features. Shaded ellipse represents bootstrapped 95% CI. **b**, Distribution of bootstrapped primary axes represented as azimuth (horizontal plane) and elevation (along the dorsal-ventral axis). **c**, Comparisons between primary axes across cellular properties. Differences of bootstrapped samples are projected to reference bases defined by primary axes of deviation of two primary axes. Shaded regions represent 95% CI. Bootstrapped values, left to right: 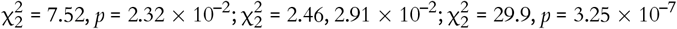. **d**, *t*-statistics of reward vs. no reward (Figure 6) plotted against each neuron’s location along the primary axes from different modalities. From left to right: *r* = –0.01, *p* = 0.91; *r* = 0.01, *p* = 0.86).

